# scDrugPrio: A framework for the analysis of single-cell transcriptomics to address multiple problems in precision medicine in immune-mediated inflammatory diseases

**DOI:** 10.1101/2023.11.08.566249

**Authors:** Samuel Schäfer, Martin Smelik, Oleg Sysoev, Yelin Zhao, Desiré Eklund, Sandra Lilja, Mika Gustafsson, Holger Heyn, Antonio Julia, István A. Kovács, Joseph Loscalzo, Sara Marsal, Huan Zhang, Xinxiu Li, Danuta Gawel, Hui Wang, Mikael Benson

## Abstract

**Background:** Ineffective drug treatment is a major problem for many patients with immune-mediated inflammatory diseases (IMIDs). Important reasons are the lack of systematic solutions for drug prioritisation and repurposing based on characterisation of the complex and heterogeneous cellular and molecular changes in IMIDs.

**Methods:** Here, we propose a computational framework, scDrugPrio, which constructs network models of inflammatory disease based on single-cell RNA sequencing (scRNA-seq) data. scDrugPrio constructs detailed network models of inflammatory diseases that integrate information on cell type-specific expression changes, altered cellular crosstalk and pharmacological properties for the selection and ranking of thousands of drugs.

**Results:** scDrugPrio was developed using a mouse model of antigen-induced arthritis and validated by improved precision/recall for approved drugs, as well as extensive *in vitro, in vivo,* and *in silico* studies of drugs that were predicted, but not approved, for the studied diseases. Next, scDrugPrio was applied to multiple sclerosis, Crohn’s disease, and psoriatic arthritis, further supporting scDrugPrio through prioritisation of relevant and approved drugs. However, in contrast to the mouse model of arthritis, great interindividual cellular and gene expression differences were found in patients with the same diagnosis. Such differences could explain why some patients did or did not respond to treatment. This explanation was supported by the application of scDrugPrio to scRNA-seq data from eleven individual Crohn’s disease patients. The analysis showed great variations in drug predictions between patients, for example, assigning a high rank to anti-TNF treatment in a responder and a low rank in a nonresponder to that treatment.

**Conclusion:** We propose a computational framework, scDrugPrio, for drug prioritisation based on scRNA-seq of IMID disease. Application to individual patients indicates scDrugPrio’s potential for personalised network-based drug screening on cellulome-, genome-, and drugome-wide scales. For this purpose, we made scDrugPrio into an easy-to-use R package (https://github.com/SDTC-CPMed/scDrugPrio).

## Introduction

Immune-mediated inflammatory diseases (IMIDs), such as rheumatoid arthritis, Crohn’s disease, and psoriatic arthritis, affect millions of people worldwide and can cause chronic pain, disability, and reduced quality of life (1). While new classes of therapies are transforming the management of IMIDs, it is still a general problem that many patients do not achieve remission with mono-(2, 3) or combinatorial therapy (3). This may be due to drug development involving testing drugs on large groups of patients, with the assumption that the drug will work similarly on all patients. Such an approach does not take into account the fact that each individual’s genetic makeup and environment are unique, leading to significant variations in drug efficacy and side effects.

Given that IMIDs are known to involve thousands of genes that are variably expressed in different cell types and show temporal and interindividual differences (4, 5), single-cell RNA sequencing (scRNA-seq) provides a promising foundation for the identification of suitable drug treatments (6). Indeed, one pioneering case report described scRNA-guided therapy of one patient with an inflammatory disease (7). The case report described successful outcomes in a patient who did not respond to standard treatment. A limitation was that drug selection was empirical rather than based on systems-level understanding of the relative importance of disease-associated cell types, pathways, and genes.

Several systematic prediction models for drug selection in cancer exist, in which omics data are leveraged to determine the chemotherapies’ “killing potential” of tumour cells (8, 9). However, these models are not immediately translatable to IMIDs as they are 1) trained on large public drug-response data (e.g., GDSC database (10) and PRISM (11)), which are thus far unavailable for IMIDs, and 2) pursuing the eradication of disease-associated cell types. Rather few methodologies are applicable to IMIDs, including 1) identification of all druggable targets (12, 13), 2) targeting enriched pathways (13, 14), 3) network-based proximity calculations (6, 15) or 4) matching of transcriptomic signatures as by Connectivity Map (CMap) (16). A limitation of these approaches is that they are developed using bulk transcriptomics or genetic variants and hence do not possess inherent solutions for rank aggregation for parallel analyses of several cell types, which limits their applicability to scRNA-seq.

Aiming to create a systematic framework for scRNA-seq-based drug prioritisation and repositioning in inflammatory diseases, we hypothesised that the limitations of previous methodologies could be overcome by transposing network-based approaches (6, 15) to a systematic and scalable strategy for network-based virtual drug screening of multicellular disease models (MCDMs). Therefore, we composed a computational framework henceforth referred to as scDrugPrio (**Fig. 1**). Using scRNA-seq-derived differentially expressed genes (DEGs) of either 1) one individual or 2) a group comparison between patients and controls, scDrugPrio starts by identifying cell type-specific drug candidates by considering both proximity in a protein‒protein interaction network and biopharmacological criteria. To rank drug candidates, scDrugPrio calculates two measures, intracellular and extracellular centrality. We used these two measures to capture two important drug properties, namely, 1) proficiency in targeting key disease-associated expression changes in a cell type and 2) the relative importance of the targeted cell type. These measures are then aggregated over all cell types to provide a final drug ranking.

**Fig. 1.**
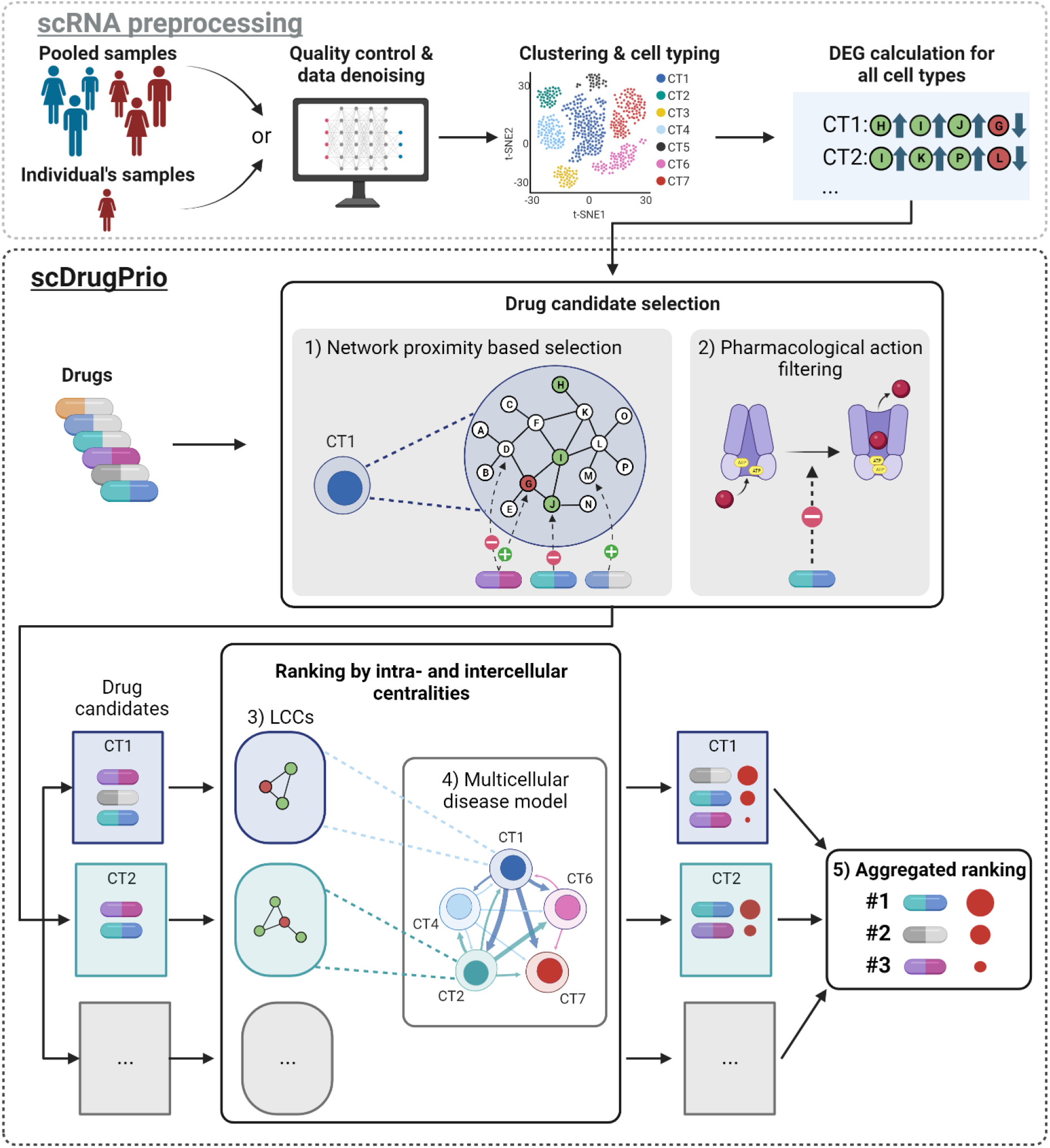
Overview of the scDrugPrio workflow. Single-cell RNA-sequencing (scRNA-seq) data from either individuals or groups of patients are preprocessed by undergoing quality control, denoising, clustering, cell typing and differentially expressed gene (DEG) calculation. DEGs for each cell type were calculated between healthy and sick samples. Using DEGs alongside information on drugs, scDrugPrio selects drug candidates (for each cell type; CT) whose gene targets are **1)** in network proximity to DEGs and **2)** who counteract disease-associated expression changes. These cell type-specific drug candidates are next ranked using intracellular and intercellular centrality. **3)** Intracellular centrality is computed based on the centrality of drug targets in the largest connected component (LCC) formed by DEGs and functions as a proxy for drug target importance. **4)** Intercellular centrality measures centrality in disease-associated cellular crosstalk networks called multicellular disease models (MCDMs). **5)** To derive a final ranking that aggregated cell type-specific drug selection and ranking into one list, drug candidates were ranked using a composite score of intra- and intercellular centralities (**Fig. S1**).

Because of the complexity and heterogeneity of IMIDs, we started by developing scDrugPrio using scRNA-seq data from a mouse model of antigen-induced arthritis. This reduced heterogeneity since the mice are inbred and the disease induced in a standardised way. Moreover, the mouse model allowed extensive *in vitro* and *in vivo* validation studies. To illustrate some potential case-of-use scenarios, we next applied scDrugPrio to cerebrospinal fluid from multiple sclerosis patients and intestinal biopsies from Crohn’s disease (CD) patients. Our analyses demonstrated drug selection and ranking capabilities through 1) the prioritisation of known drugs. For antigen-induced arthritis, drug selection was also supported by 2) experimental validation of repurposed drugs and 3) favorable comparison with previous methods. Next, we applied scDrugPrio to paired biopsies from individual CD patients, revealing its ability to capture the significant heterogeneity in individualised therapeutic prioritisations.

## Results

### The scDrugPrio framework

Aiming to create an analytical framework for scRNA-seq-driven drug prioritisation, we constructed scDrugPrio, which consists of three main modules: 1) drug candidate selection based on cell type-specific DEGs, 2) drug candidate ranking and 3) aggregated ranking of drug candidates from all cell types. For this scDrugPrio requires two components (**Fig. 1**): a) differentially expressed genes (DEGs) between sick and healthy samples for each cell type and b) drug data, including information on gene drug targets and pharmacological effects.

Preprocessing includes the calculation of DEGs based on scRNA expression from one or more healthy and one or more sick samples (**Fig. S1**). Preprocessing starts with quality control, batch correction (if needed) and data denoising of all scRNA-seq data, followed by clustering and cell typing. DEGs are computed per cell type by using sick versus healthy expressions. In the analysis below, we computed DEGs in the two modes. In the first mode, we use data from groups of sick and healthy individuals, while in the second mode, we use scRNA-seq data from paired samples of sick and adjacent healthy tissue for personalised drug prioritisations for one patient of interest.

scDrugPrio starts by computing the mean closest distance between DEGs and gene targets of drugs (henceforth referred to as drug targets) in the protein‒protein interaction network (PPIN) for each cell type and each drug candidate. Intuitively, the “closer” the drug targets are to DEGs, the better is the chance for the drug to affect the disease-associated genes (15). Specifically, a relative proximity measure (z_c_) capturing the statistical significance of the observed closest distance (d_c_) was calculated based on a comparison of d_c_ to the random expectation. Furthermore, scDrugPrio assumed that drugs that counteract the fold-change of at least one DEG will have a better chance to reverse disease-associated expression changes in the targeted cell type (following a similar idea as CMap (16)). To determine whether a drug counteracted a DEGs fold change, we considered 1) the direction of the fold change (upregulated or downregulated) and 2) pharmacological action (e.g., agonistic, antagonistic) on the targeted DEG. For each cell type, a list of drug candidates was derived by filtering out drugs with low network proximity (d_c_≥1, z_c_≥-1.64 corresponding to one-sided P < 0.05) and drugs that did not exert counteracting pharmacological action against at least one targeted DEG.

In the next step, scDrugPrio computed intra- and intercellular centrality measures that were later used to aggregate the prediction and rank drugs. To compute intracellular centralities per cell type, scDrugPrio determines the largest connected component formed by DEGs in the PPIN and next computes a drug’s centrality score based on the centrality of the drug targets within this component. Intracellular centrality hence presents an approximation of a drug’s target relevance for disease-associated expression changes in a cell type. For intercellular centrality, scDrugPrio constructs a multicellular disease model (MCDM). In short, MCDMs were based on predicted molecular interactions between differentially expressed upstream regulatory genes in any cell type and their downstream genes in any other cell type using NicheNet (17). The resulting MCDM is a network in which cell types were nodes connected by directed, weighted ligand-target interactions. Intercellular centrality refers to the centrality of cell types in the MCDM.

Finally, scDrugPrio aggregates drug predictions through aggregation of cell type-specific drug predictions and intracellular and intercellular centralities into a compound score (**Fig. 1**). The resulting scores were used to rank drugs. Hence, overall drug ranking prioritised drugs that targeted key disease-associated expression changes in the most important cell types.

### scDrugPrio development and evaluation in antigen-induced arthritis

scRNA-seq data were generated from whole joints of five inbred AIA mice and four naïve mice. After application of quality criteria (**File 1**), data included 16,751 cells with 132,459 mean reads per cell. Data for all mice were denoised jointly using a deep count autoencoder network (DCA) (18), and clusters were cell typed using marker gene expression (**Fig. 2a & S2**). Comparison of cells from AIA and naïve mice identified DEGs in 16 cell types (**Fig. 2, Supplementary Results**).

**Fig. 2.**
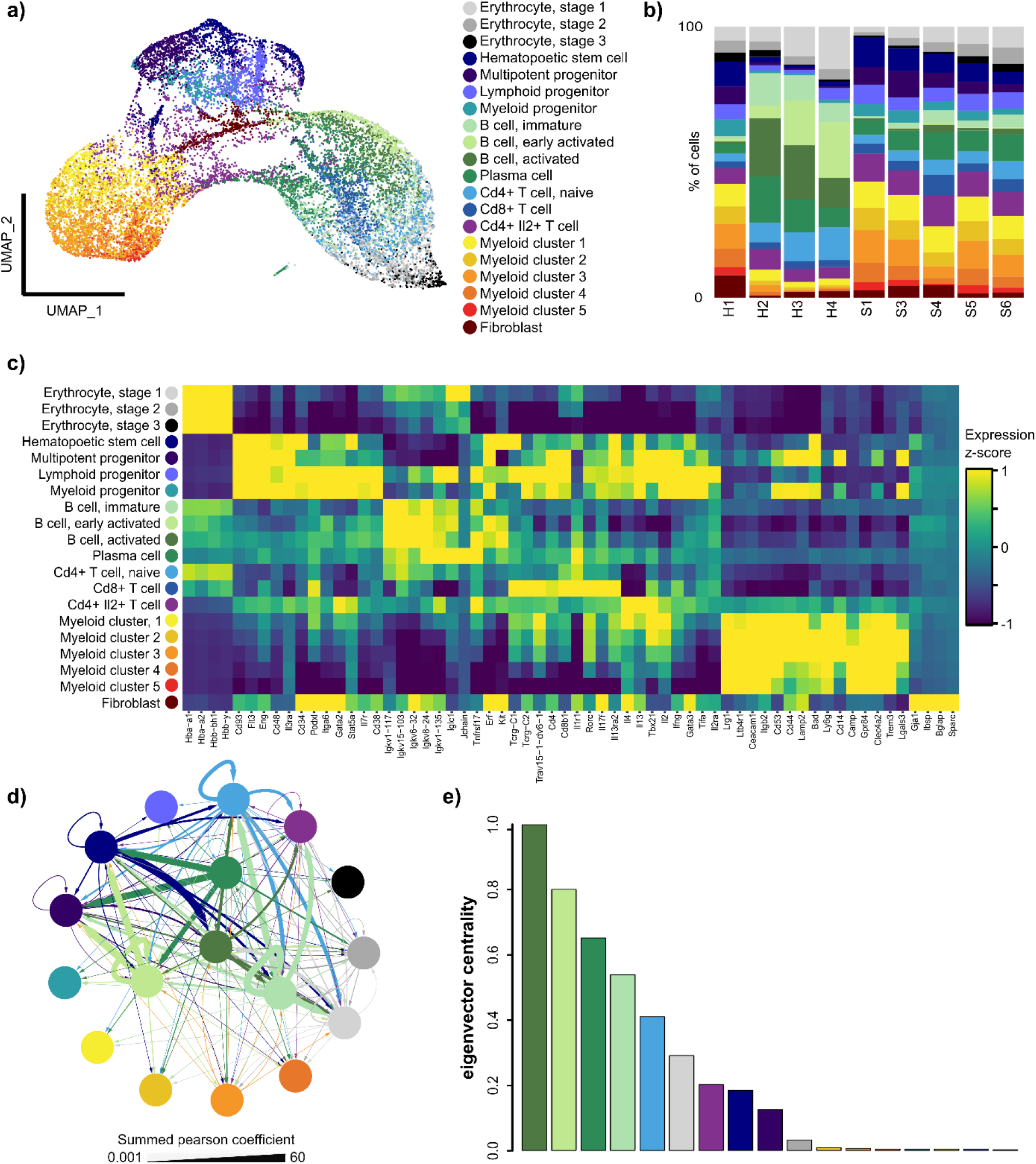
Construction and analysis of a multicellular disease model (MCDM) based on scRNA-seq analysis of a mouse model of antigen-induced arthritis (AIA). **a)** tSNE of cells (n = 16,751) pooled from all samples. **b)** Cell type proportions (%) for individual AIA (S1 to S6) and control (H1 to H4) mice. **c)** Heatmap of z-scores of the cluster-based means of normalised, denoised gene expression for cell type markers. **d)** MCDM, in which cell types were represented by nodes connected by the predicted interactions between upstream regulatory genes in any cell type and their downstream target genes in any other cell type (17). Edge width corresponds to the sum of Pearson coefficients for all interactions between two cell types, with arrows directed from upstream to downstream cell types. Edge color corresponds to the upstream cell type. As indicated by more central positions in the MCDM, the four B-cell clusters were the most central cell types. **e)** Bar plot depicting eigenvector cell type centrality in MCDM. Colours in all plots correspond to cell type colours in a).

We retrieved drug target information from DrugBank (19) for 13,339 drugs (**File 1**). From those drugs, we selected all of which had been FDA approved for human use and had at least one target in the human interactome (n = 1,840). According to the indication information in DrugBank, 57 drugs were FDA approved for RA and hence treated as true positives for the calculation of precision and recall.

Applying scDrugPrio, the final list of candidates included 334 out of 1,840 drugs; 32 drugs were established RA drugs, of which 22 ranked among the top 100 candidates (Fisher exact *P* < 10^-6^; **Fig. 3, File 1**). To further evaluate the candidates’ relevance, we collected clinical trial data from <u>clinicaltrials.gov</u> to capture the medical community’s interest in the identified candidates as RA medications. We also performed a literature review of the top 100 drugs to evaluate whether candidates had shown promise when tested in human RA or murine/rat RA models. Through a literature review, evidence for the relevance of 40 additional drugs was identified, whereas three of the top 100 ranking drugs had not shown effects in previous trials. Hence, 62.0% of the top 100 ranking candidates were either approved or had successful experimental validation in prior literature (**Fig. 3e**), and 95.4% of previously studied top 100 candidates had shown promise.

**Fig. 3.**
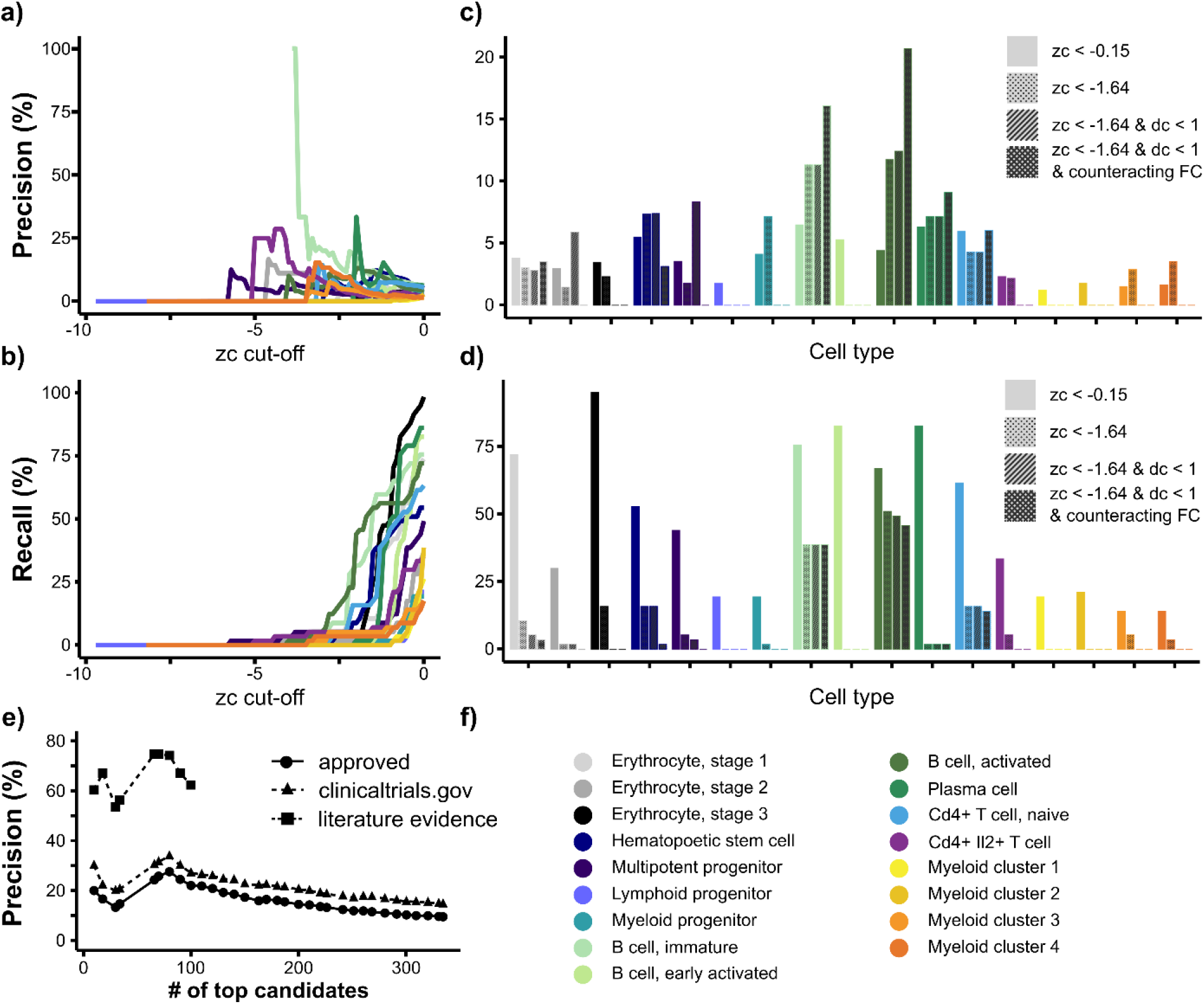
Precision and recall in relation to drug selection and ranking criteria. **a-b)** Precision and recall at different z_c_ cut-offs. In most cell types, precision increased with decreasing z_c._ Too stringent z_c_ cut-offs, by contrast, led to the exclusion of almost all candidates, including approved RA drugs. **c-d)** Precision and recall after stepwise application of drug selection criteria. Bars with no pattern represent a zc cut-off that Guney et al.^10^ had previously found to offer good coverage of known drug-disease pairs. The dotted pattern represents precision at z_c_ < -1.64 (corresponding to one-sided *P* < 0.05). The striped pattern represents precision among candidates with z_c_ < -1.64 and d_c_ < 1 (in other words, candidates passing network proximity-based selection). The crosshatch pattern represents candidates passing network proximity and pharmacological action selection. The application of selection criteria substantially increased the precision among central cell types. **e)** Precision for approved RA drugs and drugs with literature evidence among the ranked list of selected candidates. Drug ranking included rank ties. Literature evidence was gathered for the top 100 ranked candidates and is presented as triangles with a dashed line. The colour legend for **a-d)** is depicted in **f)**.

Describing scDrugPrio results in more detail, network proximity selection (d_c_<1, z_c_<-1.64) yielded no drug candidates for eight cell types and an average of 67 drug candidates (min = 2, max = 226) for the remaining twelve cell types. Precision among cell type-specific candidates ranged from 0.0% to 12.8%. However, precision of cell types that were central in the MCDM, such as activated, immature, and plasma B cells (12.8%, 11.4%, and 6.3%, respectively), outperformed random selection from the 1,840 included drugs (57/1,840 = 3.1%; Fisher exact *P* < 10^-12^, *P* < 10^-8^, *P* = 0.4, respectively). Following network proximity calculations that were performed in the absence of information on a drug’s effect on the target, we found tasonermin, a synthetic version of TNF, among the top-ranking candidates for several B-cell subtypes. Since overexpression of TNF has a crucial role in RA pathogenesis (20), this ranking supported the network proximity criterion. However, because tasonermin mimics the effects of TNF, it could worsen the disease. This finding exemplified the importance of pharmacological action selection. Pharmacological action selection resulted in an average of 43 drug candidates per cell type (min = 2, max = 137). Overall precision decreased to a median [range] of 1.56% [0.00 - 20.63], although it increased in activated, immature, and plasma B cells (20.6%, 16.1%, and 9.1%, respectively) as well as in T cells (**Fig. 3c**)

Having identified drug candidates for every cell type individually, intra- and intercellular centrality were calculated. The use of intra- and intercellular centrality measures for composite ranking was motivated by our findings that 1) intercellular centrality correlated with the significance of GWAS enrichment among cell type-specific DEGs (Pearson’s *r* [95% CI] = 0.62 [0.21 – 0.85]; P < 0.01), 2) intercellular centrality correlated with the precision among cell type-specific drug candidates (Pearson’s *r* [95% CI] = 0.77 [0.46 – 0.91]; P < 10^-3^; **Fig. S2j**), 3) drugs that targeted more than one cell type were more likely to be known RA drugs (**Fig. S2k**) and 4) intracellular centrality could improve the mean rank of known drugs more than expected by chance (**Supplementary Results**).

Using the AIA and human RA data, we benchmarked scDrugPrio against previous methodologies (**Supplementary Results**) and performed extensive additional testing, demonstrating the advantage of scRNA-seq-driven analysis over genetic variations or bulk transcriptomics. Furthermore, we evaluated drug selection criteria and performed robustness analysis (**Supplementary Results).**

### Experimental validation of scDrugPrio

To further validate scDrugPrio, five high-ranking drugs that were not found to have any prior literature support for their efficacy in RA were chosen alongside the highest-ranking RA drug (auranofin #6) serving as a positive control. We first evaluated the five drugs by *in vitro* studies of B cells. This cell type was selected because of its crucial role in the pathogenesis of RA and its central position in MCDM (21). We used previously described *in vitro* models of B-cell functions (21) measuring murine B-cell survival, activation, proliferation, and antibody production upon *in vitro* stimulation with selected drugs at various concentrations. Auranofin dramatically suppressed B-cell functions, including cell viability, proliferation, and immunoglobulin production (**Fig. 4**). Additionally, two of the five novel drugs (#114 amrinone and #100 adapalene) showed concentration-dependent *in vitro* effects on B-cell viability, proliferation, and immunoglobulin production (**Fig. 4**). Adapalene also greatly inhibited murine B-cell activation (**Fig. 4b**). The other three drugs (#91 irbesartan, #98 isosorbide, and #134 dimethyl furamate) showed little to no effect on murine B-cell functions (**Fig. S3**). Next, we examined the effects of drug candidates on the function of human B cells. Similarly, auranofin, adapalene and amrinone inhibited human B-cell viability, activation, proliferation, and IgG production (**Fig. S4**). Thus, two out of five candidate drugs prioritised by scDrugPrio, namely, adapalene and amrinone, were successfully validated by *in vitro* studies.

**Fig. 4.**
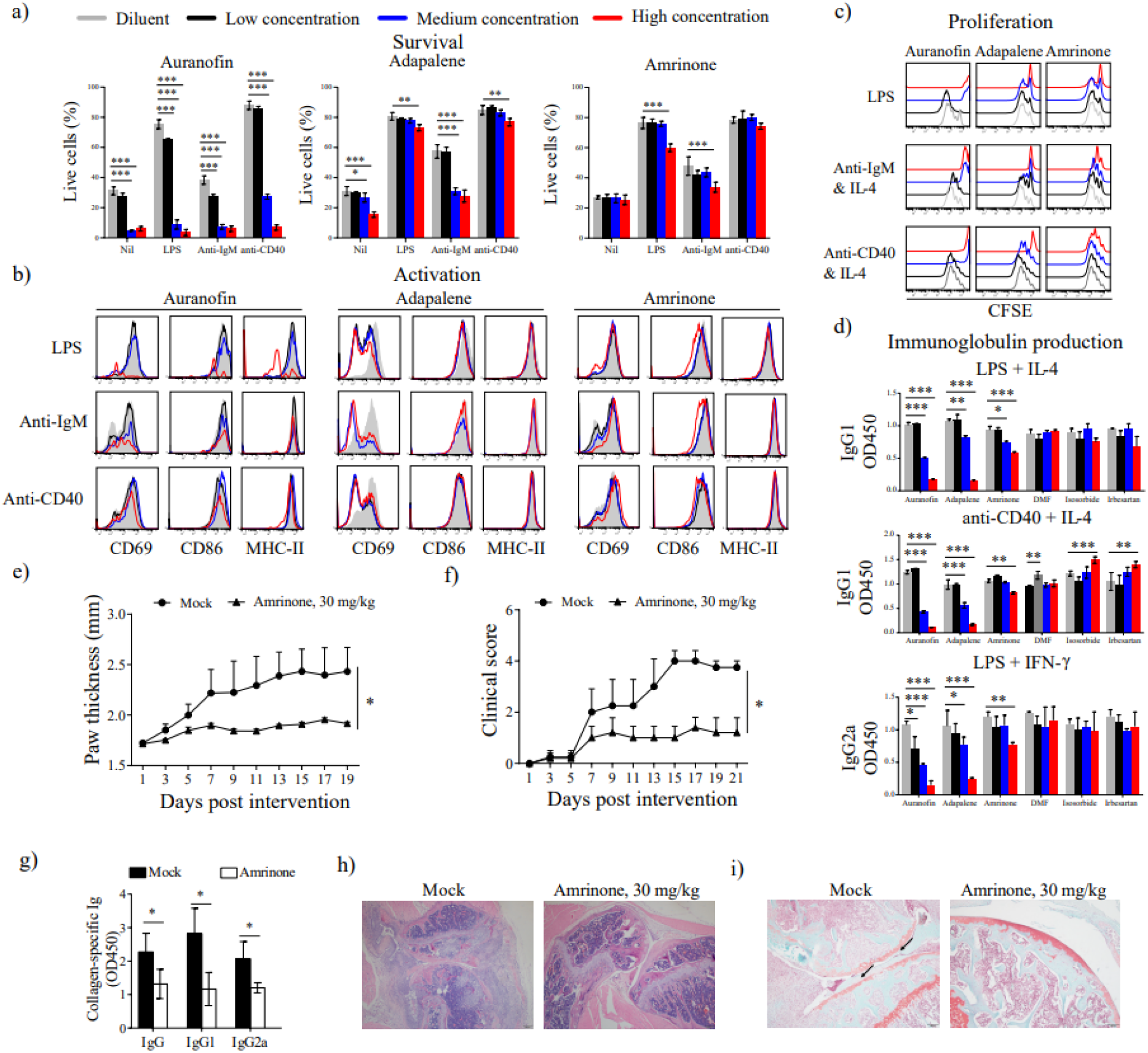
Experimental validation of amrinone in suppressing murine B-cell function and the pathogenesis of the CIA mouse model. Drug effect of the selected drugs and the positive control on *in vitro* murine **a**) B-cell survival, **b**) activation, **c**) proliferation, and **d**) immunoglobulin production. Colours represent the responses to diluent and different drug concentrations (low, medium, high); for specific concentrations, see **Table S1.** Having successfully validated adapalene and amrinone *in vitro*, we conducted *in vivo* experiments for amrinone. Mice with collagen-induced arthritis (CIA) were treated with diluent (n = 5) or amrinone (30 mg/kg, n = 5) for 3 weeks. The **e)** rear paw thickness, **f)** clinical scores, and **g)** collagen-specific serum autoantibodies were measured. Furthermore, drug efficacy was assessed by analysis of **h)** joint immune cell infiltration using H&E staining and **i)** bone erosion using Safranin-O staining. * P < 0.05, **P < 0.01, ***P < 0.001. Dimethyl furamate, DMF.

Valid drug candidates should arguably be transposable and replicable. As drug prediction was performed using data from the AIA model of arthritis, we deployed the collagen-induced arthritis (CIA) model for further *in vivo* study. We selected only amrinone for further study, as adapalene was designed for topical skin use and not for systemic delivery. CIA mice were administered 30 mg/kg amrinone i.g. for 3 weeks. This treatment significantly reduced the paw thickness (**Fig. 4e**) and clinical scores of CIA mice (**Fig. 4f**). Analysis of collagen-specific serum autoantibodies revealed a significant inhibitory effect (**Fig. 4g**). Considering that it even reduced immune cell infiltration (**Fig. 4h**) and bone erosion (**Fig. 4i**), *in vivo* studies confirmed the relevance of amrinone treatment and thereby further supported scDrugPrio.

### Application of scDrugPrio to multiple sclerosis

We next applied scDrugPrio to human IMIDs using scRNA-seq data of cerebrospinal fluid (CSF) from multiple sclerosis (MS) patients and idiopathic intracranial hypertension that served as controls (22). After application of quality cut-offs (**File 2**), the data included 33,848 cells with 47,332 mean reads per cell. Comparing MS samples with controls, DEGs were calculated from batch-corrected, normalised expression scores. scDrugPrio identified on average 59 (min = 1, max = 270) drugs in 19 cell types (**Fig. 5 & S5**). Aggregated ranking of 417 drug candidates (**Fig. S6, File 2**) included ten out of 17 approved MS drugs. Among the top 100 ranking candidates, four approved MS drugs, as well as biosimilars rituximab and obinutuzumab, were identified, which outperformed random expectation (precision 0.9%). A literature search revealed that an additional 22 of the top 100 ranking drugs had shown effects in previous studies, resulting in a precision of 28.3% (**Fig. 5d**). Seventy-one drugs had not yet been validated in previous studies, and one evaluated drug had not shown promise in previous studies. Hence, 96.7% of the studied candidates showed efficacy. Taken together, these data supported the potential of scDrugPrio to predict the response to drugs approved for that disease, as well as for repurposing other drugs.

**Fig. 5.**
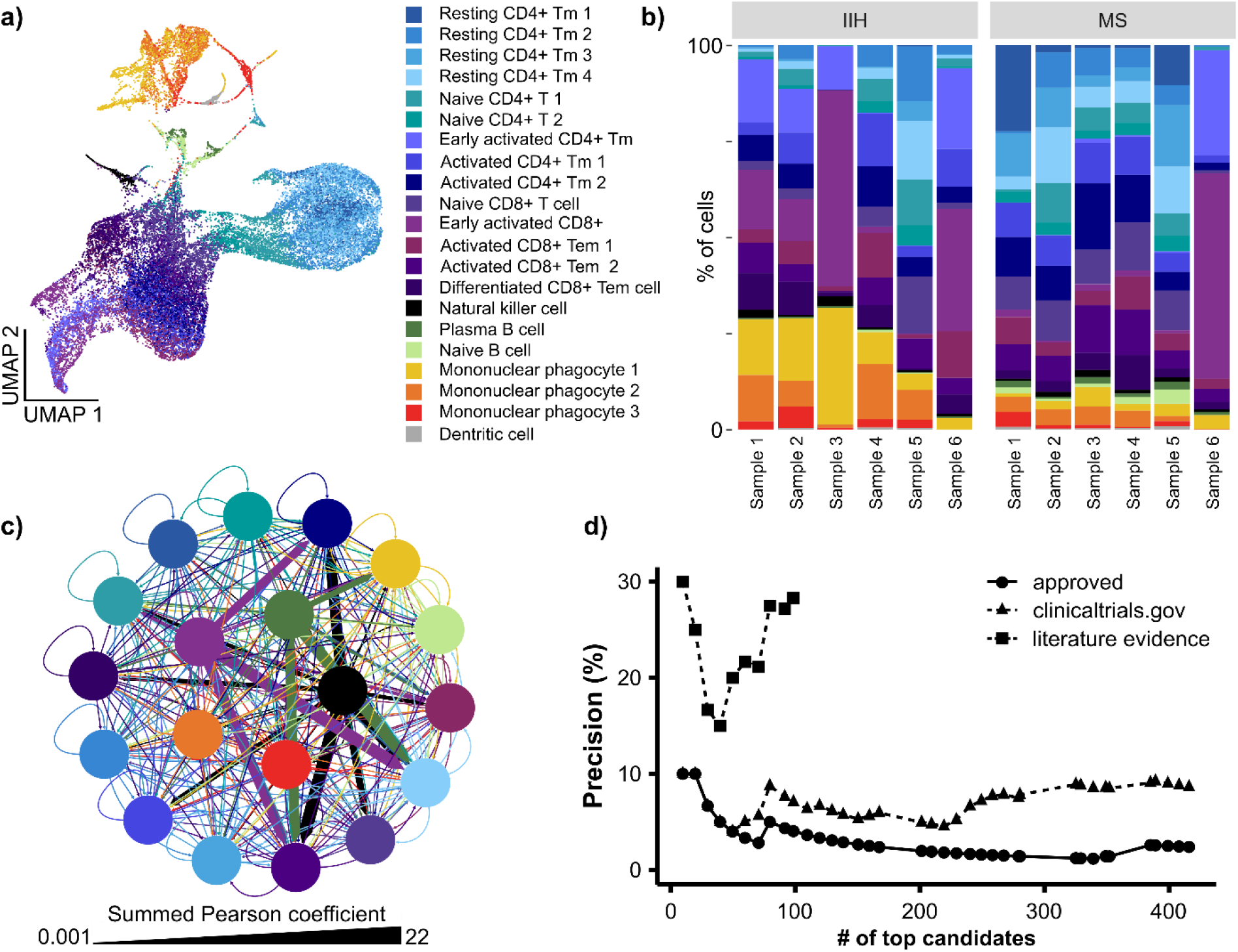
scDrugPrio applied to scRNA-seq data from cerebrospinal fluid samples of MS patients and controls. **a-b)** Clustering and cell type proportions (%) for CSF samples of idiopathic intracranial hypertension (IIH) and multiple sclerosis patients (MS). Cell type proportions showed significant interindividual differences among MS patients. **c)** The MCDM for MS was more complex than that of AIA mice. The most central cells included mononuclear phagocyte 3, natural killer cells, mononuclear phagocyte 2, early activated CD8+ T cells and plasma B cells. (in order of eigenvector centrality). **d)** Precision among ranked candidates for approved MS drugs and the top 100 drugs with literature-derived evidence. Abbreviations: Tm, T memory; Tem, T effector memory.

The value of above rank aggregation was supported as precision among cell type-specific drug candidates only ranging from 0.0% to 12.5%. The MCDM (**Fig. 5c, File 2**) indicated plasma cells, mononuclear phagocytes, natural killer cells, and activated CD8+ T cells to be of central importance, which is consistent with the current pathophysiological understanding (22, 23). Notably, the MS MCDM was more complex than the AIA MCDM regarding the number of cell types and interactions.

### Application of scDrugPrio to Crohn’s disease

Next, scDrugPrio was applied to patients with Crohn’s disease (CD) from whom scRNA-seq data (24) from paired inflamed and uninflamed intestinal tissue biopsies were available. After application of quality criteria (**File 3**), 77,416 cells were included with 3,591 mean unique molecular identifiers (UMIs) per cell. Following preprocessing, analysis was performed on batch-corrected, pooled data from all patients in which DEGs were calculated through comparison of inflamed and uninflamed samples (**Fig. S7 & S8, File 3**). The aggregated ranking included 343 drug candidates, of which five were known CD drugs. These five consisted of sulfasalazine (#91), mesalazine (#169.5), rifaximin (#238) and two anti-TNF drugs (adalimumab and infliximab, tied rank #292.5). Apart from sulfasalazine, literature evidence suggested the effectiveness of 13 additional top-ranking 100 drugs, resulting in a precision of 14%.

### Patient heterogeneity in human disease

An important difference between human data and mouse data was substantial interindividual heterogeneity in human patients. To illuminate such differences, we compared 1) cell type proportions, 2) gene expression profiles and 3) examination of latent features of the non-batch-corrected data. Mice, as expected, showed no differences in cell type proportions (chi-square P = 0.9919; **Fig. 2c**), gene expression (95% CI of misclassification rate 0.504∼0.532 in the training data and 0.638-0.655 in the test data), or latent features (**Fig. S2c**). However, both MS and CD patients showed great interindividual heterogeneity in cell type composition (chi-square *P* < 10^-15^, respectively). Heterogeneity in cell type composition, although decreased, can still be observed in the batch-corrected data (**Fig 5b & S7i**). A patient effect was also observed in the non-batch-corrected gene expression for MS (random forest, 95% CI of misclassification rate 0.075 - 0.084 in training data and 0.124 - 0.132 in testing data) and CD patients (random forest; 95% CI of misclassification rate 0.270 – 0.283 in training data and 0.378 ∼ 0.385 in testing data) as well as latent features derived from non-batch-corrected data (**Fig. S5 & S8**). Taken together, such patient effects necessitated batch correction for pooled prediction using the human data sets above.

### Potential for individualised predictions

Even though batch correction had been applied appropriately, a potential limitation of the above MS and CD analyses was that scDrugPrio was applied to pooled data derived from heterogeneous patients and controls. Patient effects might form the bases for the responder/nonresponder dichotomy, and we therefore evaluated scDrugPrios potential for individualised drug prioritisation. For this, we applied scDrugPrio to individual CD patients, using similar preprocessing as for CD data above with the following exceptions: 1) data were not batch-corrected, and 2) following denoising, cells from inflamed and uninflamed samples of each patient were clustered separately. DEGs were derived through comparison of individual patient inflamed and uninflamed cells in each cluster. DEGs showed that the eleven patients expressed important CD drug targets differently (**Fig. S9a-b**). To investigate whether such molecular differences could affect drug prediction outcomes, scDrugPrio was applied to all patients separately (**Fig. S9-11, File 4**).

Strikingly, individualised drug predictions of nine out of eleven patients (such as patient 1: 19.0%; patient 10: 20.5%, **Fig. 6 & S12**) outperformed the precision of pooled patient analysis (14.0%). Among the top 10 candidates, precisions for individualised predictions (20-70%) outperformed precision of pooled patient analysis in seven patients and equalled that of pooled patient analysis in four patients (10%). All predictions outperformed random chance (1.5%). More detailed analysis revealed interindividual differences in cell type proportions and network properties in the MCDM (**Fig. S11**) as well as different drug rankings (**Fig. S13**). Taken together, these findings supported that scDrugPrio presents a valid strategy for personalised drug prioritisation.

**Fig. 6.**
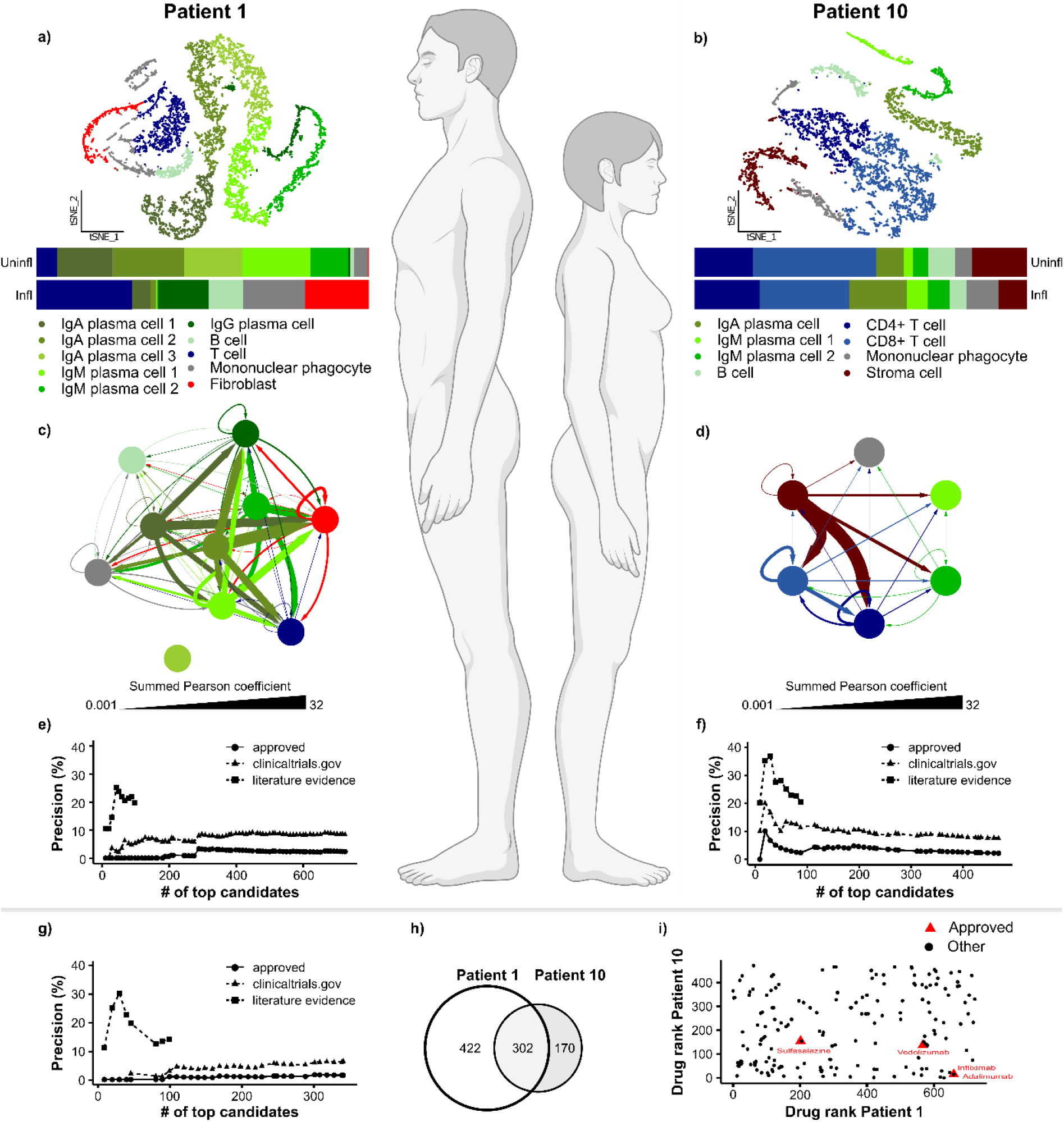
scDrugPrio for individual drug prediction. **a-b)** Cell type proportions differed greatly between two CD patients, as shown in the horizontally stacked bar plots representing paired biopsies from inflamed (infl) and uninflamed (uninfl) lesions that were taken from each patient. **c-d)** Patients also show differences in the composition and interconnectivity (representing ligand interactions) of the MCDMs. Patient 1 had a cell type for which no ligand-target interactions could be found with any other cell types in the MCDM. **e-f)** The precision for the ranked drug candidates for patient 1 was low for approved CD drugs, while literature evidence supported the top-ranking drugs, of which many are anti-inflammatory. In contrast, patient 10 had several approved CD drugs among the top 100 candidates, and a curated literature search confirmed the validity of many more candidates. **g)** Precision for prediction based on pooled patient data was poor. **h)** Venn diagram presenting the overlap of considered drug candidates for patients 1 and 10. **i)** Interindividual differences between patients 1 and 10 were reflected in the prediction outcome, as no correlation existed between the drug rank of drugs that were candidates in both patients.

To exemplify the potential of scDrugPrio for individual patients, we next compared two patients who previously had been classified as an anti-TNF responder (patient 10) and nonresponder (patient 1) based on a cellular signature score (24). In agreement with the previous classification of anti-TNF response (24), TNF had a more central role in the MCDM of patient 10 (**Fig. S10m, n**). Hence, it was not surprising that aggregated drunk ranking ranked adalimumab (anti-TNF) higher for patient 10 (#15.5) than for patient 1 (#658). As expected, adalimumab was the highest-ranking approved CD drug in patient 10. For patient 1, scDrugPrio prioritised other immunomodulatory drugs over anti-TNF treatment, namely, natalizumab (#19), human intravenous immune globulin (#21), basiliximab (#25), sarilumab (#29), and other approved CD drugs, such as methotrexate (#188) and sulfasalazine (#202).

### Application of scDrugPrio to nonresponder/responder data from patients with psoriatic arthritis highlighted the importance of local tissue samples

Since the previous analyses supported scDrugPrio’s potential for another case-of-use scenario, namely, to distinguish drug responders from nonresponders. To explore this potential, we collected peripheral mononuclear blood cells (PBMCs) from patients with psoriatic arthritis (PsA) as well as healthy controls. PBMCs were chosen because the analysis of blood samples is clinically more tractable than the analysis of biopsies. Samples were cryopreserved before treatment with either anti-TNF or anti-IL17. Treatment response was later assessed by a rheumatologist according to EULAR response criteria (25) (**File 5**). We next selected 32 patients, of whom eight were classified as responders (R) and eight as nonresponders (NR) to either of the two drugs, as well as eight healthy controls. Cryopreserved PBMCs from these patients were analysed with scRNA-seq. For preprocessing, data were divided into two data sets by treatment regimen, each data set containing the corresponding eight R and NR along with the eight healthy controls. After application of quality cut-offs, the data included 78,610 cells with 5,088 mean reads for anti-TNF analysis and 72,472 cells with 5,343 mean reads for anti-IL17 analysis. Data were batch-corrected and DCA denoised before clusters were identified and cell-typed using marker genes (**Fig. S14 & S15**, **File 5**).

For each data set, DEGs were calculated through comparison of cells from healthy controls to either R or NR. The precision for approved PsA drugs among the top 100 candidates in the respective aggregated ranking was 0% for anti-IL17 NR, 4% for anti-IL17 R, 1% for anti-TNF NR and 1% for anti-TNF R (**Fig. S14 & S15**). Unexpectedly, anti-TNF treatment received a low rank in anti-TNF R but not in NR (#333 and #88, respectively), while anti-IL-17 was not considered a candidate in either R or NR.

Further analyses of our PBMC data from all R and NR patients showed that the TNF signaling pathway was significant in only 8% and 8% of clusters, respectively. The corresponding figures for the IL-17 signaling pathway were 17% and 13% in the anti-IL17 R and NR groups, respectively. In those cell types, most pathways were downregulated, including those regulated by *TNF* and *IL17* (**Fig. S16)**. This result contrasted with previous studies of skin and synovium from PsA, which showed increased expression of *TNF* and *IL17*, as well as their pathways (26, 27). A similar dichotomy between local inflamed tissues and cells in the blood in autoimmune diseases has been previously described (28). This dichotomy can be explained by the physiological need to localise inflammatory responses to inhibit systemic and possibly fatal responses. The general clinical implication may be that drug predictions should ideally be based on local tissue samples (25).

## Discussion

The main problem in therapeutics, which serves as the basis for this study, is the large number of IMID patients who do not respond to treatment (2, 3, 29). Previous virtual drug screening methods for inflammatory diseases are based on genetic variance or bulk RNA sequencing (15, 30, 31) and hence do not consider variations in gene expression across different cell types, biopharmacological properties, or individual variations between patients with the same diagnosis. While harnessing the daunting complexity and heterogeneity for personalised treatment may seem impossible by health care standards today, this challenge should be put in the context of the suffering and costs resulting from ineffective drug treatment. Many IMIDs cause life-long morbidity and increased mortality. The yearly cost of treating an individual IMID patient may be hundreds of thousands of dollars for drugs and hospital care (1).

Recent efforts for drug toxicity screening (8, 32) support the feasibility of scRNA-seq to capture relevant cellular information. However, systematic solutions for drug prioritisation for IMIDs based on scRNA-seq remain to be devised. We therefore propose a computational framework, scDrugPrio, that extends on existing bioinformatic tools (6, 15, 17) by providing a framework for data integration, enabling drug ranking based on a multifaceted understanding of cell type-specific disease mechanisms, altered cellular crosstalk and pharmacological effects. We demonstrate that scDrugPrio yields relevant and robust drug prioritisations, outperforms previous methods (14, 33) and holds potential for individualised as well as pooled drug prioritisation and repurposing.

An important advance of scDrugPrio is that it can be applied to scRNA-seq data. The importance lies in the fact that complex diseases each involve differential expression of thousands of genes across multiple cell types (6). A previous case report (7) described one successful example of treating an individual patient with immunological diseases based on scRNA-seq data. However, the drug choices were empirical rather than systematic. Because scRNA-seq allows transcriptome-wide analyses in each of thousands of cells, it is possible to infer disease-associated changes in individual patients preferably by comparisons with noninflamed samples from the same individual or to groups of healthy individuals. Thus, scDrugPrio has the potential to personalise the treatment of individual patients. The importance of this advance is highlighted by our results and previous findings (24, 34) showing great interindividual differences in the molecular and cellular composition of human diseases. For example, we showed that scDrugPrio ranked anti-TNF treatment high in a CD patient who was classified as a responder but not in a nonresponding patient. In the latter patient, other immunomodulatory drugs, such as natalizumab, received high ranks. Natalizumab is mainly used in MS but has, in previous studies, shown positive effects in CD (35), making it a viable recommendation. These examples emphasise that successful drug screening will need to consider variations between patients with the same diagnosis.

There are several limitations of scRNA-seq-based drug predictions in IMIDs. Many of these depend on the challenges involved in harnessing complex and heterogeneous disease-associated changes with an emerging technology such as scRNA-seq. An analogy to a historical example may illustrate how such limitations may drive scientific progress. In 1970, Needleman‒Wunsch (36) and 1981, Smith‒Waterman (37) published algorithms for global and local sequence alignment, which were widely used. The limitations of those algorithms were that they were mainly useful for nucleotide but not protein sequence analyses because of limited protein sequence data and no scoring system that modelled protein evolution. During the next two decades, these problems were resolved by increasingly accurate data and methods (38). Importantly, 42 years after publication of the Smith‒Waterman algorithms for proteins, these algorithms can generate very accurate results when combined with scoring systems that were later developed (38). We propose that the limitations of scRNA-seq that we face today will lead to a similar development of increasingly accurate technologies. One obvious limitation of scRNA-seq is that mRNA and protein levels may be poorly correlated, which can limit biological interpretability. From this perspective, the use of DEGs for scDrugPrio’s pharmacological predictions is a relative strength, as DEGs have been shown to correlate significantly better with protein levels (39) compared to mRNA levels alone and hence increase the biological relevance of our predictions. Inherent limitations of scDrugPrio also derive from the use of current interactomes, which are not comprehensive in terms of proteins, interactions, and variations across cell types (40) and are prone to investigative biases. While it is impossible to address all these concerns, we explored whether network proximity-based drug selection was influenced by investigative bias through replication of key results in a smaller yet unbiased interactome. We found that precision among candidates was slightly lower, partly due to missing drug targets in the interactome, but that results were comparable. Additionally, scDrugPrio might benefit from systematic parameter optimisation, which is currently not possible due to the limited amount of suitable scRNA-seq data sets.

Predictions were based on complex or partially uncharacterised drug-target effects, which may vary between different locations in the body. The need for better characterised drug effects and the relative importance of drug targets is exemplified by etanercept, which inhibits TNF and its receptors but may activate IgG receptors. The TNF-inhibitory effects are beneficial in PsA, while those on IgG receptors are not clearly defined. However, because all these targets were downregulated in PBMCs from nonresponding PsA patients, etanercept (counteracting IgG downregulation) received a higher rank than in patients responding to anti-TNF treatment. While unexpected, this highlights the need for systematic information about the relative importance of drug targets. Future efforts aiming to address these limitations might find that the predictive capability of scDrugPrio can be further enhanced by integration of binding affinity (e.g., BindingDB) or bioactivity (e.g., ChEMBL), especially if data become more comprehensive.

The above example of etanercept in PsA highlights an important clinical concern, which to our knowledge has not been recognised in the context of drug prediction methods. While analyses of blood samples are often more tractable in routine clinical practice, disease-associated mechanisms may vary greatly between cells in blood and inflamed tissues. Our scRNA-seq analyses of PBMCs from PsA patients who did or did not respond to treatment with either anti-TNF or anti-IL-17 showed that TNF and IL-17 signaling was found only in a small portion of the PBMC cell types and, in fact, was downregulated in both responders and nonresponders. In contrast, previous studies (26, 27) of synovium from PsA patients have shown consistent upregulation of both signaling pathways. Additionally, one previous study also showed differences between synovium and skin from the same patients (26). Thus, scDrugPrio should ideally be applied to local, inflamed tissue samples of the relevant tissue.

Despite these limitations, the translational relevance of scDrugPrio was supported by analyses of precision/recall for drugs that were approved for the studied diseases, as well as by *in vitro* and *in vivo* experiments. Those experiments implied two drugs, namely, adapalene (used for acne vulgaris) and amrinone (used for congestive heart failure), that had not been previously described as candidates for RA treatment. However, both have anti-inflammatory effects and could, therefore, be effective (41, 42). This potential was supported by *in vivo* experiments in which CIA mice were treated with amrinone (adapalene is a topical skin drug and hence is not suitable for systemic treatment in this experimental system). This example also suggests a potentially important pharmacological application of scDrugPrio, namely, virtual drug repurposing by systematic screening of thousands of drugs across several inflammatory diseases, as well as in patients who do not respond to standard treatment.

Here, we show that scDrugPrio has the potential for individualised drug predictions. We have made data and tools freely available for this purpose. However, further parameter optimisation and controlled, prospective clinical studies are needed for clinical translation. If successful, this approach could lead to a radical change in health care, which today is largely based on treating groups of patients with the same diagnosis with a limited number of drugs based on a limited understanding of the underlying molecular complexity and heterogeneity with limited population-based efficacy (43).

## Acknowledgement

This work was supported by This work is supported by Doctis project, which has received funding from the European Union’s Horizon 2020 Research and Innovation Programme under Grant Agreement N° 848028; Swedish Cancer Society CAN 2017/411; Cocozza Foundation; National Natural Science Foundation of China 82171791, US National Institutes of Health grants HL155107 and HL155096; American Heart Association grant 957729; European Union’s Horizon 2021 Research and Innovation Programme grant 101057619 and Mag-Tarmfonden (grant 1-23). The computations were partially enabled by resources provided by the Swedish National Infrastructure for Computing (SNIC) at Linköping University partially funded by the Swedish Research Council through grant agreement no. 2018-05973.

## Author contributions

SS designed and performed the bioinformatics and analyses, which were supervised by DG, OS, IAK, MG, and MB. HH, XL and HZ performed scRNA-seq. SL performed the scRNA-seq data extraction, which was led by DG and MB. SS, DE, MS and YZ analysed the data. SM and AJ were responsible for clinical studies of drug responses in patients with PsA. HW performed the experimental validation. SS, JL, DG, OS, and MB took the lead in writing the manuscript, and all authors provided critical feedback.

## Material & Methods

### scDrugPrio’s computational framework

As indicated in **Fig. S1**, scDrugPrio requires 1) an adjusted scRNA-seq matrix, 2) disease-associated differentially expressed genes (DEGs) for each cell type from either group-based comparison of healthy and sick samples or from inflamed and noninflamed samples of one individual, 3) a protein‒protein interaction network (PPIN) and 4) drug-target information. scDrugPrio then utilises this information for cell type-specific drug selection, calculation of drug ranking measures and finally rank aggregation.

For drug selection, scDrugPrio first computes the mean closest network distance (d_c_) between cell type-specific DEGs and drug targets in the PPIN for each cell type-drug combination. To calculate z-scores (z_c_) for network distance, permutation tests (1,000 iterations) were performed in which both cell type-specific DEGs and drug targets were randomised in a bin-adjusted manner (15) before the mean closest distance was calculated. The minimal bin size for randomisation was set at 100 genes. Drugs that did not have any target in the interactome were removed from the analysis (n = 4 for literature-curated PPIN). Based on network distance, we selected only drugs that targets were significantly close (z_c_ < -1.64 corresponding to one-sided P < 0.05) to DEGs and that frequently targeted DEGs directly (d_c_ < 1). These cut-offs were chosen based on our empirical observations (**Fig. 3a-d & S2h,j**) and previous knowledge (44). As the significance of network proximity can depend on the number of DEGs relative to the size of the network, cut-offs allowing only the top significant DEGs to enter analysis were implemented when needed. Cell type-specific drug candidates were selected further by requiring drugs to counteract the fold change of at least one targeted DEG. This criterion intuitively removes drugs that likely will not help to restore transcriptomic homeostasis. For this purpose, the pharmacological action of the drugs on their targets was determined. Binary drug action (activating/enhancing or inhibiting) on the drug target was recorded for each drug (**File 1-5)**. If the pharmacological effect of the drug on the target had not been specified explicitly in DrugBank (19), a literature search was performed using the drug name and gene symbol of the targeted DEG as search terms in PubMed and Google Scholar. Additional information gathered from the literature can be found in **Files 1-5**. In case the pharmacological effect of the drug on a target, despite a literature search, could not be classified as enhancing/activating or inhibiting, the drug target was assumed to not counteract fold-change.

Drug ranking by intra- and intercellular centralities was motivated by empirical observations (**Fig. 3e & S17**) in our study as well as previous indications of the biological importance of disease modules (6, 45) and central cell types in MCDMs (6). For calculation of intracellular centrality, disease modules for each cell type were defined as the largest connected component (LCC) formed by a cell type’s DEGs in the PPIN. For LCC identification, the igraph R package (46) was utilised. To avoid overparameterisation, the eigenvector centrality (47) of DEGs in the LCC was calculated using the CINNA R package (48). For each drug, the intracellular centrality was calculated as the geometric mean of its differentially expressed target centrality scores in the cell type-specific LCCs. If a drug did not target any DEG included in the LCC, intracellular centrality was set to zero.

Intercellular centrality was calculated using MCDMs that modelled disease-associated cellular crosstalk. For the creation of MCDMs, first, cell type interactions were predicted using NicheNet (17). Briefly, NicheNet predicts and ranks ligand–target links between interacting cells by combining their expression data with prior knowledge on signalling and gene regulatory networks. As suggested by Browaeys et al (17), Pearson correlation was used to measure each ligand’s ability to predict the gene expression of genes in the *gene set of interest* compared to *background genes* in the receiving cell type. This means that a ligand has a strong positive correlation coefficient if its cognate receptor and the downstream genes of that receptor are all differentially expressed in the downstream cell type. We downloaded the human ligand-target model as well as the human ligand‒receptor network (downloaded from https://zenodo.org/record/3260758 April 2020). Cell type-specific DEGs constituted the *gene set of interest*. A set of *potentially active ligands* was defined as the intersection of ligands included in the downloaded human ligand-target model and ligands among respective cell type DEGs. *Background genes* for each cell type were defined as genes (***i***) in the denoised single-cell expression matrix ***D*** of ***k*** cell type-associated cells that showed a mean aggregate expression, ***Ea(i)***, over ***Ea(i)*** ≥ 0.2. This definition of background genes was similar to definitions by Browaeys et al. (17) and Puram et al. (49). At the chosen cut-off, we identified ca. 10,000 background genes that corresponded to the recommended amount for NicheNet calculations (17).

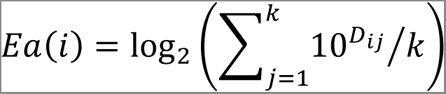

All genes were translated to human Entrez gene symbols using human-mouse orthologues downloaded from NCBI (August 2019). Ligand activity analysis in NicheNet was performed for all possible cell type pairs, including self-interactions, excluding cell types that did not express DEGs. In the next step of MCDM construction, directed cell type interactions were derived from ligand activity results and weighted by NicheNet-derived Pearson coefficients.

Only ligand interactions with a positive Pearson correlation coefficient were considered negative Pearson coefficients that reflected the association of a ligand with background genes and therefore were not biologically relevant. The resulting MCDM was visualised using Cytoscape 3.6.1 (50), and for visualisation purposes, the sum of Pearson coefficients that described the directed interaction between two cell types was used. Supplemental analysis supported the relevance of identified ligand interactions (**Supplementary Results**). Eigenvector centrality was calculated for each cell type based on the weighted, directed interactions in the MCDM using the igraph and CINNA R package (46, 48). The intercellular centrality of each drug was computed as the sum of MCDM centralities of the cell types that had selected the drug as a candidate. While eigenvector centrality is well tailored to capture central disease-associated cell types in the MCDM, considering both direct and indirect node connections (47), multiple centrality measures are available. We evaluated several of them, finding them to yield similar results to eigenvector centrality (**Supplementary Results, File 1-5**).

Final rank aggregation involved the calculation of a drug’s compounded intra- and intercellular centrality. For this, we calculated combined intracellular centrality for each drug as the sum of drug-specific intracellular centralities in all cell types. The centrality compound score consisted of a drug *intercellular centrality + 0.1 x combined intracellular centrality a*nd thereby emphasised the importance of intercellular centrality over intracellular centrality. Intracellular centrality effectively worked as a tiebreaker. Drugs were ranked based on centrality compound scores, using the average position for ties.

### Drug data

We retrieved data on 13,339 drugs from DrugBank (19) (downloaded July 2019) and selected only drugs that had been or currently FDA approved (n = 4,021), were indicated for use in humans (n = 1,964), and had at least one human protein target (n = 1,864). Of those, drug targets could be translated to human Entrez IDs for 1,844 drugs. The drug-target interactions used are provided in **File 1**. Sets of drugs that are approved for each disease were identified according to DrugBank’s (19) “Indication” category (**Supplementary Results**, **File 1-3 & 5**). Unless otherwise specified, precision is calculated using these disease-specific sets of FDA-approved drugs as *relevant drugs* or true positives.

For validation of ranked drug candidates, we also downloaded data from www.clinicaltrials.gov (September 2023). Data included information on 465,269 clinical trials registered from September 17^th,^ 1999, to September 7^th^, 2023. Clinical trials (n = 70,396) for 1,085 of the included 1,844 drugs were found. To derive information on the disease relevance of drug candidates, we filtered clinical trials further by MESH terms, resulting in sets of 724, 494, 532 and 140 drugs that had been tried for rheumatoid arthritis, multiple sclerosis, Crohn’s disease and psoriatic arthritis, respectively (**File 1-5**). Even though the outcome of such trials is largely unknown, using drugs registered for clinical trials alongside approved drugs for calculation of precision tests scDrugPrio’s ability to capture the pharmacological consensus of the medical community on drugs with an expected effect.

For further validation of drug ranking, we also performed a literature search for the top 100 ranked drug candidates of each data set. We systematically searched PubMed and Google Scholar between June 2020 and December 2022 using the specific disease denotation and the drug name as search terms. No restrictions or filters were applied. The relevance of the identified articles was screened by title and abstract. When no relevant articles were identified, the drug name was replaced by the substance name, and another search was conducted. To be eligible, studies had to 1) include a control group, 2) be a human clinical study or rodent experiment, 3) measure inflammatory activity and 4) be accessible. When several studies were identified that reported contradictory results, the drug was labelled as having a previously reported effect, reasoning that it would be impossible to determine the evidence and accuracy level in every such instance. In **Files 1-4**, a summary on the nature of the identified article and a full reference is provided, listed by drug.

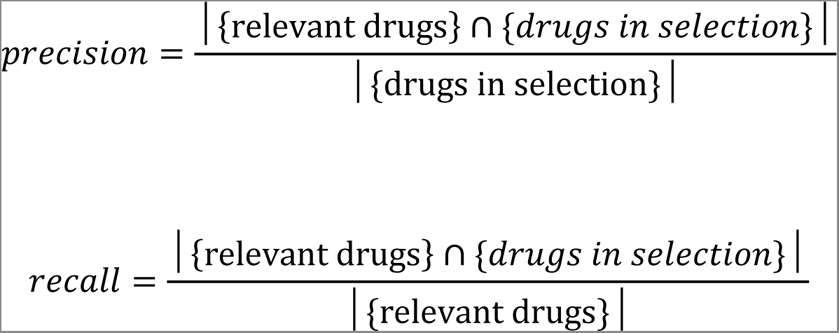

When referring to precision among the top 100 candidates, we refer to all candidates with rank ≤ 100.

### Protein‒protein interaction network

The human interactome was derived from do Valle et al.(51). The literature-curated interactome included 351,444 protein‒protein interactions (PPIs) connecting 17,706 unique proteins and was annotated using Entrez Gene IDs. The largest connected component included 351,393 PPIs and 17,651 proteins. Only the largest connected component was used for further analysis.

### Antigen-induced arthritis mouse model

Antigen-induced arthritis (AIA) was triggered in six 8-week-old, anaesthetised female 129/Sve mice by intra-articular injection of methylated bovine serum (mBSA) in the left knee joint after having presensitised mice to mBSA. The left knee joints of four naïve mice were injected with phosphate-buffered saline (PBS, 20 μL) and used as a negative control. One week after intra-articular triggering of AIA, mice were sacrificed, and joints were either used for immunohistochemistry or scRNA-seq. Histochemical preparation was performed as previously described (6), and specimens were examined in a blinded manner for pannus formation, cartilage and subchondral bone destruction, and synovial hypertrophy on an arbitrary scale, 0–3, as described by Magnusson et al. (52). For the scRNA-seq experiment, joint tissue was minced to ∼1 mm^3^ pieces, which were digested by collagenase Ⅳ (1.5 mg/mL) and DNase Ⅰ (100 µg/mL) at 37°C. Dissociated cells were passed through a 70-µm cell strainer. Single-cell suspensions were resuspended in RPMI-1640 at a density of 1 × 10^5^ cells/mL for cell loading. One mouse in which AIA had been triggered developed only mild arthritis (arthritis score 0.5) and was therefore excluded from further analysis. All experimental procedures were performed according to the guidelines provided by the Swedish Animal Welfare Act and approved by the Ethical Committee for Research on Animals in Stockholm, Sweden (N271-14).

scRNA sequencing was performed using the Seq-Well technique (53) following a described protocol (6). Briefly, prepared single-cell suspensions were counted and coloaded with barcoded and functionalised oligo-dT beads (Chemgenes, Wilmington, MA, USA, cat. No. MACOSKO-2011-10) on microwell arrays synthesised as described by Gierahn et al. (53). For each sample, 20,000 live cells were loaded per array to bind with oligo-dT beads. Beads were collected for capturing mRNA and preparing the library following cell lysis, hybridisation, reverse transcription and transcriptome amplification. Except for one library, which was sequenced alone, libraries from three samples were pooled for sequencing (**Table S2**), resulting in a coverage of 6.6 reads per base. Four libraries were prepared for each sample using the Nextera XT DNA Library Preparation Kit (Illumina, San Diego, CA, USA; cat. No. FC-131-1096) according to the manufacturer’s instructions. Each library was sequenced once, except for one library, which was sequenced twice using the NextSeq 500/550 system.

The single-cell data were processed into digital gene expression matrices following James Nemesh, McCarrol’s lab Drop-seq Core Computational Protocol (version 1.0.1, http://mccarrolllab.com) using *bcl2fastq* Conversion and Picard software. To increase the read depth for the cells, each sample was sequenced multiple times (**Table S2**), and the fastq files for each sample were merged before further alignment steps. The indexed reference for alignment of reads was generated from GRCm38 (June 2017, Ensembl) using STAR software (2.5.3). Only primary alignments towards the reference genome were considered during downstream analyses, according to the mapping quality using STAR software.

### Sampling and sequencing of psoriatic arthritis patients

#### Sampling

Psoriatic arthritis (PsA) patients and controls were recruited by the Immune-Mediated Inflammatory Diseases Consortium (IMIDC) (54). PsA patients were recruited from different rheumatology departments from university hospitals belonging to the IMIDC. All PsA patients were diagnosed according to the CASPAR diagnostic criteria for PsA (55) with > 1 year of disease evolution and > 18 years old at the time of recruitment. Exclusion criteria for PsA included the presence of any other form of inflammatory arthritis, rheumatoid factor levels greater than twice the normality threshold or confirmed presence of an inflammatory bowel disease. PBMCs were sampled prior to treatment with anti-TNF or anti-IL17 and cryopreserved. Treatment response was classified at week 12 according to the EULAR response (25) (**File 5**). For the anti-TNF study, 6 males and 10 females were included. The corresponding figures for anti-IL-17 treatment were 3 males (2 responders) and 13 females (6 responders). Simultaneously, healthy age- and sex-matched control subjects (**File 5**) were recruited from healthy volunteers recruited through the Vall d’Hebron University Hospital in Barcelona (Spain). All the controls were screened for the presence of any autoimmune disorder, as well as for first-degree family occurrence of autoimmune diseases. None were found to be positive. Four males and four females were included. The study was approved by the Hospital Universitari Vall d’Hebron Clinical Research Ethics Committee. Protocols were reviewed and approved by the local institutional review board of each participating center.

#### Cell thawing

PBMCs cryopreserved at -80°C were thawed in a 37°C water bath and transferred with a bored tip to a 15 ml Falcon tube containing 14 ml of 37°C prewarmed RPMI medium supplemented with 10% FBS (Thermo Fisher Scientific). Samples were centrifuged at 300x g for 10 min at (room temperature) RT, the supernatant was removed, and pellets were resuspended in 1 ml of 1X PBS (Thermo Fisher Scientific) supplemented with 1% BSA (PN 130-091-376, Miltenyi Biotec) and 10 µL of DNase I (PN LS002007, Worthington-Biochem). After incubation at RT for 10 min with periodic shaking, the cells were filtered with a 20 µm strainer (PN 43-10020-70, Cell Strainer) into a new 15 ml falcon on ice, and the filter was washed by adding 9 mL of cold 1× PBS. Samples were concentrated afterwards by centrifugation at 300 × g for 10 min at 4°C and resuspended in 1 × PBS with 0,05% BSA for further assessment of cell numbers and viability with the TC20™ Automated Cell Counter (Bio-Rad). Samples balanced by responders and nonresponders for each treatment were mixed in pools of 8 patients at a 50:50 ratio and concentrated by centrifugation in an appropriate volume of 1 × PBS-0.05% BSA to obtain a final cell concentration > 4,000 cells/µL, suitable for 10× Genomics scRNA-sequencing. The suspension was filtered again with a 20 µm strainer, and the cell concentration was verified by counting with the TC20™ Automated Cell Counter.

#### Cell encapsulation and library preparation

Cells were partitioned into Gel BeadInEmulsions (GEMs) by using the Chromium Controller system (10 × Genomics). Each pooled sample was loaded into two channels with a target recovery of 35,000 cells per channel to ensure a minimum final recovery of 2,000 cells per sample condition. After GEM-RT incubation, the resulting cDNAs were purified with SPRI beads. To ensure maximal cDNA recovery, a second Sylane bead purification was performed on the supernatant from the first purification, and both products were eluted together and preamplified for 13 cycles, following the 10 × Genomics protocol. cDNA was quantified on an Agilent Bioanalyzer High Sensitivity chip (Agilent Technologies), and 100 ng was used for library preparation. Gene Expression (GEX) libraries were indexed with 13 cycles of amplification using the Dual Index Plate TT Set A (10 × Genomics; PN-3000431). The size distribution and concentration of full-length GEX libraries were verified on an Agilent Bioanalyzer High Sensitivity chip. Finally, sequencing of GEX libraries was carried out on a NovaSeq 6000 sequencer (Illumina) using the following sequencing conditions: 28 bp (Read 1) + 10 bp (i7 index) + 10 bp (i5 index) + 90 bp (Read 2) to obtain approximately >20,000 paired-end reads per cell.

#### 3’ Single-cell RNA sequencing (scRNA-seq)

PBMC samples from 32 patients and 8 healthy controls were evenly mixed in pools of 8 donors per library following a multiplexing approach based on donor genotype, as in Kang et al. (56), for a more cost- and time-efficient strategy. Importantly, libraries were designed to pool samples together from the same treatment (anti-TNF or anti-IL17) but mixing patients with a different response to treatment. With this approach, we aimed to avoid technical artifacts that could mask subtle biological differences between responders and nonresponders. To profile the cellular transcriptome, we processed the sequencing reads with 10X Genomics Inc. software package CellRanger v6.1.1 and mapped them against the human GRCh38 reference genome.

#### Library demultiplexing

The donor’s genotypes (VCF format) were simplified by removing SNPs that were unannotated or located in the sexual Y, pseudoautosomal XY or mitochondrial chromosomes (chr 0, 24, 25 and 26, respectively). As genotypes were assembled using the human GRCh19 reference genome, we converted them to the same genome assembly used to map the sequencing reads, the human GRCh38 reference genome, using the USCS LiftOver (https://genome.ucsc.edu/cgi-bin/hgLiftOver) command line executable. To meet the LiftOver required format (BED format), we used an available wrapper script (liftOver_vcf.py) to support input/output from VCF format (57). The library demultiplexing by donor was performed with cellsnp-lite v1.2.2 in Mode 1a (57), which allows genotyping single-cell GEX libraries by piling-up the expressed alleles based on a list of given SNPs. To do so, we used a list of 7.4 million common SNPs in the human population (MAF > 5%) published by the 10,00 Genome Project consortium and compiled by Huang et al. (57). Importantly, we used the default parameters, setting the MAF > 5% (--minMAF 0.05) and requesting genotyping in addition to counting (--genotype). Then, we performed donor deconvolution with vireo v0.5.6 (58), which assigns the deconvoluted samples to their donor identity using known genotypes while detecting doublets and unassigned cells. Finally, we discarded detected doublets and unassigned cells before moving on to the downstream processing steps.

### scRNA-seq data sets and preprocessing

Below, we describe all of the scRNA-seq data sets and the preprocessing steps of the current work as outlined in **Fig. S1**. As most preprocessing steps were applied to scRNA-seq expression matrices of all data sets, we will describe them jointly.

#### Quality cut-offs

Starting from a raw scRNA-seq gene expression matrix, the quality of cells was assured by application of quality cut-offs that aimed to filter out low-quality cells (few genes, low read depth), dying cells (high expression of mitochondrial genes) and doublets (unexpectedly high reads and large number of genes). While arbitrary, these specific cut-offs were adapted to the corresponding data set and reported in **File 1-5** (59). Genes that were expressed in less than three cells were excluded from further analysis.

#### Batch correction

In case a high degree of interindividual expression differences existed, batch correction was performed according to a previously established pipeline (60). In short, we used Seurat’s function *findIntegrationAnchors()* (61) for the list of objects that corresponded to each individual. These anchors were later used by *IntegrateData()* (61) to integrate the data from individuals to correct for patient-specific differences as suggested in (62).

#### Denoising

Next, data for all cells were processed by a deep count autoencoder (DCA) model (18), which is a neural network performing a nonlinear principal component analysis (PCA). The DCA method is initiated by computing a library size, log- and z-score normalised expression matrix, which is taken as an input to the neural network, and the output of the neural network log10 transformed), and denoised single-cell expression matrix ***D***, which has the same features as the original data but is corrected for various sources of noise in the data. The DCA method also outputs a representation of the original single-cell data in a latent space. This representation has many fewer features than the original data, which is particularly important for performing accurate cluster analyses. The intercellular expression differences are generally better represented in this latent space than in purely linear PCA models, and the latent space representation is also corrected for single-cell data artefacts such as dropouts and varying library sizes.

*Clustering analysis* was performed using the Seurat v3.1 package (61) on the DCA-derived latent representation. A shared nearest neighbor graph was constructed, and neighborhood overlap between every cell and its *k*-nearest neighbors was calculated based on the Jaccard index using the *FindNeighbors()* function on all supplied latent features. Next, clusters were identified through application of the Louvain algorithm to the shared nearest neighbor graph using the *FindClusters()* function along with a specified resolution setting. The resolution parameter and *k* for *k-*nearest neighbor analysis were tailored to each data set and are reported below. Clusters were visualised through *RunTSNE ()*.

#### Analyses of interindividual molecular heterogeneity

After denoising and clustering, heterogeneity among samples was determined. For this, we trained a flexible machine learning model that attempts to find a decision boundary between the given groups of cells. If this model results in a high misclassification rate for test data, it indicates that groups are highly mixed. More specifically, the data were randomly divided in half and used for training and testing the model. A random forest classifier (63) was used to classify cells from sick samples based on what patient the sample was derived from. Cross-validation with 10-fold and grid search (64) was used to find the most appropriate hyperparameters of the random forest. The bootstrap (65)percentile method (65) was used to construct the 95% confidence intervals for training and test misclassification rates. The Scikit package (66) from Python (3.7.9) was used to perform the analysis.

Furthermore, patient heterogeneity was explored through comparison of cell type proportions and examination of latent features of the non-batch-corrected data. Interindividual differences in cell type proportions were explored by the application of the chi-square test to the proportions of cell types in sick samples. Latent feature comparison was conducted visually through tSNE visualisation of the latent features of each patient. The results for these analyses are found in the **Supplementary Results**.

#### Cell typing

While cell typing is not crucial for scDrugPrio (which might be performed on unlabelled clusters), we cell typed clusters to enhance biological interpretation. Cell types were assigned to each cluster based on the relative coexpression of several known cell type marker genes. For AIA data, each gene’s expression was expressed as a fraction of a cell’s total gene expression score. For visualisation of gene expression differences between clusters, z-scores were calculated. Z-scores for single cells were derived by comparison of one cell’s gene fraction to all other cells’ gene fractions. Z-scores for gene expression of clusters were derived by comparison of the average gene fraction in a cluster to the cluster-averaged gene fractions of cells in other clusters. Murine cell type-specific marker genes for the RA data were derived from the online resources of the R&D systems (www.rndsystems.com/research-area; accessed July 2020). Cell typing of the human data sets was performed using DCA denoised gene fractions and utilised combinations of marker genes (**File 2 & 3**).

#### Differentially expressed genes

For each cell type separately, differentially expressed genes (DEGs) were calculated by comparing denoised gene expression of cells derived from healthy samples vs. cells from sick samples. For this purpose, the *FindMarkers()* function in Seurat (61) was used to deploy a scRNA-seq-tailored hurdle model supplied by the MAST package (67). Genes were considered significantly differentially expressed when they showed an absolute log fold change greater than or equal to 1.5 and a Bonferroni-adjusted *P* < 0.05. The fold change cut-off was motivated by previous studies (68) and aimed to decrease the number of DEGs for later network calculations.

#### Antigen-induced arthritis

Quality cut-offs resulted in a total of 16,751 cells (**File 1**). Genes were annotated as murine NCBI Gene Symbols. Following denoising, clustering using k = 20 and a resolution of 0.6 resulted in the identification of 20 clusters that were cell typed. Heterogeneity analysis used n_estimators = 1,500, max_depth = 15, min_samples_split = 50, and min_samples_leaf = 50 and showed no significant heterogeneity. DEGs and the denoised expression matrix were translated to human Entrez gene IDs using human-mouse orthologues downloaded from NCBI (August 2019).

#### Multiple sclerosis

A unique molecular identifier (UMI) matrix (22) for cerebrospinal fluid (CSF) of five human multiple sclerosis (MS) patients and five human patients with idiopathic intracerebral hypertension (IIH) were downloaded from the Gene Expression Omnibus (GEO) database (GSE138266). Gene annotation was translated from human Ensembl gene IDs to human Entrez gene IDs and symbols using the HUGO Gene Nomenclature Committee (HGNC) database (69) (downloaded November 2020). After the application of quality cut-offs (**File 2**), we derived 33,848 cells. Initial preprocessing was performed without batch correction using cluster parameters *k* = 10 and resolution = 0.2 after DCA denoising to derive 17 clusters. Interindividual heterogeneity was assessed as described below using the following hyperparameters: n_estimators = 500, max_depth = 30, min_samples_split = 50, min_samples_leaf = 25. Since we noticed substantial patient-related heterogeneity (**Fig. S5a**), preprocessing was repeated, including batch correction, DCA denoising, and clustering using k = 15 and resolution = 0.35 to derive 21 clusters that were cell typed using known marker genes derived from the original publication (22) (**File 2**). The number of DEGs ranged from 0 to 10,076.

#### Crohn’s disease

A unique molecular identifier (UMI) matrix (24) for eleven human Crohn’s disease (CD) patients was downloaded from GEO (GSE134809). Data for each patient included intestinal biopsies from one inflamed site and one uninflamed site. After application of quality cut-offs (**File 3**), we derived 77,416 cells. Gene annotation was translated from human Ensembl gene IDs to human Entrez gene IDs and symbols using the HGNC database (2020-11-08) (69). Initially, data were DCA denoised without applying batch correction. Clustering was performed using *k* = 15 and resolution = 0.8. Interindividual molecular heterogeneity was assessed as described below using the following hyperparameters: n_estimators = 1,000, max_depth = 20, min_samples_split = 50, min_samples_leaf = 25. Since there was substantial interindividual heterogeneity (**Fig. S8a**), preprocessing was repeated now batch-correcting before DCA denoising. Clustering was again performed using *k* = 15 and resolution = 0.8.

#### Individual Crohn’s patients

For individual patient predictions, we used the same quality cut-offs as for the pooled analysis of CD patients. As interindividual heterogeneity does not affect the predictions made for individual patients, these calculations were performed on non-batch-corrected data. DCA denoising was applied to the joint data, and gene annotation was translated to Entrez gene IDs. Thereafter, scRNA-seq data were separated by patient, and cells from each patient were clustered individually. An individual patient cluster was assigned a cell type based on which cluster it most resembled in the joined CD analysis, as measured by the number of shared cell identifiers. DEGs were then calculated between cells from sick and healthy samples. Visualisations of data were in part created using <u>BioRender.com</u>.

#### Psoriatic arthritis

Data sets were divided into one anti-TNF and one anti-IL17 data set, including responders (R), nonresponders (NR) and healthy controls. We filtered out the doublet and unassigned cells as well as those that did not meet the quality cut-off criteria (**File 5**) and derived 78,610 cells with 5,088 mean reads for the anti-TNF data set and 72,472 cells with 5,343 mean reads for the anti-IL17 data set. In both data sets, 19,415 cells were derived from healthy controls. Data sets were batch-corrected, and DCA was performed. For anti-TNF, clustering was performed using *k* = 25 and resolution = 0.25. The corresponding parameters for anti-IL17 were *k* = 15 and resolution = 0.45. The remaining downstream analysis was performed for responders and nonresponders separately, which meant that DEGs were calculated between responders and healthy controls and between nonresponders and healthy controls. For the anti-TNF data set, the number of DEGs ranged from 0 to 5,989 for R and from 0 to 5,877 for NR. The corresponding figures for anti-IL17 were 0 to 3,097 and 0 to 3,284. Next, scDrugPrio was applied to DEGS from anti-TNF R and NR as well as anti-IL17 R and NR separately.

### In vitro validation of potential novel drugs

To validate the predicted novel drugs, *in vitro* culture of murine and human B cells upon activation with the indicated stimuli was employed to assess the effects of the predicted drugs on B-cell survival, activation, proliferation, and antibody production. Three doses for each predicted drug were used to challenge *in vitro* cultured B cells (**Table S1**). For the assessment of potential novel drugs on murine B-cell survival and activation, 300,000 murine naïve B cells (Lin^-^B220^+^CD43^-^) were enriched by flow cytometric sorting and cultured in the presence of AffiniPure F(ab’)₂ Fragment goat anti-mouse anti-IgM (10 μg/mL, CAT: 115-006-075, Jackson ImmunoResearch), anti-mouse CD40 (10 μg/mL, Clone:1C10, Biolegend), or LPS (10 μg/mL) for 24 hours. B-cell survival was determined by flow cytometric analysis of propidium iodide (PI)^+^ cells. Surface CD69, CD86, and MHC-II were used as readouts for assaying B-cell activation. For the analysis of B-cell proliferation, purified B cells were stained with carboxyfluorescein succinimidyl ester (CFSE) (1 μM) before *in vitro* culture for three days. To determine the effects of the predicted drugs on antibody production, 200,000 purified murine naïve B cells were stimulated with anti-CD40 (10 μg/mL) + IL-4 (10 ng/mL), LPS (10 μg/mL) + IL-4 (10 ng/mL), or LPS (10 μg/mL) + IFN-γ (10 ng/mL) for six days.

For the analysis of novel drugs predicted to regulate the biology of human B cells, peripheral blood mononuclear cells (PBMCs) were isolated from the buffy coat as previously described (70, 71). Human naïve B cells were subsequently preenriched by MACS sorting using a B-Cell Isolation Kit II (Miltenyi) and further purified by flow cytometric sorting of CD19^+^CD27^-^ B cells. Purified human naïve B cells were cultured in 96-well plates in the presence of AffiniPure F(ab’)₂ Fragment goat anti-human IgG + IgM (5 μg/mL, CAT: 109-006-127, Jackson ImmunoResearch), anti-human CD40 (5 μg/mL, Clone: G28.5, Bio X Cell), and IL-21 (10 ng/mL, PeproTech).

B-cell survival was determined by flow cytometric analysis of propidium iodide (PI)+-cells. Surface CD69, CD86, and MHC-II were used as readouts for assaying murine B-cell activation. Surface CD69 was assayed for the measurement of human B-cell activation. For the analysis of B-cell proliferation, purified B cells were prestained with carboxyfluorescein succinimidyl ester (CFSE) (1 μM) before *in vitro* culture for three days. Murine IgG2a and IgG1, as well as human IgG in the supernatant, were determined by enzyme-linked immunosorbent assay (ELISA) using goat anti-mouse Ig, goat-anti-mouse IgG1-HRP and goat-anti-human IgG2a-HRP, goat anti-human Ig, and goat-anti-human IgG-HRP (SouthernBiotech) as previously described (72).

### In vivo validation of predicted drugs

Amrinone was tested for treating collagen-induced arthritis (CIA). For this, male DBA1/J mice purchased from GemPharmatech (China) were immunised intradermally with 100 μg of chicken type II collagen (2 mg/mL, Chondrex, USA) emulsified with complete Freund’s adjuvant (CFA, 1 mg/mL) and boosted on day 21 with 100 μg of chicken type II collagen emulsified with incomplete Freund’s adjuvant (IFA). Mice were i.g. given with diluent (n = 5) or amrinone (30 mg/kg, n = 5) daily from day 21 for 3 weeks. The rear paw thickness and the clinical arthritis score for each limb were recorded every other day from 0 to 4 with a maximal score of 16 for each mouse according to the previous protocol (73). Mice were maintained in a specific pathogen-free animal facility at Xuzhou Medical University, and all animal studies were performed in accordance with protocols approved by the Animal Experimental Ethics Committee of Xuzhou Medical University (202012A162).

Mice were sacrificed on day 21 post drug intervention. Serum was collected for the analysis of collagen-specific autoantibodies by enzyme-linked immunosorbent assay (ELISA) as previously described (74). Briefly, diluted serum was incubated in a 96-well ELISA plate precoated with chicken type II collagen (5 μg/mL). Goat-anti-mouse IgG1-HRP, goat-anti-mouse IgG2a-HRP, and goat-anti-mouse IgG-HRP (SouthernBiotech) were used as detection antibodies. Knee joints were fixed in 4% formaldehyde and subsequently decalcified with decalcification solution (ServiceBio, China) for one week. The specimens were next embedded in paraffin, and sagittal sections (4 µm) were cut. The sections were stained with hematoxylin and eosin (H&E) for the histological analysis of immune cell infiltration and Safranin-O for the analysis of bone erosion as previously described (72, 74).

### Comparison of scRNA-seq-based screening outcomes for rheumatoid arthritis to other data types and prediction methods

Briefly, we benchmarked outcomes based on the scRNA-seq-derived DEGs against microarray data (GSE55235 & GSE93272) (75, 76), GWAS Catalog (77) genes and OMIM (78) genes as well as combinations of these data sets. scDrugPrio was compared to previous methods such as 1) identifying druggable DEGs, targeting key enriched pathways (79), CMAP (33) drug predictions and the empirical drug selection of Kim et al. (14). Predictions were also replicated using the smaller, unbiased HuRI PPIN (80) (8,236 proteins, 52,150 interactions) to ensure the absence of knowledge bias. More information can be found in the **Supplementary Results**.

### Data and code availability

scRNA-seq data that support the findings of this study are openly available at Gene Expression Omnibus (GEO), reference number GSE193536. Unless otherwise stated, analysis was performed in R 3.6.3. The code for data cleaning and analysis associated with the current submission is available at https://github.com/SDTC-CPMed/scDrugPrio.

