## Supplemental information for "scDrugPrio: A framework for the analysis of single-cell transcriptomics to address multiple problems in precision medicine in immune-mediated inflammatory diseases"

[1]Centre for Personalised Medicine, Linköping University; Linköping, Sweden.
[2]Department of Gastroenterology and Hepatology, University Hospital, Linköping, Sweden.
[3]Division of ENT, CLINTEC, Karolinska Institute, Stockholm, Sweden.
[4]Division of Statistics and Machine Learning, Department of Computer and Information Science, Linkoping University; Linköping, Sweden.
[5]Mavatar, Inc., Stockholm. Sweden
[6]Division for Bioinformatics, Department of Physics, Chemistry and Biology, Linköping University; Linköping, Sweden.
[7]CNAG-CRG, Centre for Genomic Regulation (CRG), Barcelona Institute of Science and Technology (BIST), 08028 Barcelona, Spain
[8]Universitat Pompeu Fabra (UPF), 08002 Barcelona, Spain
[9]Grup de Recerca de Reumatologia, Institut de Recerca Vall d'Hebron, Barcelona, España
[10]Department of Physics and Astronomy, Northwestern University, Evanston, IL 60208, USA.
[11]Northwestern Institute on Complex Systems, Northwestern University, Evanston, IL 60208, USA.
[12]Division of Cardiovascular Medicine, Channing Division of Network Medicine, Department of Medicine, Brigham and Women's Hospital, Harvard Medical School; Boston, MA, USA.
[13]Jiangsu Key Laboratory of Immunity and Metabolism, Department of Pathogenic Biology and Immunology, Xuzhou Medical University; Jiangsu, China.
[¤]Joint last authors.

#### Supplementary Results

*Identification of known disease-drug pairs based on DrugBank*

Drug-disease associations were identified based on the "Indication" category in DrugBank. RA drugs were indicated for use in "Rheumatoid arthritis"/"RA". Drugs for Crohn's Disease were indicated for use in "Crohn's Disease", "IBD" and "Inflammatory Bowel Disease". Drugs for MS were selected based on the following DrugBank indications: "Active secondary progressive Multiple Sclerosis", "Progressive Multiple Sclerosis (PMS)", "Relapsing Remitting Multiple Sclerosis (RRMS)", "Progressive Relapsing Multiple Sclerosis", "Multiple sclerosis exacerbation", "Multiple Sclerosis, Primary Progressive", "Acute Multiple sclerosis", and "Relapsing Multiple Sclerosis (RMS)". PsA drugs were identified indicated for use on "Active Psoriatic arthritis", "Active Psoriatic arthritis", "Severe Psoriatic Arthritis", and "Psoriatic Arthritis".

*The biological importance of the multicellular disease model of antigen-induced arthritis is supported by enrichment analyses*

Since genetic variants have previously been associated with relevant disease mechanisms (1), gene set enrichment analysis was performed to evaluate the relevance of transcriptome-defined DEGs. Hence, for each cell type, a gene set enrichment was performed. To assess the relevance of the MCDM, we correlated eigenvector centralities of cell types in the MCDM and enrichment significance of GWAS gene enrichment among cell type-specific DEGs.

For this, the GWAS Catalog (2) version 1.0.2 was downloaded (January 2019), and GWAS genes were extracted for the following traits: "Celiac disease or Rheumatoid arthritis", "Rheumatoid arthritis”, “Rheumatoid arthritis (ACPA-negative)”, “Rheumatoid arthritis (ACPA-positive)”, and “Rheumatoid arthritis (rheumatoid factor and/or anti-cyclic citrullinated peptide seropositive)”. Intergenic SNPs were mapped towards the closest gene. Only GWAS genes with a genome-wide significance of $P < 10^{-8}$ were considered. GWAS gene enrichment among DEGs was calculated for each cluster separately with Fisher’s exact test using genes with Ea ≥ 0.2 (Eq. 1) as background genes. We found that intercellular cell type centrality correlated with GWAS enrichment significance.

The relevance of the inferred interactions in the MCDM were further supported by gene set enrichment analysis, which showed that known drug targets for human RA were significantly ($P < 0.05$) enriched among the ligands with the collectively most impactful interactions (as measured by summed Pearson correlation). Furthermore, we found that targets for US Food and Drug Administration (FDA)-approved RA drugs were enriched among DEGs of central cell types, such as immature and activated B cells (FDR adjusted $P < 0.01$ respectively; **Fig. S2d-g**). Additionally, we performed a preranked gene set enrichment analysis (51) with the targets of RA drugs and the list of ligands ranked by the summed Pearson coefficient of all cell type interactions they were involved in as the input. We used 100,000 permutations, seed 149 and otherwise default parameters and found that RA drug targets were significantly ($P < 0.05$) enriched among the most impactful ligands (as measured by summed Pearson correlation).

*Drug network degree supports intracellular centrality as a ranking measure*

To test whether network centrality could be a potential proxy for drug effects, we examined the network degrees of all drugs included in DrugBank, as well as the degree of drugs that were approved for inflammatory diseases. For this analysis, we calculated the network degree of all genes included in the LCC of the literature-derived PPIN using the *degree()* function included in the igraph package (3). Next, we summarized the network degrees of drug targets. As many drugs had several targets, which complicated comparison to the background PPIN, we calculated 1) the max network degree of all targeted proteins, 2) the mean network degree, and 3) the sum of all targeted proteins network degrees (**Fig. S17**). Drugs generally tended to target higher network degree targets. This observation became especially apparent when examining the network degree of drugs approved for the inflammatory diseases included in this study. Although antibody drugs target only a few proteins, they still tend to target proteins with higher network degrees. This is an interesting finding, as it could guide new drug target discovery strategies as well as spark additional research efforts aiming to examine why low network degree targets are underrepresented.

*Intercellular and intracellular centrality for drug ranking*

To evaluate intercellular centrality as a ranking measure in AIA, we performed a permutation test (1,000 iterations) that showed that the mean rank of established RA drugs was significantly better than the random expectation obtained from random permutations of intercellular centrality mean ranks (88.03 vs 168.07 ± 16.44; one-sided $P < 10^{-6)}$.

When ranking drugs only by intracellular centrality, the mean rank of established RA drugs was significantly better than random expectation (110.55 vs 167.20 ± 16.22; one-sided $P < 10^{-3}$), similar to the analysis of intercellular centrality above. To validate whether ranking based on both inter- and intracellular centrality held additional value, drugs were ranked primarily by combined intercellular centrality and second by intracellular centrality. When comparing the mean rank of

RA drugs in this ranking to the random expectation obtained from 1,000 iterations in which drugs were ranked by intercellular centrality scores and randomly permuted intracellular centrality scores, we found that additional ordering according to mean intracellular centrality did not result in significantly better or worse ranking; however, we found that it acted as a tiebreaker for drugs achieving the same intercellular centrality-derived rank and increased precision for literature drugs among the top ranking drugs. We hence use a composite score derived from inter- and intracellular centrality for ranking.

*Comparison of eigenvector centrality to other centralities*

In the context of intercellular centrality, we calculated different centrality algorithms. The results can be found in the supplementary data of the respective data set (**File 1-5**). Different centrality approaches (e.g., eigenvector centralities, K-core decomposition, Kleinberg's hub centrality scores, Laplacian centrality, leverage centrality, group centrality) all rank the centrality of cell types similarly. This is exemplified by a high correlation between cell type eigenvector centrality scores and K-core decomposition (Pearson $r = 0.77$; $P < 10^{-3}$), Kleinberg's hub centrality scores ($r = 0.72$; $P < 0.01$), Laplacian centrality ($r = 0.72$; $P < 0.01$), leverage centrality ($r = 0.85$; $P < 10^{-4}$), and group centrality ($r = 0.77$; $P < 10^{-3}$). This comparison was performed in the AIA data set.

To test whether drug ranking using other centrality algorithms yielded similar results, we tested the above centrality algorithms that had yielded similar intercellular centralities. Comparison of results was based on Pearson correlation between drug ranks of eigenvector centrality-based results to that of drug ranks derived by other algorithms. Out of the above, we found that K-core decomposition ($r = 0.80$; $P < 10^{-300}$), Kleinberg's hub centrality scores ($r = 0.86$; $P < 10^{-300}$), Laplacian centrality

($r$ = 0.86; P < $10^{-300}$), and group centrality ($r$ = 0.89; P < $10^{-300}$) produced similar drug rankings (**File 1**). This comparison was performed in the AIA data set.

*Selection of the number of topmost significant DEGs used for network proximity calculation*

As drug prioritization was dependent on the significance of network proximity (derived by bootstrap), high numbers of DEGs relative to the size of the network (literature PPIN LCC includes 17,651 unique proteins) result in $P(z_c) > 0.05$ for most drugs. Hence, a significance threshold was introduced to each data set individually that specified the number of topmost significant DEGs used in proximity calculations. To investigate where this threshold should be set, proximity calculations were performed over a range of different cut-offs (from a minimum of 100 DEGs up to all possible DEGs). The chosen threshold aimed to maximize precision and recall while simultaneously being consistent within a range of tried cut-offs. The network proximity calculations for MS patients and pooled CD patients were limited to the top 1,800 significant DEGs of each cell type. As these calculations consume immense computational resources and had initially been performed on non-batch-corrected data, batch-corrected data inherited their thresholds. For individual CD patient network proximity calculations, the threshold was set to the top 3,500 significant DEGs. For the PBMC data sets of PsA anti-TNF R and NR as well as PsA anti-IL17 R and NR, cut-offs were calculated on the batch-corrected data before network predictions were limited to the 3,000 top significant DEGs.

*Validation of scDrugPrio compared to direct drug target identification and network predictions based on other disease genes.*

To investigate the feasibility of developing a drug prioritization framework based on single-cell derived DEGs, we first tested whether the identified DEGs carried similar amounts of information as other available gene sets, such as genetic variations (from GWAS Catalog (4) and OMIM (5)), microarray-derived (GSE55235 & GSE93272) (4) DEGs for rheumatoid arthritis, and genes for which cell type single-cell expression values correlated with clinical disease severity scores of the mice (referred to as correlated genes).

Furthermore, we evaluated different denoising strategies on the gene sets of interest aiming to condense and filter the gene set to derive better precision and recall for known RA drugs among the predicted drug candidates. Such denoising strategies included the use of 1) overlapping genes between gene sets (e.g., DEGs found in more than one cell type might arguably carry a higher predictive value), 2) the largest connected component (LCC) of a gene set, 3) Fisher pathway enrichment analysis using KEGG pathways (6), 4) the topmost significant genes in a gene set, and 5) combinations of these.

Drug candidate selection for the gene set of interest was attempted by 1) selecting all drugs that targeted at least one gene in the gene set of interest or by 2) applying the previously described network proximity-based approach (7) for which a cut-off of $z_c < -1.64$ was used (**Fig. S18–S19**). All of the above predictions were repeated in the smaller, unbiased HuRI PPIN (8) (8236 proteins, 52150 interactions) to ensure the absence of knowledge bias (**Fig. S20**).

Lastly, we benchmarked our approach to a preexisting approach (Connectivity-Map) developed for use on bulk RNA-seq data (**Fig. S21**) and compared our results to results from a previous case report by Kim et al. in which scRNA-seq had been used to inform empirical drug choice for a patient who did not respond to standard treatment.

*OMIM genes.* We downloaded genes for "Rheumatoid arthritis" from the Online Mendelian Inheritance in Man (OMIM) database (5) (June 2019).

*Microarray-derived DEGs.* Microarray data for synovial tissue were downloaded from GEO (GSE55235) (9) and included data for healthy controls and RA patients. For further analysis, we selected the synovial tissue samples of treated RA patients (n = 10) derived during joint replacement/synovectomy, as well as control samples that were obtained from postmortem joints (n = 10) of healthy individuals. Microarray data for whole blood samples of drug-naïve RA patients (n = 30) and healthy individuals (n = 30) were downloaded from GEO (GSE93272) (9, 10).

Data were downloaded utilizing the *getGEO()* function in the GEOquery R package (11). Microarray probes were translated to human Entrez Gene IDs utilizing the respective annotation files supplied by Affymetrix. If a probe ID corresponded to several Entrez Gene IDs, the probe was excluded from further analysis. If several probe IDs corresponded to the same Entrez Gene ID, only the probe ID with the highest fold change between sick and healthy samples was retained. DEG calculation was based on the expression values of the included samples and was conducted using the limma R package (12). First, a linear model for each gene's expression in all independent samples was fitted using the *lmFit()* function. Next, the *eBayes()* function is used to calculate the moderated t-statistics, moderated F-statistics and log-odds of differential expression by empirical Bayes moderation of the standard errors towards a common value. P values were adjusted using the Benjamini–Hochberg false discovery rate (FDR). Only DEGs with FDR-adjusted P values < 0.05 were considered for further analysis (**File 1**).

*Genes correlated with arthritis severity score ("Correlated genes").* For every cell type, we divided cells by mouse of origin and calculated the mean gene expression for every gene. Next, we correlated the mean gene expression of a cell type with the paired arthritis severity score using a Pearson correlation. To investigate whether the correlation was stronger than expected by chance, a bootstrap algorithm was employed that randomly deviated cell type and mouse-specific mean expression values by -0.2 to 0.2. P values for every gene were derived by comparison of the gene's correlation coefficient to the bootstrap mean correlation coefficient and standard deviation thereof. Only genes with $P < 0.05$ were considered for further analysis.

*Largest connected component*. The largest connected component (LCC) of a disease-associated gene set in a given PPIN was identified through application of the *induced_subgraph()* function of the igraph R package *(3)*. The largest such subgraph was selected for further analysis.

*Pathway enrichment analysis using KEGG pathways.* Human KEGG pathways were downloaded from KEGG (6) (January 2020) using the KEGGREST R package. A Fisher enrichment analysis was conducted to identify which pathways were significantly enriched with a specific gene set of interest. P values for all pathway enrichments were FDR adjusted. Pathway genes with P < 0.05 were considered gene sets of interest in further analysis.

*Valdiation in the HuRI protein–protein interaction network.* For validation, initial network outcomes were replicated using the smaller yet unbiased HuRI PPIN (8). Network proximity calculations were repeated in the same manner as previously described. Genes that were not included in the HuRI PPIN were removed from the list of drug targets as well as the gene sets of interest prior to calculation. As a result, the calculation included 888 drugs (compared to 1,840 in the literature-curated PPIN calculations), of which 30 were known RA drugs. Generally, recall and precision were slightly lower among the candidates in the HuRI-based network proximity screening (**Fig. S20**) but were otherwise comparable to results in the literature-curated PPIN (**Fig. S18 & S19**).

*Pseudobulk RNA-seq data were used as input for Connectivity-Map.* Since there are no alternative systematic methods that compute drug prioritization based on single-cell data, we compare our method with CMAP (13), which outputs a drug ranking by using signatures of bulk data and drug targets associated with these signatures. First, we converted healthy and sick scRNA-seq samples from AIA mice into pseudobulk expression by using *AggregateExpression*() in Seurat (14). Cells from each mouse were sampled twice to create two bulk profiles per mouse to be able to reach significance in later DEG calculations. Next, we computed DEGs between healthy and sick pseudobulk samples using the limma package as described above (12), which resulted in 1,105 and 23 genes that were up- and downregulated, respectively. The above approach ensured maximum comparability between methods, given that the transformation of scRNA-seq data to bulk data allows the same data set to be used for comparison of scDrugPrio and CMAP.

For its computations, the CMAP method needs a database containing information about genes that are targeted by each drug in the repository. To make a relevant comparison between CMAP and scDrugPrio, we used the 1840 DrugBank-derived drugs that entered network analysis in scDrugPrio as such a database. We sorted significant DEGs based on absolute log-fold change before inputting the by CMAP required 150 top DEGs.

CMAP returned a ranked list including all 1840 drugs; however, only 95 drugs had a predicted CMAP effect > 0. Five approved RA drugs were found among these 95 drugs (**Fig. S20)**. After the literature search had been performed (similar to the literature search performed for the AIA data), 20 additional drugs were suggested to have an effect, and 10 drugs had been studied previously and were either found to have no effect or exacerbating effects on disease. The precision for approved RA drugs was 5.3% (compared to 22% for scDrugPrio) and improved to 26.3% following the literature search (compared to 62% for scDrugPrio). Of the drugs that had been previously tested, 71.4% had shown success (compared to 95.4% for scDrugPrio).

*Robustness analysis of scDrugPrio*

To evaluate parameter dependency and thereby repeatability and reproducibility, we applied scDrugPrio to the AIA data set varying set cut-offs (**Fig. S22**). Such cut-offs included 1) the number of DEGs used for network calculations, including network proximity and intracellular centrality calculations, 2) the significance of $z_c$ (set to one-sided $P < 0.05$), 3) the network distance cut-off ($d_c < 1$), 4) pharmacological filtering criteria of a drug having to counteract the fold change of at least one targeted DEG, and 5) the background gene cut-off used during NicheNet ligand activity analysis for creation of the MCDM. The biopharmacological criteria (4) required manual evaluation, which complicates permutation. However, permutation was also deemed unnecessary, given that the biopharmacological criteria, applied to the same underlying data, would be expected to select/exclude the same set of drug candidates in every run.

To test the robustness of scDrugPrio to the other set cut-offs, we selected and ranked drug candidates over a variety of cut-offs. For the number of DEGs, no cut-off was originally set for the AIA data, resulting in up to 12,769 DEGs in the network calculations. Four more DEG cut-offs were chosen through random sampling within a preset range from 8,000 DEGs to the original cut-off. Following random sampling of the number of DEGs, network proximity calculations were performed using the original cut-off as well as 12,558, 12,148, 10,561 and 8,921 DEGs. The same numbers of DEGs were used for intracellular centrality calculations. Drug selection was conducted using all possible combinations of $d_c < 0.8$, 0.9, 1.0, 1.1 or 1.2 and $P(z_c) < 0.01$, 0.03, 0.05, 0.07, 0.09, resulting in a total of 25 combinations of $z_c$-derived P values and $d_c$. The original background gene cut-off was $Ea(i) \geq 0.2$. We calculated NicheNet-derived intercellular centralities for a range of cut-offs spanning from 0.10 to 0.28 using 0.02 increments. Using all possible cut-off combinations for 1-3) and 5), we derived 1250 different sets of ranked drug candidates.

Aiming to evaluate robustness, the ranks of drug candidates were compared to the original drug ranking using Pearson correlation. A Pearson correlation coefficient of 1 would represent a perfect replication of the original drug ranking despite changed cut-offs. Our analysis showed a median [min – max] Pearson correlation coefficient of 0.861 [0.704 – 0.999], indicating that scDrugPrio was a stable and relocatable performance over the chosen threshold ranges (**Fig. S22**). The number of DEGs seemed to affect the correlation coefficient the most, followed by the $z_c$ cut-off. P values were Bonferroni corrected and remained significant in all instances (**Fig. S22**).

*Interindividual heterogeneity among sick patients/samples*

To explore the variation between patients/samples, we compared 1) cell type proportions, 2) gene expression profiles and 3) examination of latent features of the non-batch corrected data. Interindividual differences in cell type proportions were explored by the application of the chi-square test to the proportions in patient samples. Heterogeneity between patients' gene expression profiles was explored through a random forest approach as described in the Methods section. Latent feature comparison was conducted visually through tSNE visualization of sick cell latent features derived from DCA (15). Cells were coloured based on the patient/sample to which they belonged. This allows for a visual appreciation of general molecular differences between patients. As seen, latent features for AIA mice show that mice mix well (**Fig. S2c**), speaking to no underlying differences in their gene expression profiles. However, MS (**Fig. S5a,b**) and CD patients (**Fig. S8a,b**) show a more nonoverlapping pattern, indicating heterogeneity.

To understand whether heterogeneity between patient samples could be due to clinical factors, such as sex and disease progression, we downloaded available clinical data for the MS patients. We performed a correlation analysis between continuous clinical variables and cell type proportions of individual subjects (**Fig. S22**). Analysing patient heterogeneity, we derived cell type proportions from the clustering of the non-batch-corrected MS data. The clinical variables included age as well as standard CSF parameters of all included patients, namely, total cells per µl, granulocytes per µl, red blood cells per µl, lymphocytes per µl, glucose (mg/d), lactate (mmol/l), and protein (mg/l).

From these parameters, granulocytes and red blood cell counts were excluded because they did not show any meaningful variation between patients.

We tested the dependency between the cell type proportions of each of the IIH and MS subjects and their associated clinical data. We calculated Spearman correlations to identify if any of the cell type proportions significantly correlated with a continuous clinical variable (**Fig. S22a**, **File 2**). A significant correlation was found between:

1. Age and
    a. Activated terminally differentiated effector CD8+ T cells (P = 0.025, RHO = 0.64)
2. Total number of cells per µl and:
    a. Naïve CD4+ T cells (P = 0.04, RHO = -0.63),
    b. Early activated CD4+ memory T cells (P = 0.02, RHO = 0.69),
    c. Activated terminally differentiated effector CD8+ T cells (P = 0.001, RHO = -0.84)
3. Number of lymphocytes and:
    a. Naïve CD4+ T cells (P = 0.03, RHO = -0.71)
    b. Early activated CD4+ memory T cells (P = 0.02, RHO = 0.74)
    c. Activated terminally differentiated effector CD8+ T cells (P = 0, RHO = -0.93)
4. Lactate concentration in CSF in nmol/l and:
    a. Resting CD4+ memory T-cell 1 (P = 0.02, RHO = -0.72)
    b. Early activated CD8+ effector memory cells (P = 0.01, RHO = 0.75),
    c. Resting CD4+ memory T-cell 3 (P = 0.04, RHO = -0.66)
    d. Plasma cells (P = 0.01, RHO = 0.78).

Taken together, this strongly supported our cell typing approach, as the clinically measured lymphocyte count correlated well with the proportion of CD4+ and CD8+ typed cell clusters per patient. Interestingly, even age was correlated with cell type proportions of CD8+ T cells.

Furthermore, we acquired categorical clinical data in the form of the following variables:

- #stFLAIRles; number of supratentorial FLAIR hyperintense lesions in magnetic resonance imaging.
- MRI DIS+; criteria for dissemination in space fulfilled by MRI at the time of lumbar puncture. Gd+; any gadolinium-enhancing lesion detected on MR.
- OCB DIT+; criteria for dissemination in time fulfilled by the presence of oligoclonal bands in CSF at the time of lumbal puncture.
- Clinical DIT+; criteria for dissemination in time fulfilled by the presence of at least two clinical relapses.
- EDSS DC; Kurtzke Expanded Disability Status Scale (EDSS) score at discharge, i.e., after treatment of the relapse occurring at the lumbar puncture.
- EDSS FU; latest available clinical follow-up in months after lumbar puncture.
- OCB+; any oligoclonal band (OCB) detected in CSF.
- OCB type and OCB were classified as being either undetectable, restricted to CSF, detected in serum and additionally in CSF, or not determined.
- CSF index; CSF/serum indices for albumin and immunoglobulin G (IgG) were calculated. The CSF index was evaluated as being either unaffected (none), showing intrathecal IgG synthesis (Ig only), showing barrier dysfunction (barrier only), or showing both intrathecal IgG synthesis and barrier dysfunction (barrier and Ig).
- Sex (male or female).
- Prior relapse; number of clinical relapses prior to the one occurring at lumbar puncture.

For the purpose of finding associations between categorical clinical variables and cell type proportions, we implemented a logistic regression (*glm()* R function, package *stats* v.4.0.4). Cell type proportions were used as predictors and individual clinical data as the predicted variable (for example, to test if the cell type proportions are associated with sex, we created a model where cell type proportions were independent variables and sex is the predicted variable). We found no significant association between any of the independent variables and categorical clinical variables (**File 2**).

*3D-network figure for drug candidate interactions with plasma cell DEGs of AIA mice*

To understand the interactions between DEGs and drug candidates, selected based on $z_c < -1.64$ and $d_c < 1$, we created a 3D visualisation (https://scpred.shinyapps.io/3D_network/) for the most central cell type, activated B cells, in the AIA mouse data. Interactions between DEGs (blue) represent protein–protein interactions (PPIs) described in the literature-curated PPI network by do Valle et al. (16). DEG node size is based on fold change. Drug candidates are connected to their respective gene drug targets by edges. Potential drug candidates are shown in red. Established drugs for human rheumatoid arthritis are represented in yellow. The higher the absolute value of a drug on the Y-axis, the higher the drug rank. Drug candidates that counteracted at least one DEG fold change received positive Y-axis values, while drug candidates that did not counteract the fold change of any targeted DEG received negative Y-axis values. By clicking on one of the nodes, the neighbouring nodes are highlighted. Details on the computational environment are provided in https://github.com/SDTC-CPMed/scDrugPrio.

### Supplementary Tables

**Table S1. Drug concentrations for *in vitro* validation studies.**

| **Drug** | **In vitro concentrations (low, medium, high)** |
|---|---|
| **Auranofin**(17, 18) | 10 nM, 50 nM and 250 nM |
| **Dimethyl fumarate(19)** | 5 μM, 25 μM and 100 μM |
| **Irbesartan(20)** | 2 μM, 10 μM and 50 μM |
| **Amrinone(21)** | 10 μM, 50 μM and 250 μM |
| **Isosorbide(22, 23)** | 5 μM, 25 μM and 100 μM |
| **Adapalene(24, 25)** | 200 nM,1 μM and 5 μM |

| **Table S2. Library preparation and sample pooling.** | | | |
|---|---|---|---|
| **Sample** | n libraries prepared | n samples sequenced and merged | n samples per Illumina array |
| Joint_Healthy_mouse_1 | 4 | 4 | 3 |
| Joint_Healthy_mouse_2 | 4 | 4 | 3 |
| Joint_Healthy_mouse_3 | 4 | 5 | 3 (1 for one of the libraries) |
| Joint_Healthy_mouse_4 | 4 | 4 | 3 |
| Joint_Sick_mouse_1 | 4 | 4 | 3 |
| Joint_Sick_mouse_3 | 4 | 4 | 3 |
| Joint_Sick_mouse_4 | 4 | 4 | 3 |
| Joint_Sick_mouse_5 | 4 | 4 | 3 |
| Joint_Sick_mouse_6 | 4 | 4 | 3 |

To increase the read depth for each sample, multiple libraries were prepared and sequenced. For healthy mouse 3, one of the libraries was sequenced twice, wherein one was sequenced alone on one array, without pooling with other samples.

### Supplementary Figures

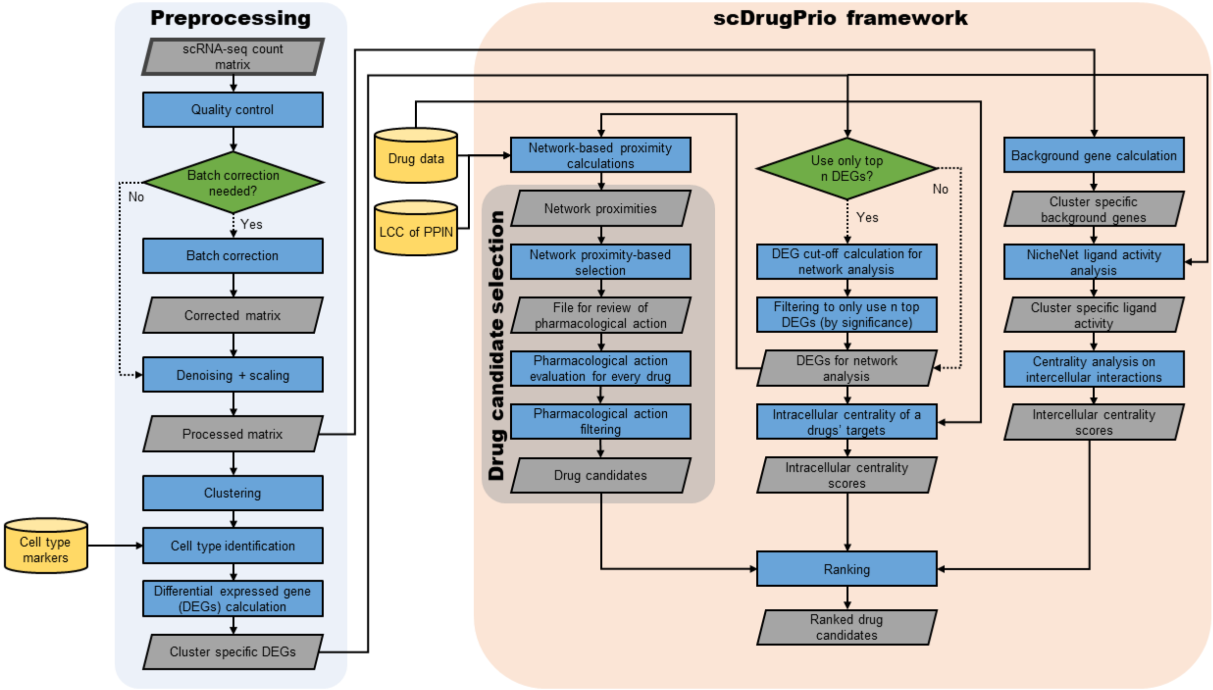


**Fig. S1. scDrugPrio computational workflow, input and output data.** Initially, data are preprocessed following a standard approach, resulting in the calculation of cell type-specific differentially expressed genes (DEGs) between sick and healthy cells. The need for batch correction was evaluated through heterogeneity analyses as described in the methods and supplementary methods section. The DEG calculation utilized either paired samples (e.g., one healthy and one sick sample) from an individual or pooled samples from several healthy controls and patients. The scDrugPrio framework then performs drug selection using DEGs and drug data for network proximity calculation. The cell type-specific drug candidates are aggregated into a final drug ranking using intracellular and intercellular centrality. Gray parallelograms represent data files, yellow cylinders represent external data and green rombs indicate decision points.

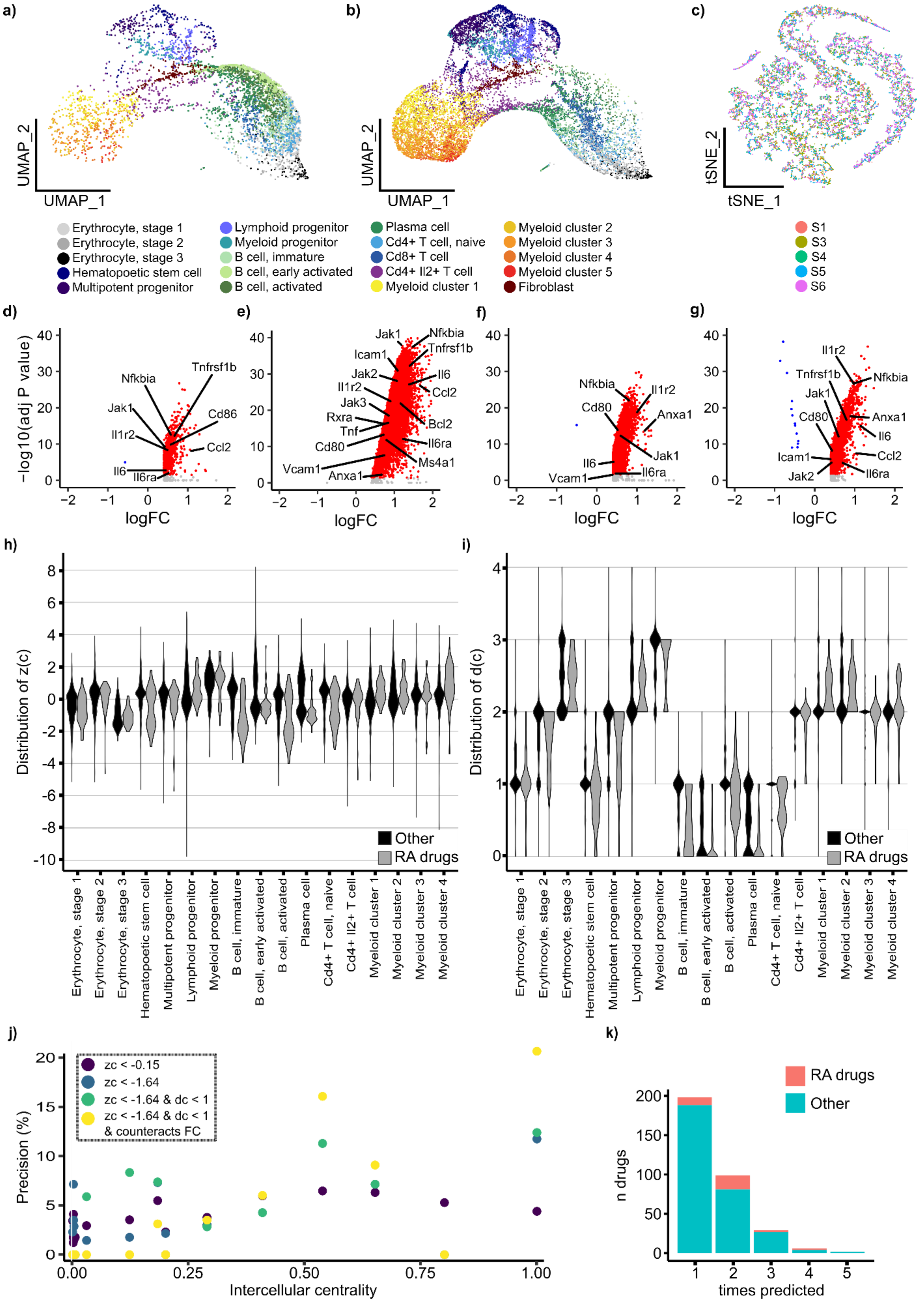
a)
b)
c)
UMAP_2
UMAP_1
tSNE_2
tSNE_1
Erythrocyte, stage 1
Erythrocyte, stage 2
Erythrocyte, stage 3
Hematopoetic stem cell
Multipotent progenitor
Lymphoid progenitor
Myeloid progenitor
B cell, immature
B cell, early activated
B cell, activated
Plasma cell
Cd4+ T cell, naive
Cd8+ T cell
Cd4+ Il2+ T cell
Myeloid cluster 1
Myeloid cluster 2
Myeloid cluster 3
Myeloid cluster 4
Myeloid cluster 5
Fibroblast
S1
S3
S4
S5
S6
d)
e)
f)
g)
−log10(adj P value)
logFC
Nfkbia
Tnfrsf1b
Jak1
Cd86
Il1r2
Ccl2
Il6
Il6ra
Icam1
Jak2
Jak3
Rxra
Tnf
Cd80
Vcam1
Anxa1
Bcl2
Ms4a1
h)
i)
Distribution of z(c)
Distribution of d(c)
Other
RA drugs
j)
k)
zc < -0.15
zc < -1.64
zc < -1.64 & dc < 1
zc < -1.64 & dc < 1
& counteracts FC
Precision (%)
Intercellular centrality
n drugs
times predicted

**Fig. S2**. **Additional cluster information as well as network proximities between rheumatoid arthritis (RA) drugs and DEGs of AIA mice.** UMAP plots including only cells from **a**) control mice or **b)** AIA mice. **c)** tSNE representation of cells' latent features of AIA mice to explore interindividual heterogeneity. Cells were colored based on the AIA mouse from which they originated (S1-S6; as indicated by legend). As cells from different AIA mice overlap nicely, no heterogeneity is apparent. **d–g)** Volcano plots of DEGs (FC > 1.5 and FDR-adjusted $P < 0.05$) that are upregulated (red) or downregulated (blue) in B-cell clusters. DEGs that were targeted by approved RA drugs are indicated by arrows. Plots present the DEGs for **d)** activated B cells, **e)** early activated B cells, **f)** plasma cells, and **g)** immature B cells. **h&i)** Violin plots showing $z_c$ and $d_c$ distributions, respectively, for each cell type. **j)** Precision among cell type drug candidates correlated with the cell type eigenvector centrality in the MCDM. Drug candidates were selected using four different sets of criteria, namely, 1) the basic $z_c$ cut-off of $z_c < -0.15$ suggested by Guney et al.(7), 2) $z_c < -1.64$ (corresponding to P < 0.05), 3) drugs that fulfil both $z_c < -1.64$ and $d_c < 1$ and 4) drugs fulfilling our final selection criteria ($z_c < -1.64$, $d_c < 1$ and counteracting the fold change (FC) of at least one targeted DEG). Pearson correlation was calculated based on drugs passing the final selection criteria (Pearson's $r$ [95% CI] = 0.77 [0.46 – 0.91], $P < 10^{-3}$). **g)** The proportion of known RA drugs was increased among final candidates that reoccurred in several cell types.

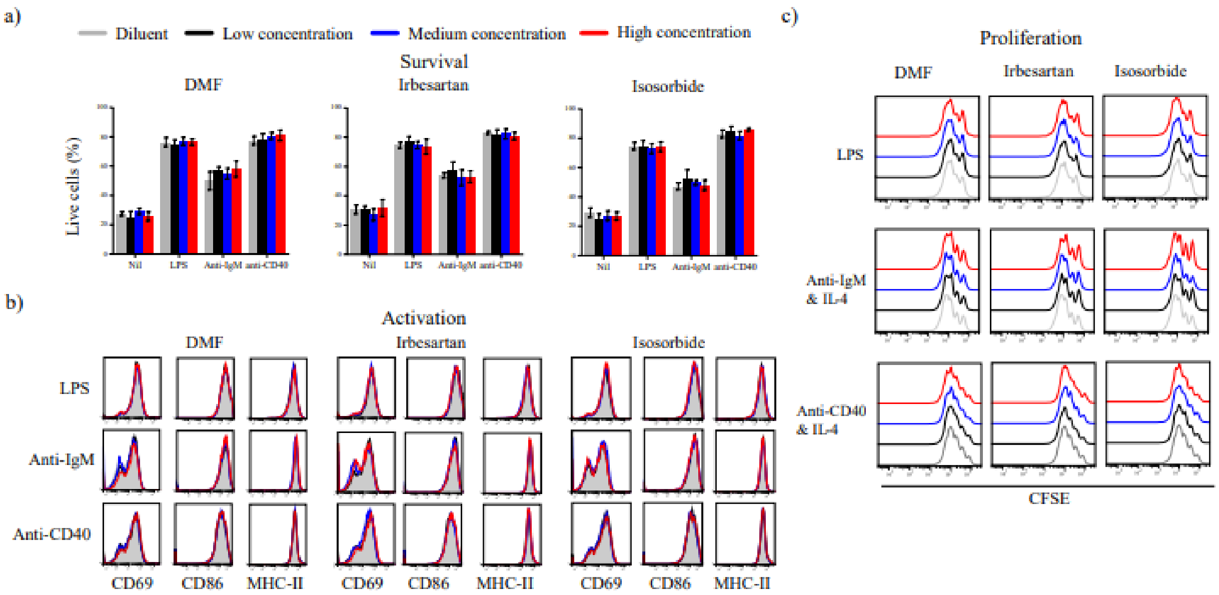


**Fig. S3**. **DMF, irbesartan and isosorbide have no effect on murine B-cell viability, activation or proliferation.** Drug effect of the selected drugs on *in vitro* murine (**a**) B-cell survival, (**b**) activation, and (**c**) proliferation. Purified murine B cells were stimulated with the indicated B-cell modulators in the presence of dimethyl fumarate (DMF), irbesartan, and isosorbide at different concentrations. Drug concentrations can be found in Table S1.

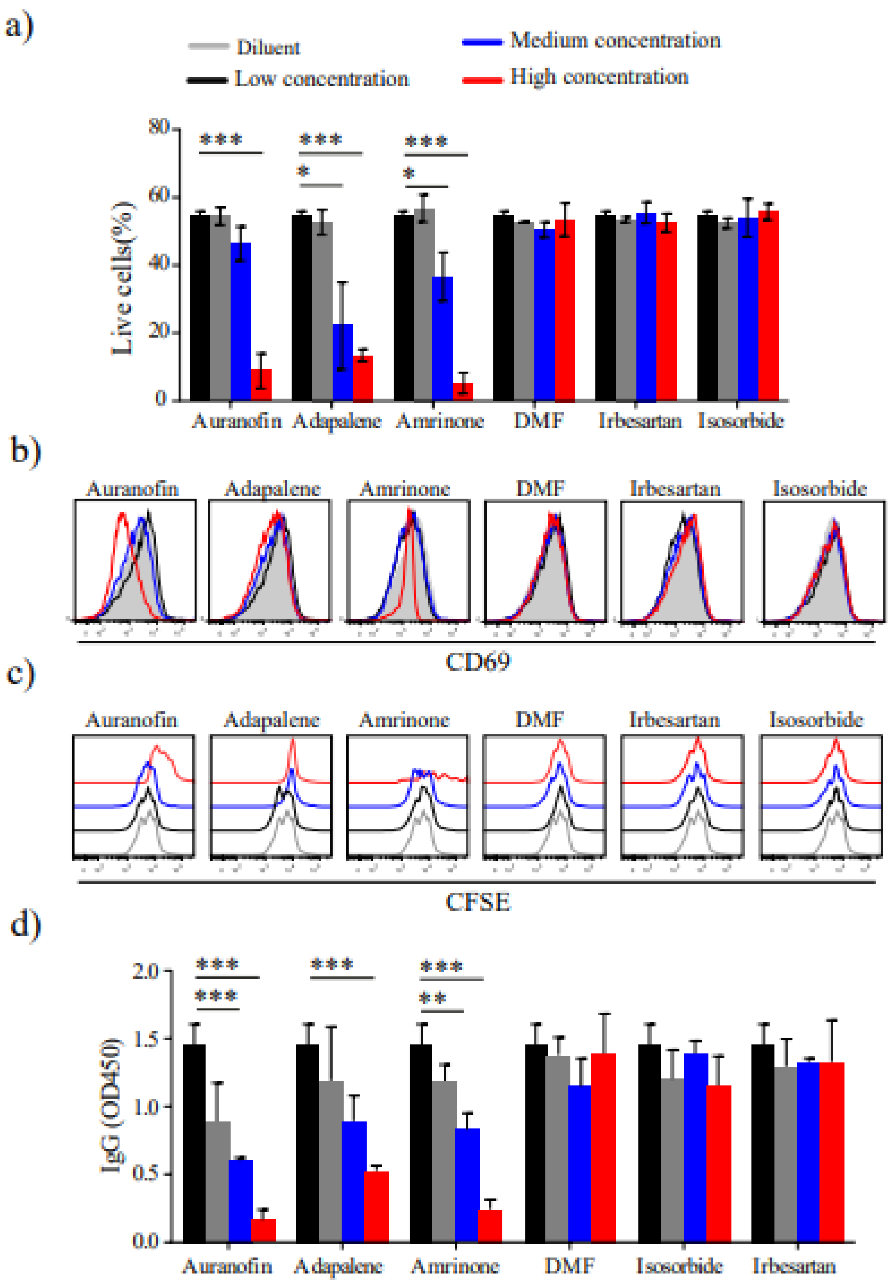


**Fig. S4. *In vitro* analysis of selected drugs on human B-cell (a) viability, (b) activation, (c) proliferation and (d) IgG production.** Purified human naïve B cells were activated with goat anti-human IgG + IgM (5 μg/mL), anti-human CD40 (5 μg/mL) and IL-21 (10 ng/mL) in the presence of selected drugs at the indicated concentrations. Drug concentrations can be found in Table S1. DMF, dimethyl furamate. * P < 0.05, **P < 0.01, ***P < 0.001.

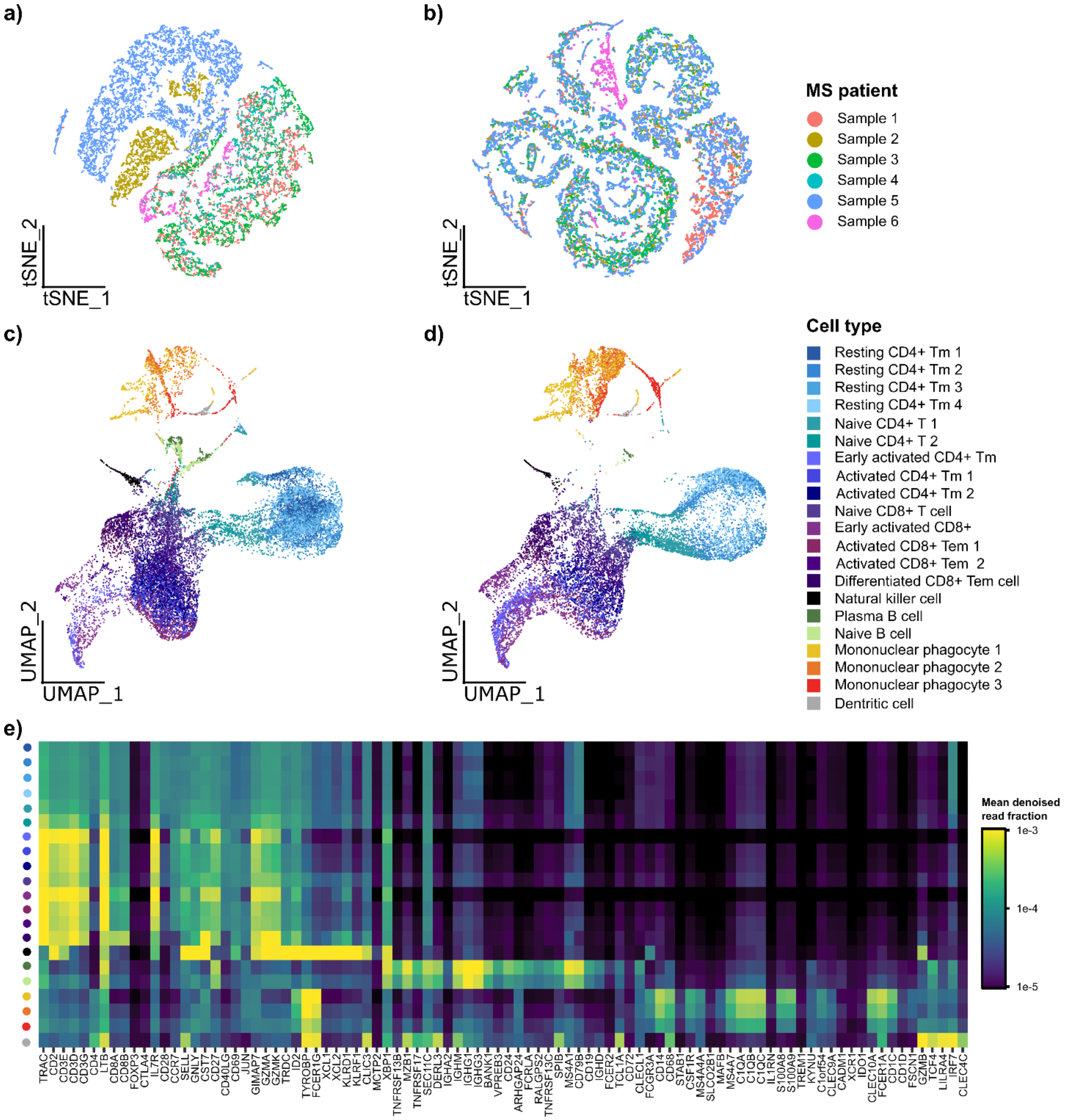


**Fig. S5. Cell typing and selection of the number of DEGs for network-based drug screening in scRNA-seq data from human multiple sclerosis (MS) patients.** tSNE representation of latent features for MS patients **a)** before and **b)** after batch correction. Cells were coloured based on which MS patient they originated from (patient 1 to 6). In contrast to **Fig. S1c,** we can see that cells from different patients do not mix well before batch correction (**a**), indicating transcriptomic heterogeneity between patients. UMAP clustering plots corresponding to **Fig. 5a** including only cerebrospinal fluid cells from **c**) MS patients or **d)** controls (idiopathic intracranial hypertension patients). **e)** Cell typing by cluster means of DCA-denoised, log10 adjusted gene expression fractions for known marker genes. Abbreviations: Tm, memory T cell; Tem, effector memory T cell.

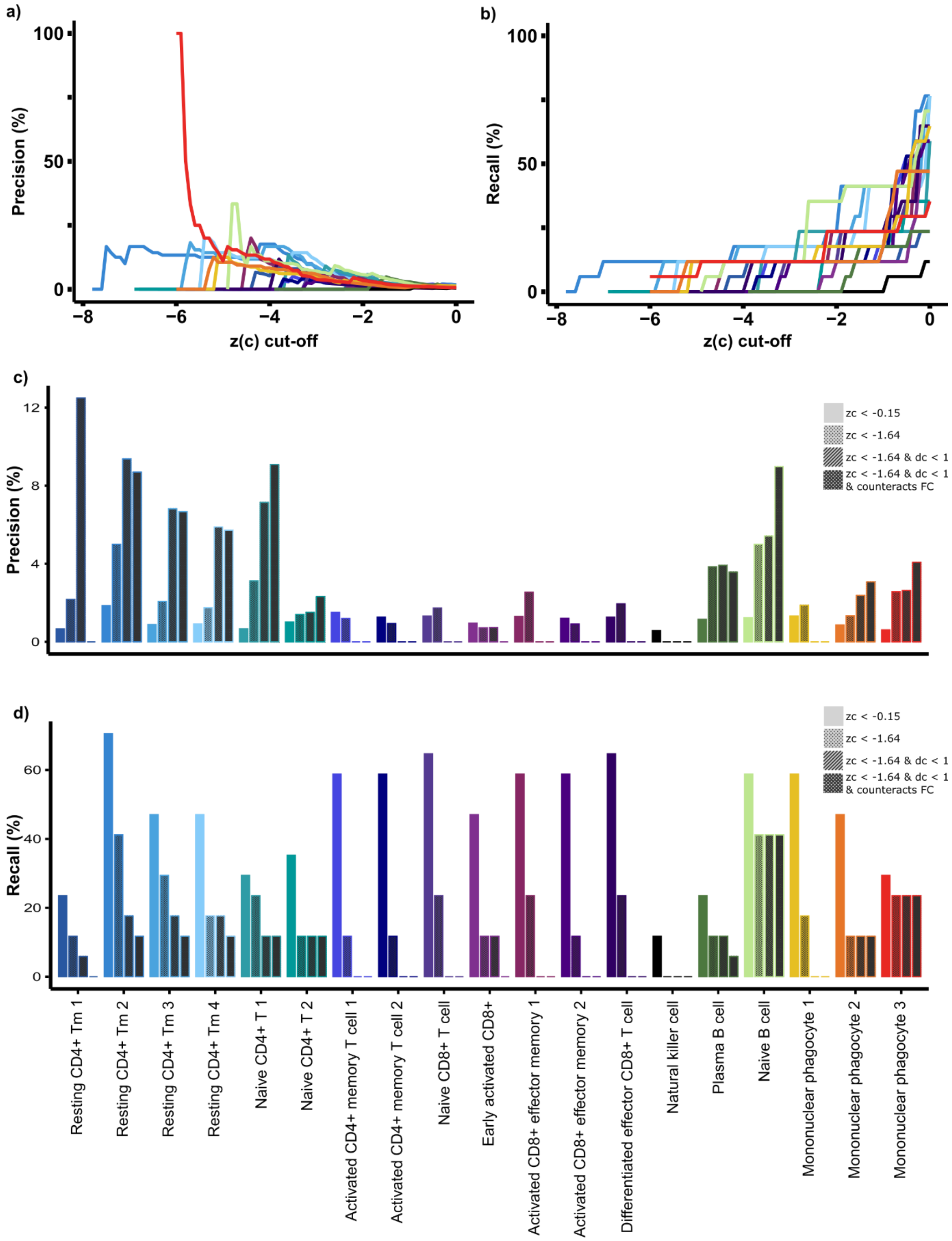


**Fig. S6. Drug candidate selection for multiple sclerosis (MS) patients. a)** Precision and **b)** recall curves for approved drugs among candidates at different $z_c$ cut-offs indicate that decreasing $z_c$ increases precision among central cell types in the MCDM. **c&d)** Precision (c) and recall (d) after stepwise application of drug selection criteria to candidates selected based on network distances to the top 3,000 significant DEGs.

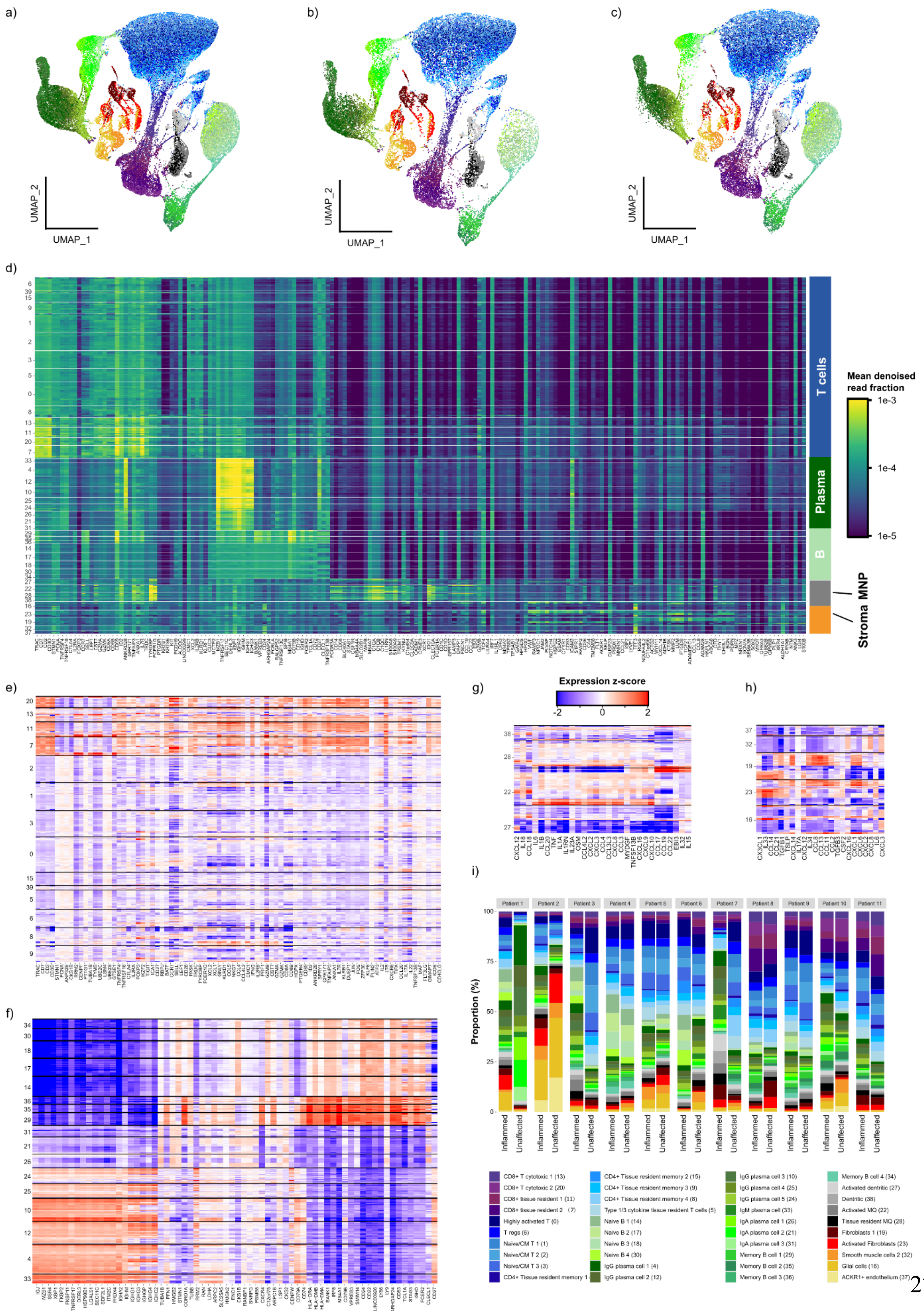
a)
b)
c)
UMAP_2
UMAP_1
d)
T cells
Plasma
B
Stroma
MNP
Mean denoised read fraction
1e-3
1e-4
1e-5
e)
f)
g)
h)
Expression z-score
-2
0
2
i)
Proportion (%)
Patient 1
Patient 2
Patient 3
Patient 4
Patient 5
Patient 6
Patient 7
Patient 8
Patient 9
Patient 10
Patient 11
Inflammed
Unaffected
CD8+ T cytotoxic 1 (13)
CD8+ T cytotoxic 2 (20)
CD8+ tissue resident 1 (11)
CD8+ tissue resident 2 (7)
Highly activated T (0)
T regs (6)
Naive/CM T 1 (1)
Naive/CM T 2 (2)
Naive/CM T 3 (3)
CD4+ Tissue resident memory 1
CD4+ Tissue resident memory 2 (15)
CD4+ Tissue resident memory 3 (9)
CD4+ Tissue resident memory 4 (8)
Type 1/3 cytokine tissue resident T cells (5)
Naive B 1 (14)
Naive B 2 (17)
Naive B 3 (18)
Naive B 4 (30)
IgG plasma cell 1 (4)
IgG plasma cell 2 (12)
IgG plasma cell 3 (10)
IgG plasma cell 4 (25)
IgG plasma cell 5 (24)
IgM plasma cell (33)
IgA plasma cell 1 (26)
IgA plasma cell 2 (21)
IgA plasma cell 3 (31)
Memory B cell 1 (29)
Memory B cell 2 (35)
Memory B cell 3 (36)
Memory B cell 4 (34)
Activated dentritic (27)
Dentritic (38)
Activated MQ (22)
Tissue resident MQ (28)
Fibroblasts 1 (19)
Activated Fibroblasts (23)
Smooth muscle cells 2 (32)
Glial cells (16)
ACKR1+ endothelium (37)

**Fig. S7. Clustering and cell typing of Crohn's disease patient data. a)** tSNE plot including all cells from inflamed and unaffected intestinal lesions of all patients. Clusters are colour coded by cell types in the colour legend of i). **b&c)** are tSNE plots that include only cells from unaffected and inflamed intestinal biopsies. **d)** Heatmap representing individual cell expression of major cell type marker genes. Gene expression values correspond to DCA(26)-adjusted gene expression value fractions of a cell's total DCA-adjusted gene expression values. Individual cells are grouped by the cluster that is represented by a number on the y-axis corresponding to the number in the colour legend of i). White lines separate clusters for visibility. Bars on the right side indicate the major cell type. **e-g)** Heatmaps of single-cell gene expression fraction-based z-score for further stratification of T cells (e), B cells (f), MNPs (g) and stroma/glia cells (h). **i)** Stacked bar plots representing the fractions of cells per cell type and sample. Abbreviations: ILC, innate lymphocyte cells; MNP, mononuclear phagocytes. Heatmap x labels can be found in **File 3.**

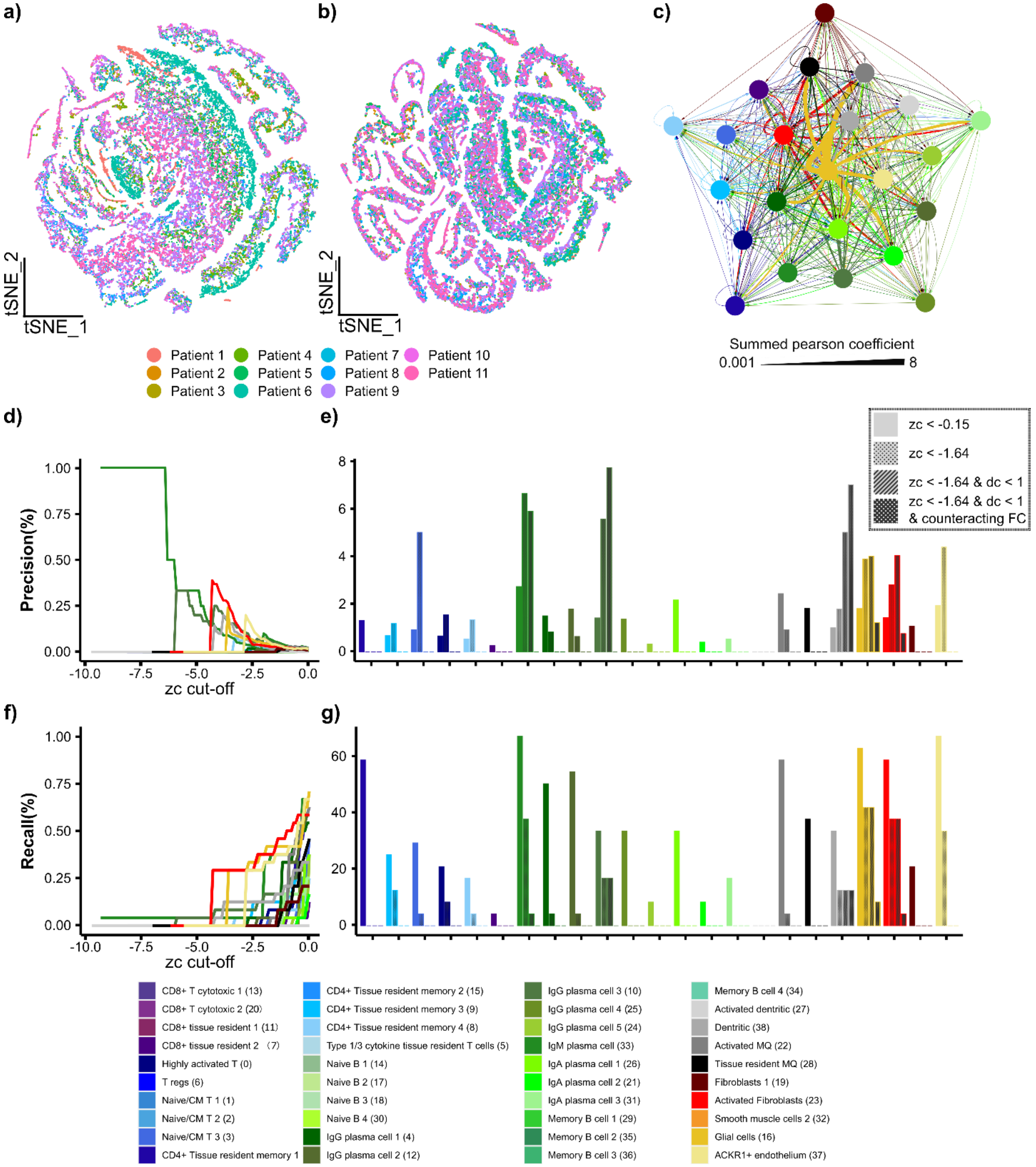


**Fig. S8. Clustering and cell typing of Crohn's disease patient data.** tSNE representation of cell latent features for CD patients **a)** before and **b)** after batch correction. Cells were coloured based on which CD patient they originated from (patient 1 to 11, see **File 3** for more information). Before batch correction, (**a**) cells from patients do not mix well, indicating underlying differences in gene expression profiles. **c)** MCDM based on NicheNet (27)-derived cellular interactions among cell types. Nodes represent cell types, and directed edges indicate ligand interactions from NicheNet. Edge width corresponds to the summed Pearson correlation coefficient of all ligands in the upstream cell type having a potential effect on the downstream cell type. **d&f)** Precision & recall curves at different $z_c$ cut-offs when entering only the top 1,800 DEGs into network distance calculations. **e&g)** Precision and recall for known CD drugs depending on selection criteria (indicated by pattern) using only the top 1,800 significant DEGs for network calculations. Colours in **c-g)** match the colour legend in the bottom.

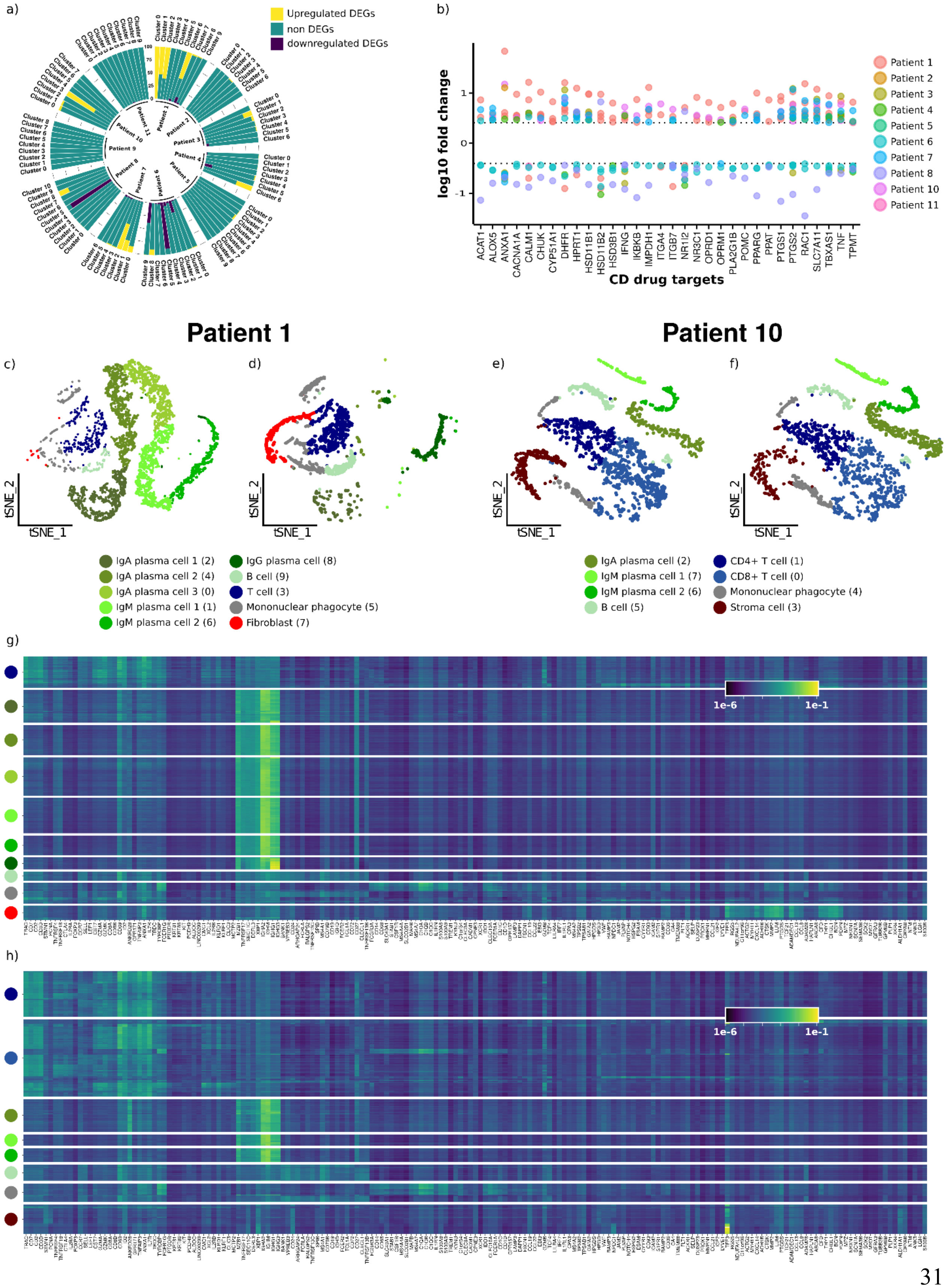
a)
Upregulated DEGs
non DEGs
downregulated DEGs
b)
log10 fold change
CD drug targets
ACAT1
ALOX5
ANXA1
CACNA1A
CALM1
CHUK
CYP51A1
DHFR
HPRT1
HSD11B1
HSD11B2
HSD3B1
IFNG
IKBKB
IMPDH1
ITGA4
NR1I2
NR3C1
OPRD1
OPRM1
PLA2G1B
POMC
PPARG
PPAT
PTGS1
PTGS2
RAC1
SLC7A11
TBXAS1
TNF
TPMT
Patient 1
Patient 2
Patient 3
Patient 4
Patient 5
Patient 6
Patient 7
Patient 8
Patient 10
Patient 11
Patient 1
Patient 10
c)
d)
e)
f)
tSNE_1
tSNE_2
IgA plasma cell 1 (2)
IgA plasma cell 2 (4)
IgA plasma cell 3 (0)
IgM plasma cell 1 (1)
IgM plasma cell 2 (6)
IgG plasma cell (8)
B cell (9)
T cell (3)
Mononuclear phagocyte (5)
Fibroblast (7)
IgA plasma cell (2)
IgM plasma cell 1 (7)
IgM plasma cell 2 (6)
B cell (5)
CD4+ T cell (1)
CD8+ T cell (0)
Mononuclear phagocyte (4)
Stroma cell (3)
g)
1e-6
1e-1
h)
1e-6
1e-1

**Fig. S9. Clustering and cell typing of individual Crohn's disease (CD) patients. a)** Fraction of differentially expressed genes out of all genes. For this, data were divided by patient and clustered separately. For patient 1 and 10, clusters were cell typed, and cluster ID in a) corresponds to the number in parentheses in the figure legend of c-d) and e-f), respectively. **b)** Fold change of known CD drug targets for all patient-specific DEGs in all clusters. The colour represents the patients; in some instances, several clusters of a patient differentially expressed a certain drug target. **c-d)** tSNE plot showing only cells from the uninflamed (c) and inflamed (d) biopsy of patient 1. **e-f)** Same as c-d) for patient 10. **g)** Heatmap representing individual cell expression of major cell type marker genes used for cell typing the clusters of patient 1. Gene expression values correspond to DCA(26) adjusted gene expression value fractions of a cell's total DCA adjusted gene expression values. Individual cells are grouped by cluster. Colours on the y-axis correspond to the colour legend of c-d). Comparable to the heatmap in **Fig. S4d**. **h)** Same as g) for patient 10; colours on the Y-axis correspond to colour legend of e-f).

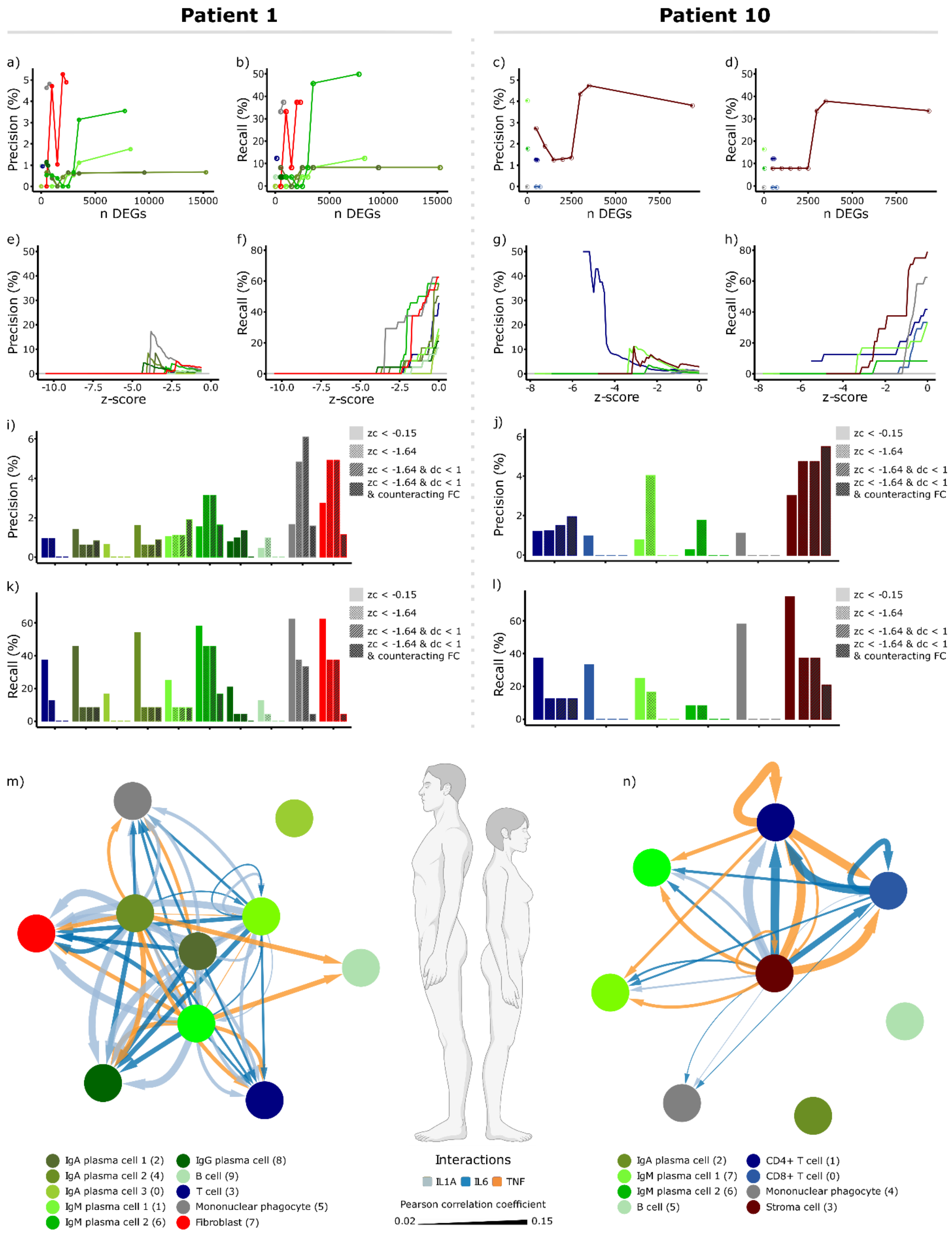

Patient 1
Patient 10
a)
b)
c)
d)
e)
f)
g)
h)
i)
j)
k)
l)
m)
n)
Precision (%)
Recall (%)
n DEGs
z-score
zc < -0.15
zc < -1.64
zc < -1.64 & dc < 1
zc < -1.64 & dc < 1
& counteracting FC
IgA plasma cell 1 (2)
IgA plasma cell 2 (4)
IgA plasma cell 3 (0)
IgM plasma cell 1 (1)
IgM plasma cell 2 (6)
IgG plasma cell (8)
B cell (9)
T cell (3)
Mononuclear phagocyte (5)
Fibroblast (7)
Interactions
IL1A IL6 TNF
Pearson correlation coefficient
0.02 0.15
IgA plasma cell (2)
IgM plasma cell 1 (7)
IgM plasma cell 2 (6)
B cell (5)
CD4+ T cell (1)
CD8+ T cell (0)
Mononuclear phagocyte (4)
Stroma cell (3)


492

**Fig. S10**. **Drug prediction outcomes based on individual Crohn's disease patients**. The panel is divided in the middle and represents outcomes for patients 1 and 10 on the left and right sides, respectively. **a-d)** Cell type precision and recall for known CD drugs at $z_c < -1.64$, shown as a function of the number of an individual patient's top significant DEGs that entered network distance calculations. **e-h)** Precision & recall curves at different $z_c$ cut-offs when entering only the top 3,500 DEGs for each patient into network distance calculations. **i-l)** Precision and recall for known CD drugs depending on selection criteria (indicated by pattern), using only the top 3,500 significant DEGs of each patient for network calculations. **m-n)** MCDMs highlighting only *IL1A*, *IL6* and *TNF* ligand interactions between cell types from patients 1 and 10. Edge width is scaled by the NicheNet(27) Pearson correlation coefficient, which is a measure of each single ligand association with the observed differential gene expression in the downstream cell type.

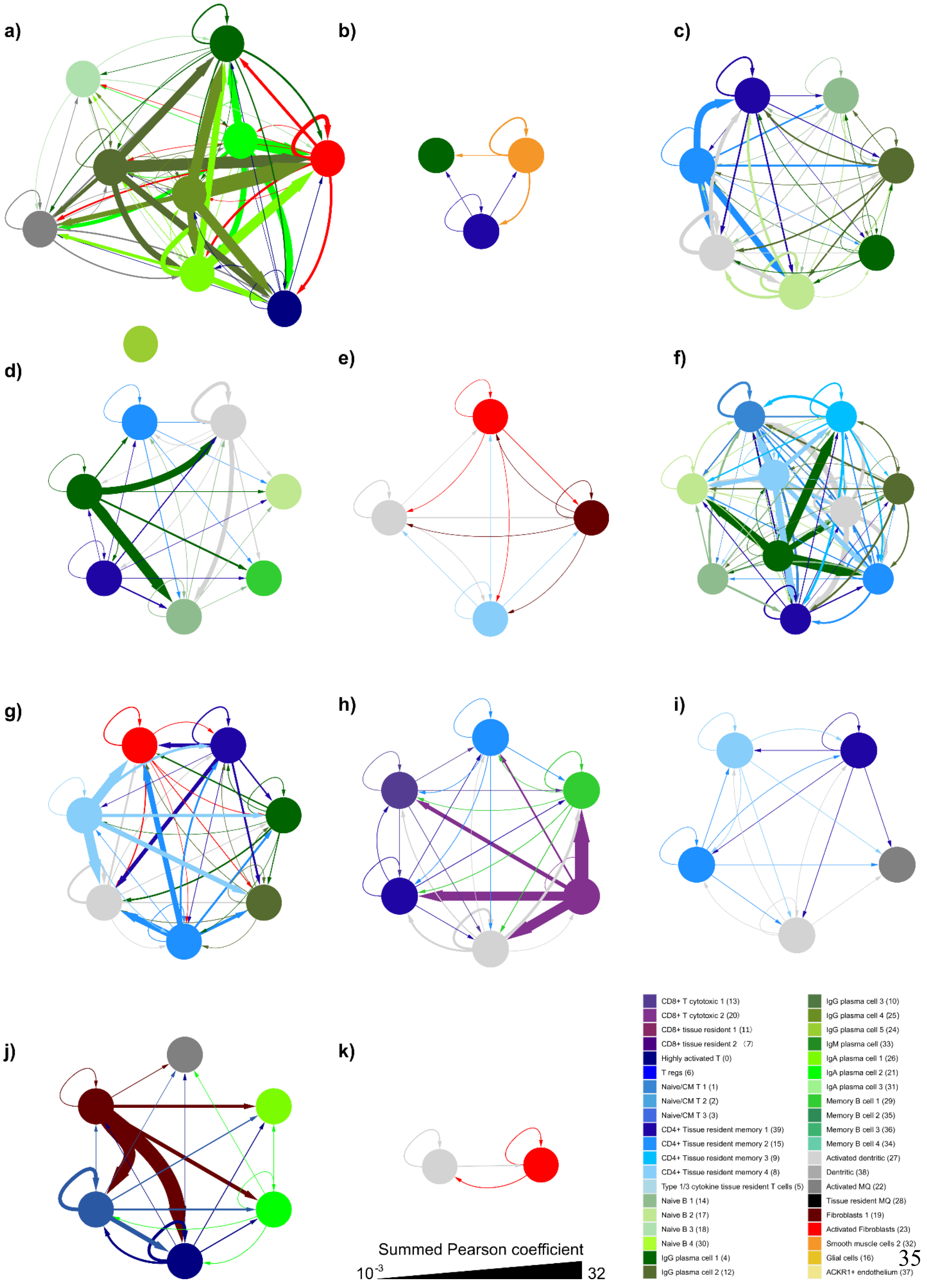
a)
b)
c)
d)
e)
f)
g)
h)
i)
j)
k)
Summed Pearson coefficient
$10^{-3}$
32
CD8+ T cytotoxic 1 (13)
CD8+ T cytotoxic 2 (20)
CD8+ tissue resident 1 (11)
CD8+ tissue resident 2 （7）
Highly activated T (0)
T regs (6)
Naive/CM T 1 (1)
Naive/CM T 2 (2)
Naive/CM T 3 (3)
CD4+ Tissue resident memory 1 (39)
CD4+ Tissue resident memory 2 (15)
CD4+ Tissue resident memory 3 (9)
CD4+ Tissue resident memory 4 (8)
Type 1/3 cytokine tissue resident T cells (5)
Naive B 1 (14)
Naive B 2 (17)
Naive B 3 (18)
Naive B 4 (30)
IgG plasma cell 1 (4)
IgG plasma cell 2 (12)
IgG plasma cell 3 (10)
IgG plasma cell 4 (25)
IgG plasma cell 5 (24)
IgM plasma cell (33)
IgA plasma cell 1 (26)
IgA plasma cell 2 (21)
IgA plasma cell 3 (31)
Memory B cell 1 (29)
Memory B cell 2 (35)
Memory B cell 3 (36)
Memory B cell 4 (34)
Activated dentritic (27)
Dentritic (38)
Activated MQ (22)
Tissue resident MQ (28)
Fibroblasts 1 (19)
Activated Fibroblasts (23)
Smooth muscle cells 2 (32)
Glial cells (16)
ACKR1+ endothelium (37)

**Fig. S11. MCDMs of all individual CD patients. a-k)** Correspond to patients 1 to 11. Nodes correspond to cell types that expressed DEGs and are coloured according to the colour legend in the lower right corner. Patient 1 had one cell type that was not connected to the rest of the remaining MCDM. Except for patients 1 and 10, which were cell typed as described in **Fig. S8g&h, c**ell types were derived from the overall cell typing for the pooled patient data (**Fig. S6d**). The directed edges are coloured by colour of the upstream cell type. Edge width corresponds to the summed Pearson correlation coefficient.

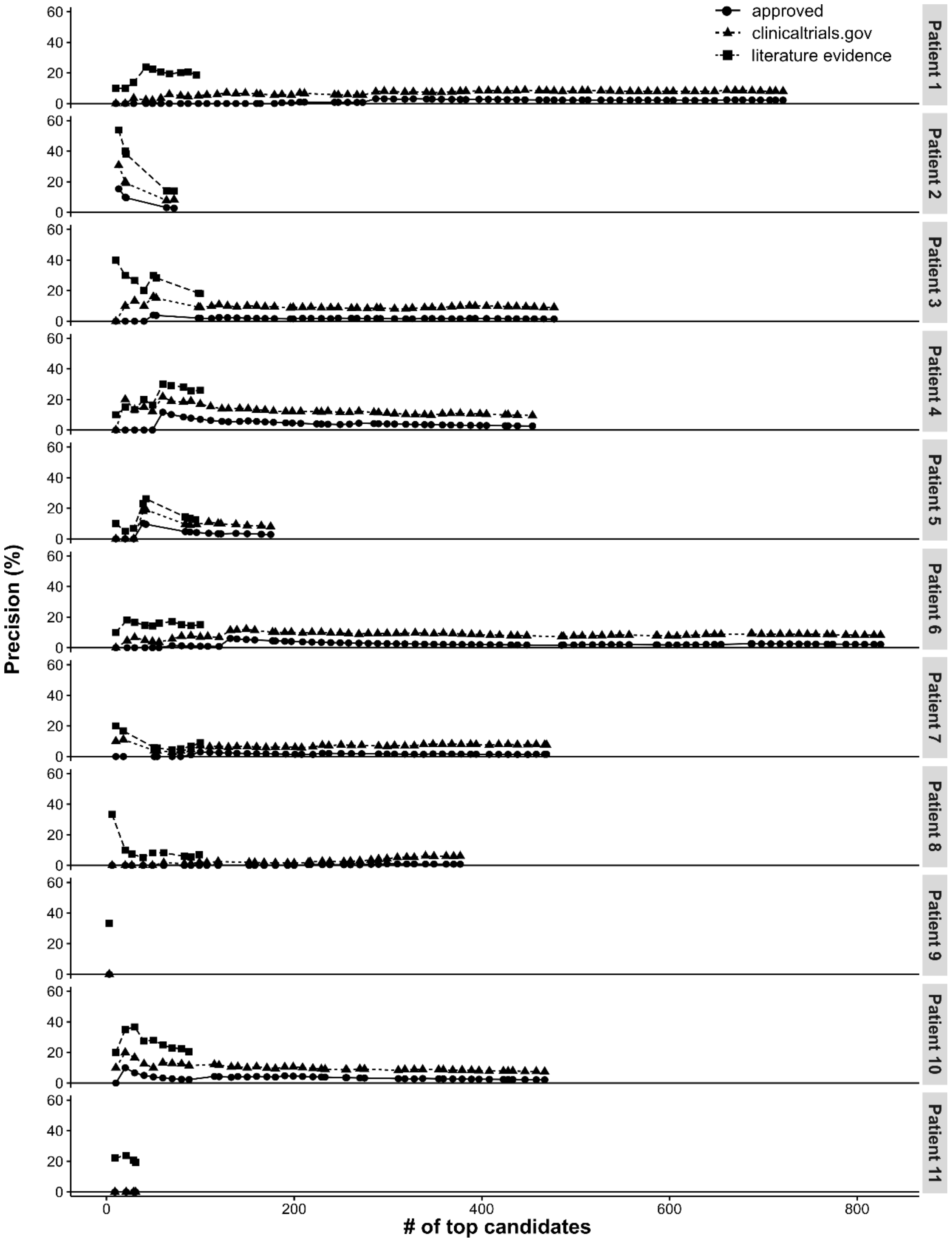


**Fig. S12. Precision among ranked candidates of all individual CD patients.** Drug rank on the x-axis and precision on the y-axis. Showing precision for approved CD drugs, drugs were registered for clinical trials in CD as well as for drugs with literature evidence as indicated by shape and line type. Literature evidence was only collected for candidates with a rank up to 100.

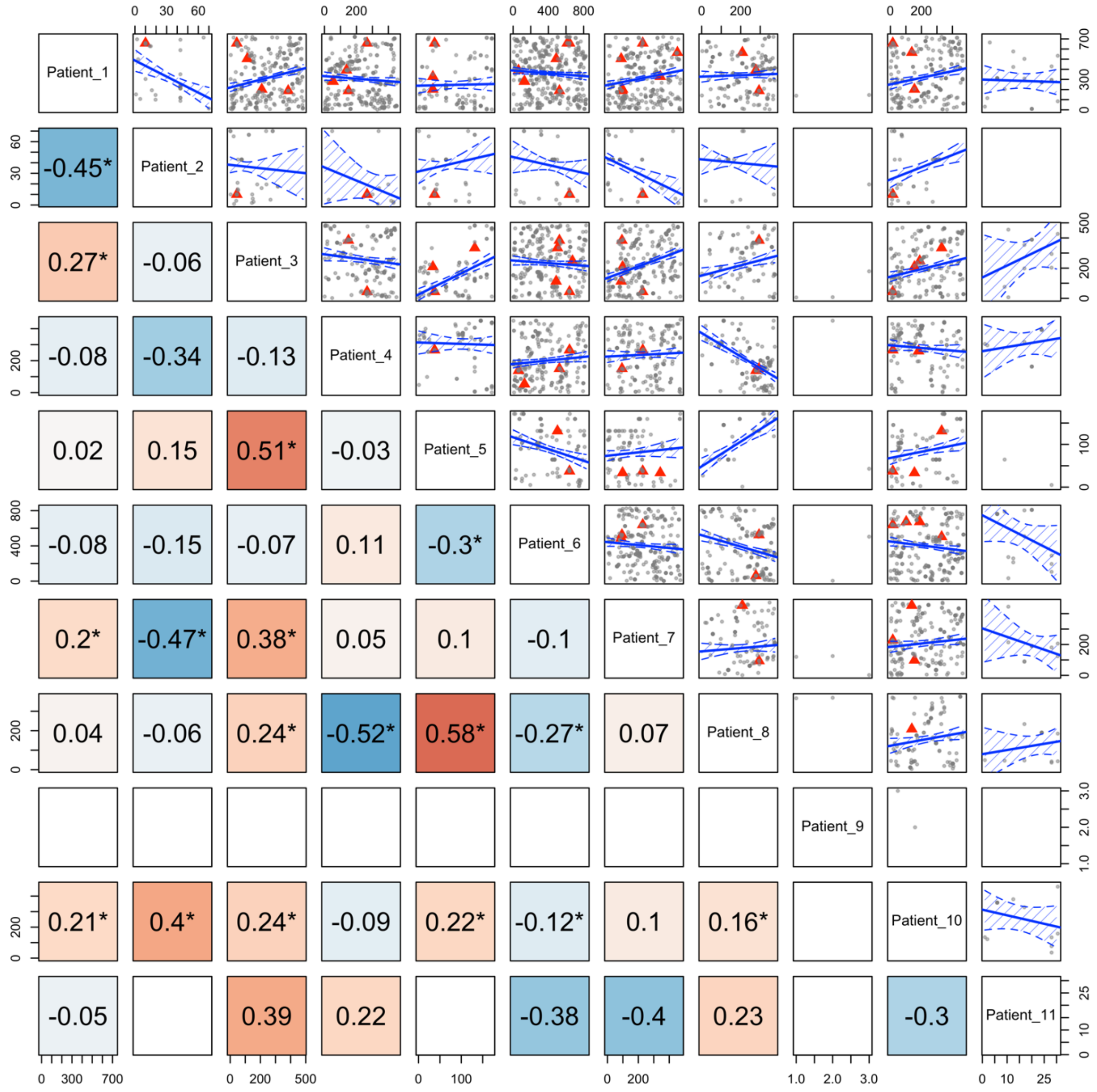


**Fig. S13. Correlation between individual Crohn's disease patient drug ranks.** Patients are indicated on the diagonal axis. The upper right portion of this panel presents scatter plots depicting the correlation of drug ranks in drug predictions from two individual patients. Known CD drugs are depicted by red triangles, and all other drugs are presented as gray dots. In cases where more than 5 drugs overlapped between patients, a Pearson correlation was calcuated. The correlation coefficients are presented in the lower left panel and are coloured by correlation coefficient (0 > blue and red > 0), and significant P values are indicated by asterisks. Blue lines in the upper right panel show the Pearson correlation coefficient and 95% confidence intervals.

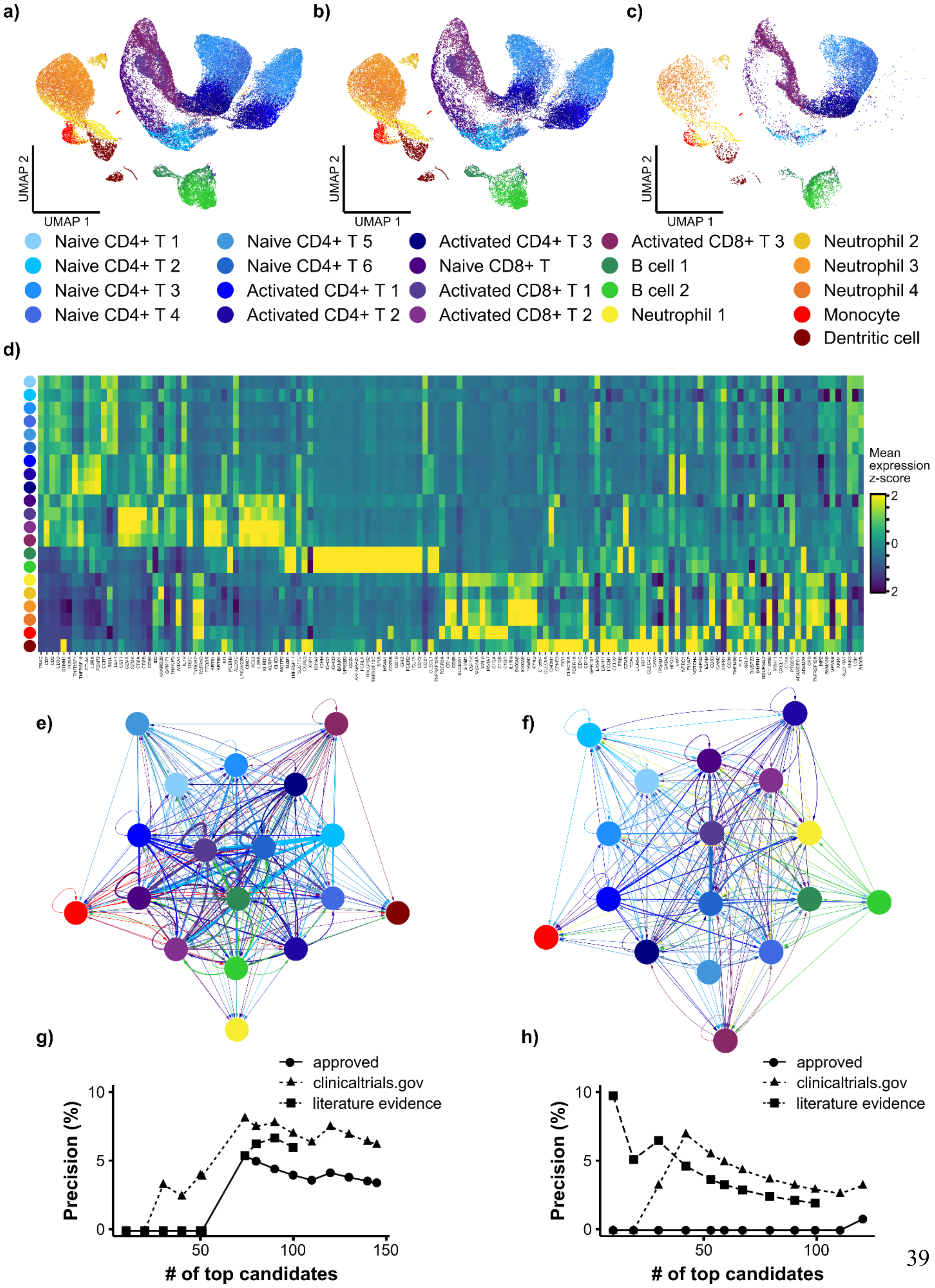
a)
b)
c)
UMAP 2
UMAP 1
Naive CD4+ T 1
Naive CD4+ T 2
Naive CD4+ T 3
Naive CD4+ T 4
Naive CD4+ T 5
Naive CD4+ T 6
Activated CD4+ T 1
Activated CD4+ T 2
Activated CD4+ T 3
Naive CD8+ T
Activated CD8+ T 1
Activated CD8+ T 2
Activated CD8+ T 3
B cell 1
B cell 2
Neutrophil 1
Neutrophil 2
Neutrophil 3
Neutrophil 4
Monocyte
Dentritic cell
d)
Mean expression z-score
2
0
-2
e)
f)
g)
h)
approved
clinicaltrials.gov
literature evidence
Precision (%)
### of top candidates
0
5
10
50
100
150

**Fig. S14. scDrugPrio applied to psoriatic arthritis patients who were or were not anti-IL17 responders.** UMAP visualisation of cells from **a)** controls and PsA patients (both responders and nonresponders to anti-IL17), **b)** only PsA patients, and **c)** only controls. **d)** Cell typing performed on marker genes; cell type is indicated by cluster colours to the left. MCDMs were created for **e)** anti-IL17 responders and **f)** nonresponders. Drug rankings for **g)** responders and **h)** nonresponders were derived separately. Responders achieve a precision of 4% for approved PsA drugs and 6% for drugs with literature evidence among the top 100 ranking drugs. The approved PsA drugs are cortisone derivates. Nonresponders have a precision of 0% for approved PsA drugs and 2% for drugs with literature evidence among the top 100 ranking drugs.

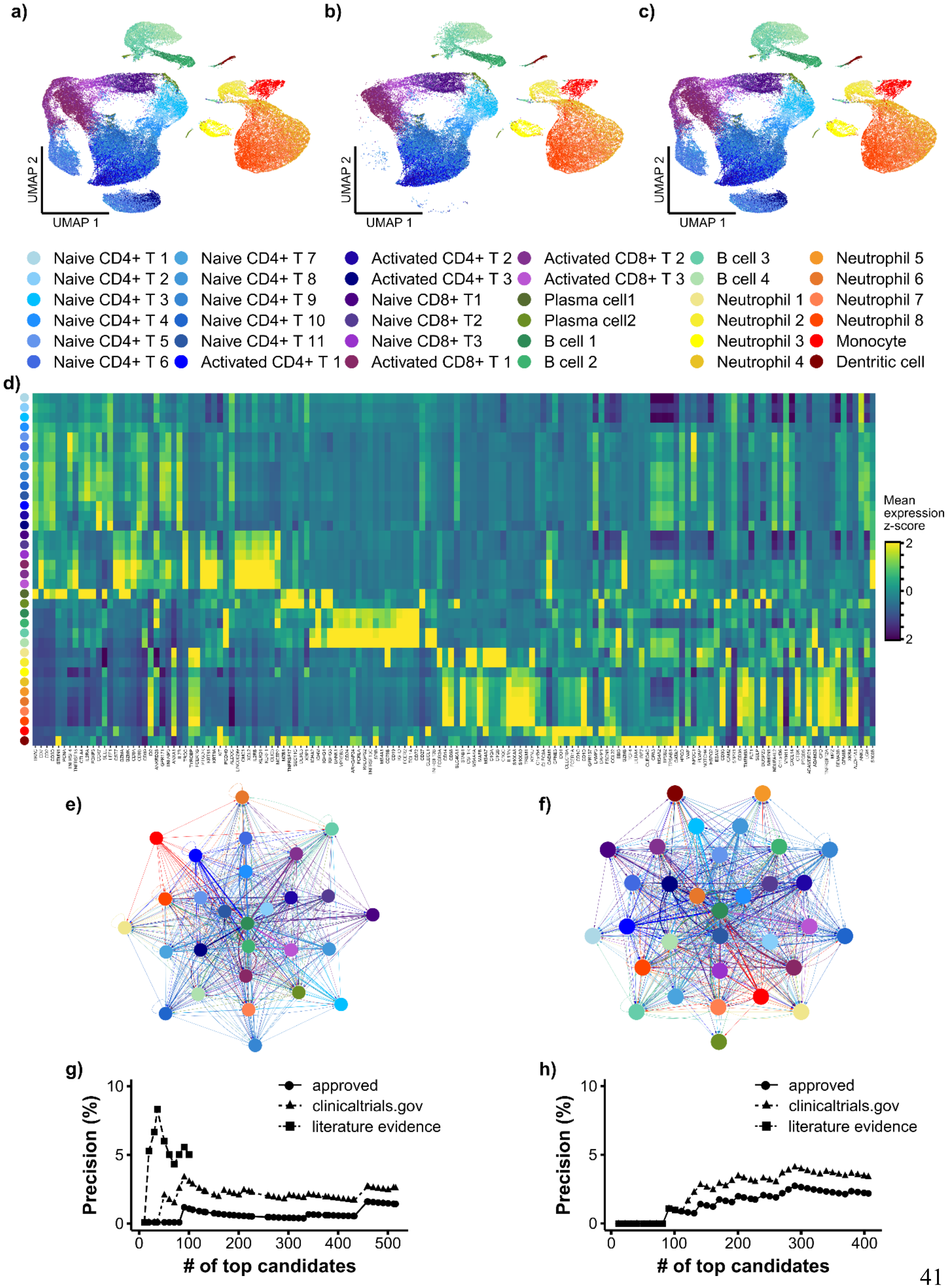
a)
b)
c)
UMAP 2
UMAP 1
Naive CD4+ T 1
Naive CD4+ T 2
Naive CD4+ T 3
Naive CD4+ T 4
Naive CD4+ T 5
Naive CD4+ T 6
Naive CD4+ T 7
Naive CD4+ T 8
Naive CD4+ T 9
Naive CD4+ T 10
Naive CD4+ T 11
Activated CD4+ T 1
Activated CD4+ T 2
Activated CD4+ T 3
Naive CD8+ T1
Naive CD8+ T2
Naive CD8+ T3
Activated CD8+ T 1
Activated CD8+ T 2
Activated CD8+ T 3
Plasma cell1
Plasma cell2
B cell 1
B cell 2
B cell 3
B cell 4
Neutrophil 1
Neutrophil 2
Neutrophil 3
Neutrophil 4
Neutrophil 5
Neutrophil 6
Neutrophil 7
Neutrophil 8
Monocyte
Dentritic cell
d)
Mean expression z-score
2
0
-2
e)
f)
g)
h)
Precision (%)
### of top candidates
approved
clinicaltrials.gov
literature evidence

**Fig. S15. scDrugPrio applied to psoriatic arthritis patients who were or were not anti-TNF responders.** UMAP visualisation of cells from **a)** controls and PsA patients (both responders and nonresponders to anti-TNF), **b)** only PsA patients, and **c)** only controls. **d)** Cell typing performed on marker genes; cell type is indicated by cluster colours to the left. MCDMs were created for **e)** anti-TNF responders and **f)** nonresponders. Drug rankings for **g)** responders and **h)** nonresponders were derived separately. Responders achieve a precision of 1% for approved PsA drugs and 4% for drugs with literature evidence among the top 100 ranking drugs. The approved PsA drugs are cortisone derivates. Nonresponders have a precision of 1% for approved PsA drugs and 1% for drugs with literature evidence among the top 100 ranking drugs.

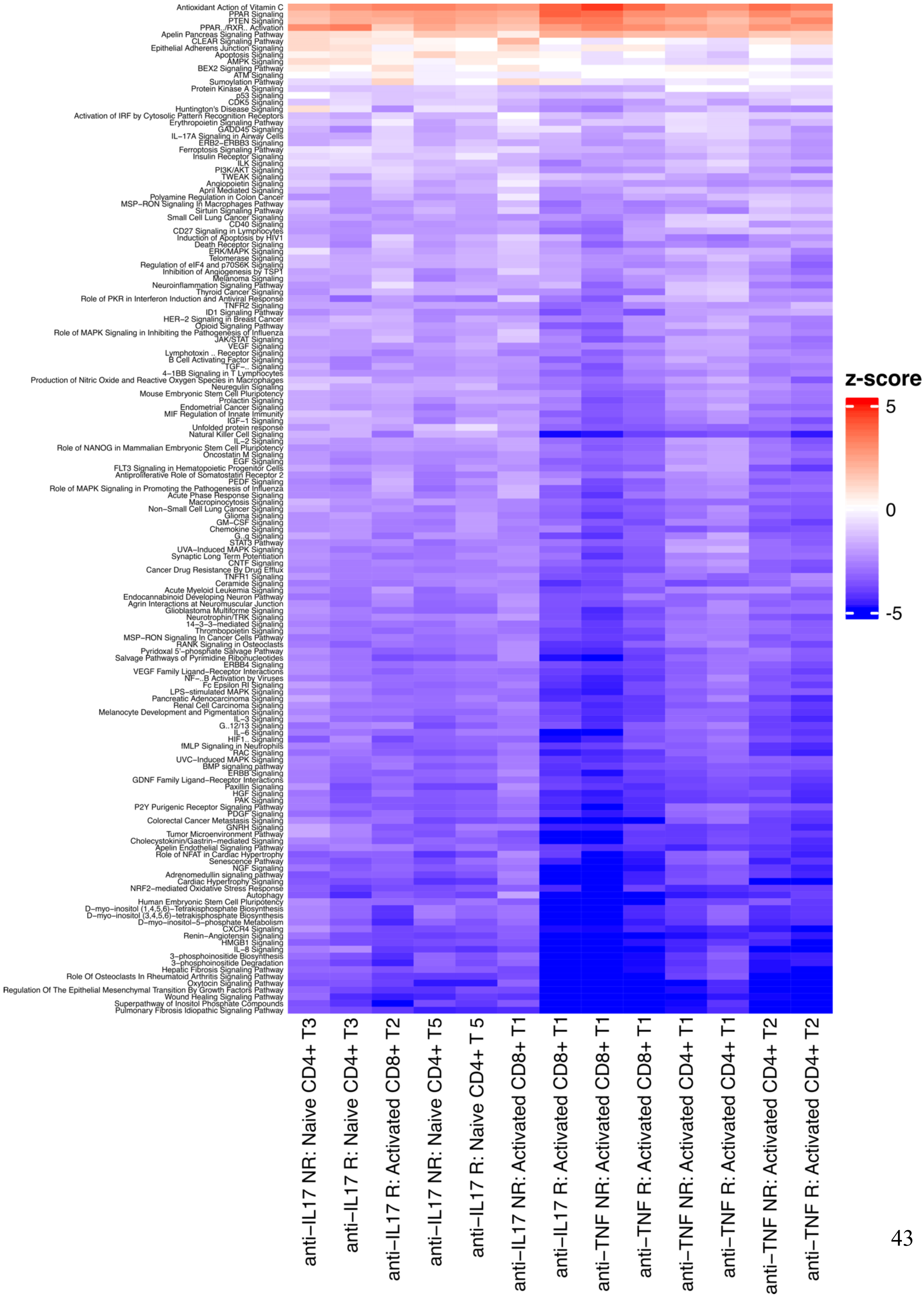

Antioxidant Action of Vitamin C
PPAR Signaling
PTEN Signaling
PPAR../RXR.. Activation
Apelin Pancreas Signaling Pathway
CLEAR Signaling Pathway
Epithelial Adherens Junction Signaling
Apoptosis Signaling
AMPK Signaling
BEX2 Signaling Pathway
ATM Signaling
Sumoylation Pathway
Protein Kinase A Signaling
p53 Signaling
CDK5 Signaling
Huntington's Disease Signaling
Activation of IRF by Cytosolic Pattern Recognition Receptors
Erythropoietin Signaling Pathway
GADD45 Signaling
IL−17A Signaling in Airway Cells
ERB2−ERBB3 Signaling
Ferroptosis Signaling Pathway
Insulin Receptor Signaling
ILK Signaling
PI3K/AKT Signaling
TWEAK Signaling
Angiopoietin Signaling
April Mediated Signaling
Polyamine Regulation in Colon Cancer
MSP−RON Signaling In Macrophages Pathway
Sirtuin Signaling Pathway
Small Cell Lung Cancer Signaling
CD40 Signaling
CD27 Signaling in Lymphocytes
Induction of Apoptosis by HIV1
Death Receptor Signaling
ERK/MAPK Signaling
Telomerase Signaling
Regulation of eIF4 and p70S6K Signaling
Inhibition of Angiogenesis by TSP1
Melanoma Signaling
Neuroinflammation Signaling Pathway
Thyroid Cancer Signaling
Role of PKR in Interferon Induction and Antiviral Response
TNFR2 Signaling
ID1 Signaling Pathway
HER−2 Signaling in Breast Cancer
Opioid Signaling Pathway
Role of MAPK Signaling in Inhibiting the Pathogenesis of Influenza
JAK/STAT Signaling
VEGF Signaling
Lymphotoxin .. Receptor Signaling
B Cell Activating Factor Signaling
TGF−.. Signaling
4−1BB Signaling in T Lymphocytes
Production of Nitric Oxide and Reactive Oxygen Species in Macrophages
Neuregulin Signaling
Mouse Embryonic Stem Cell Pluripotency
Prolactin Signaling
Endometrial Cancer Signaling
MIF Regulation of Innate Immunity
IGF−1 Signaling
Unfolded protein response
Natural Killer Cell Signaling
IL−2 Signaling
Role of NANOG in Mammalian Embryonic Stem Cell Pluripotency
Oncostatin M Signaling
EGF Signaling
FLT3 Signaling in Hematopoietic Progenitor Cells
Antiproliferative Role of Somatostatin Receptor 2
PEDF Signaling
Role of MAPK Signaling in Promoting the Pathogenesis of Influenza
Acute Phase Response Signaling
Macropinocytosis Signaling
Non−Small Cell Lung Cancer Signaling
Glioma Signaling
GM−CSF Signaling
Chemokine Signaling
G..q Signaling
STAT3 Pathway
UVA−Induced MAPK Signaling
Synaptic Long Term Potentiation
CNTF Signaling
Cancer Drug Resistance By Drug Efflux
TNFR1 Signaling
Ceramide Signaling
Acute Myeloid Leukemia Signaling
Endocannabinoid Developing Neuron Pathway
Agrin Interactions at Neuromuscular Junction
Glioblastoma Multiforme Signaling
Neurotrophin/TRK Signaling
14−3−3−mediated Signaling
Thrombopoietin Signaling
MSP−RON Signaling In Cancer Cells Pathway
RANK Signaling in Osteoclasts
Pyridoxal 5'−phosphate Salvage Pathway
Salvage Pathways of Pyrimidine Ribonucleotides
ERBB4 Signaling
VEGF Family Ligand−Receptor Interactions
NF−..B Activation by Viruses
Fc Epsilon RI Signaling
LPS−stimulated MAPK Signaling
Pancreatic Adenocarcinoma Signaling
Renal Cell Carcinoma Signaling
Melanocyte Development and Pigmentation Signaling
IL−3 Signaling
G..12/13 Signaling
IL−6 Signaling
HIF1.. Signaling
fMLP Signaling in Neutrophils
RAC Signaling
UVC−Induced MAPK Signaling
BMP signaling pathway
ERBB Signaling
GDNF Family Ligand−Receptor Interactions
Paxillin Signaling
HGF Signaling
PAK Signaling
P2Y Purigenic Receptor Signaling Pathway
PDGF Signaling
Colorectal Cancer Metastasis Signaling
GNRH Signaling
Tumor Microenvironment Pathway
Cholecystokinin/Gastrin−mediated Signaling
Apelin Endothelial Signaling Pathway
Role of NFAT in Cardiac Hypertrophy
Senescence Pathway
NGF Signaling
Adrenomedullin signaling pathway
Cardiac Hypertrophy Signaling
NRF2−mediated Oxidative Stress Response
Autophagy
Human Embryonic Stem Cell Pluripotency
D−myo−inositol (1,4,5,6)−Tetrakisphosphate Biosynthesis
D−myo−inositol (3,4,5,6)−tetrakisphosphate Biosynthesis
D−myo−inositol−5−phosphate Metabolism
CXCR4 Signaling
Renin−Angiotensin Signaling
HMGB1 Signaling
IL−8 Signaling
3−phosphoinositide Biosynthesis
3−phosphoinositide Degradation
Hepatic Fibrosis Signaling Pathway
Role Of Osteoclasts In Rheumatoid Arthritis Signaling Pathway
Oxytocin Signaling Pathway
Regulation Of The Epithelial Mesenchymal Transition By Growth Factors Pathway
Wound Healing Signaling Pathway
Superpathway of Inositol Phosphate Compounds
Pulmonary Fibrosis Idiopathic Signaling Pathway
z-score
5
0
-5
anti−IL17 NR: Naive CD4+ T3
anti−IL17 R: Naive CD4+ T3
anti−IL17 R: Activated CD8+ T2
anti−IL17 NR: Naive CD4+ T5
anti−IL17 R: Naive CD4+ T 5
anti−IL17 NR: Activated CD8+ T1
anti−IL17 R: Activated CD8+ T1
anti−TNF NR: Activated CD8+ T1
anti−TNF R: Activated CD8+ T1
anti−TNF NR: Activated CD4+ T1
anti−TNF R: Activated CD4+ T1
anti−TNF NR: Activated CD4+ T2
anti−TNF R: Activated CD4+ T2

**Fig. S16. Ingenuity pathway analysis of cell types enriching IL17 or TNF KEGG pathways.** To investigate why scDrugPrio does not prioritize valid targets based on PBMC data from PsA patients, we checked whether treatment-relevant pathways were enriched among the DEGs of responders and nonresponders to the respective drug. We found that only a few cell types were enriched in the TNF-α and IL17 KEGG pathways. Since KEGG does not enable prediction of up- or downregulation, we redid pathway enrichment for the relevant cell types in ingenuity pathway analysis (IPA). In this heatmap, we present IPA-derived z-scores for up- (positive) or down (negative) pathway regulation. The IL17A signalling pathway and TNFR signalling pathway were either downregulated or insignificantly enriched, with a z-score trending towards downregulation.

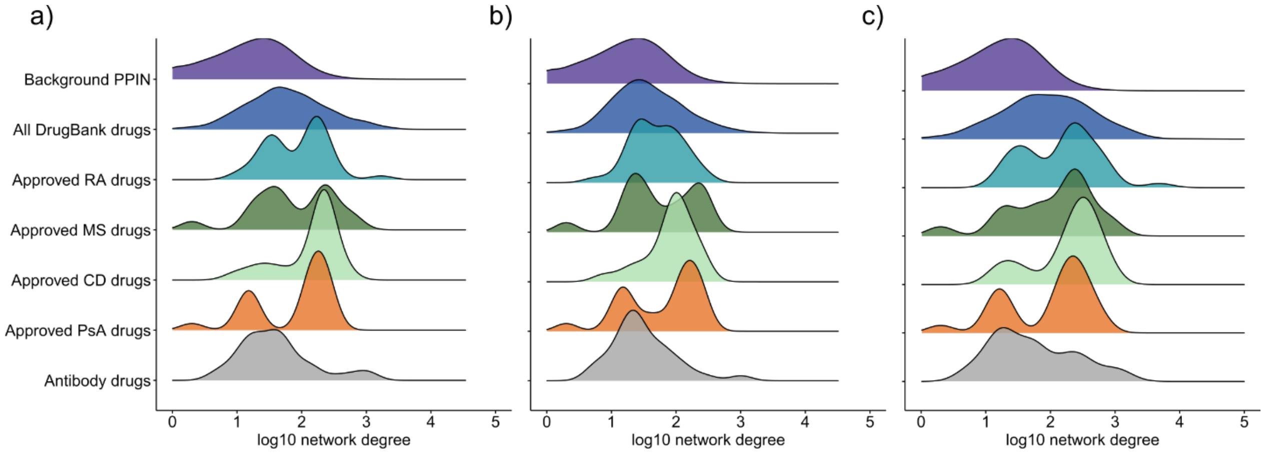


**Fig. S17 Distribution of network degrees.** In this figure, we visualize the distributions for **a)** max, **b)** mean and **c)** sum of drug targets' network degrees compared to the distribution of network degrees in the LCC of the literature-derived PPIN. As updated treatment regimens for inflammatory diseases often have antibody-based drugs as cornerstones, we visualised their network degrees separately. A total of 316 unique antibody drugs were identified by screening DrugBank for drug names ending with -ab.

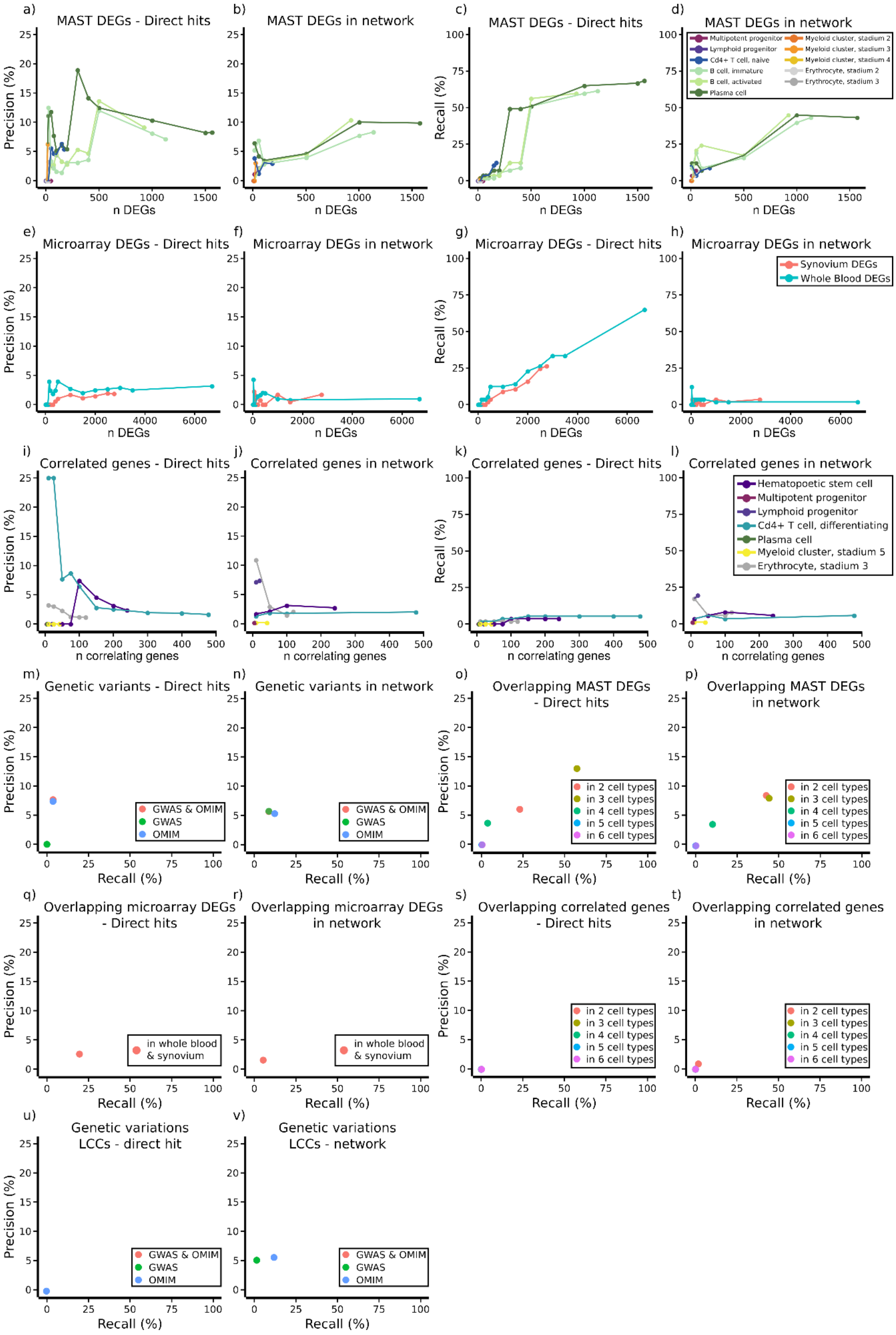
a)
MAST DEGs - Direct hits
Precision (%)
n DEGs
b)
MAST DEGs in network
n DEGs
c)
MAST DEGs - Direct hits
Recall (%)
n DEGs
d)
MAST DEGs in network
Multipotent progenitor
Lymphoid progenitor
Cd4+ T cell, naive
B cell, immature
B cell, activated
Plasma cell
Myeloid cluster, stadium 2
Myeloid cluster, stadium 3
Myeloid cluster, stadium 4
Erythrocyte, stadium 2
Erythrocyte, stadium 3
n DEGs
e)
Microarray DEGs - Direct hits
Precision (%)
n DEGs
f)
Microarray DEGs in network
n DEGs
g)
Microarray DEGs - Direct hits
Recall (%)
n DEGs
h)
Microarray DEGs in network
Synovium DEGs
Whole Blood DEGs
n DEGs
i)
Correlated genes - Direct hits
Precision (%)
n correlating genes
j)
Correlated genes in network
n correlating genes
k)
Correlated genes - Direct hits
Recall (%)
n correlating genes
l)
Correlated genes in network
Hematopoetic stem cell
Multipotent progenitor
Lymphoid progenitor
Cd4+ T cell, differentiating
Plasma cell
Myeloid cluster, stadium 5
Erythrocyte, stadium 3
n correlating genes
m)
Genetic variants - Direct hits
Precision (%)
GWAS & OMIM
GWAS
OMIM
Recall (%)
n)
Genetic variants in network
GWAS & OMIM
GWAS
OMIM
Recall (%)
o)
Overlapping MAST DEGs
- Direct hits
Precision (%)
in 2 cell types
in 3 cell types
in 4 cell types
in 5 cell types
in 6 cell types
Recall (%)
p)
Overlapping MAST DEGs
in network
in 2 cell types
in 3 cell types
in 4 cell types
in 5 cell types
in 6 cell types
Recall (%)
q)
Overlapping microarray DEGs
- Direct hits
Precision (%)
in whole blood
& synovium
Recall (%)
r)
Overlapping microarray DEGs
in network
in whole blood
& synovium
Recall (%)
s)
Overlapping correlated genes
- Direct hits
Precision (%)
in 2 cell types
in 3 cell types
in 4 cell types
in 5 cell types
in 6 cell types
Recall (%)
t)
Overlapping correlated genes
in network
in 2 cell types
in 3 cell types
in 4 cell types
in 5 cell types
in 6 cell types
Recall (%)
u)
Genetic variations
LCCs - direct hit
Precision (%)
GWAS & OMIM
GWAS
OMIM
Recall (%)
v)
Genetic variations
LCCs - network
GWAS & OMIM
GWAS
OMIM
Recall (%)

**Fig. S18 Precision and recall as a function of different feature selection methods and different data sets. a-d)** Precision (a, b) and recall (c, d) derived based on the top x most significant MAST DEGs. Selected genes were checked for known drug targets, and drugs that targeted at least one DEG ("Direct hits") were selected as candidates for calculation of precision (a) and recall (c). Selected genes were also used as input to the previously described network-proximity screening, and drugs that showed $z_c < -1.64$ in the literature curated PPIN were selected as candidates for calculation of precision (b) and recall (d). **e-h)** correspond to a-b) for microarray-derived DEGs. **i-l)** corresponds to a-b) for genes derived by correlation of each cell type's gene expression values with the arthritis score of the AIA mice. **m&n)** shows precision/recall plots for GWAS genes and OMIM genes (as previously defined) as well as a combined gene list. **o&p)** Precision/recall plot investigating the predictive capability of MAST DEGs that overlapped between x cell types. **q&r)** corresponding to o&p) for overlapping microarray-derived DEGs. **s&t)** corresponding to o&p) for cell type-specific genes correlated with arthritis score. **u&v)** Precision/recall plot for the LCCs formed by genetic variations.

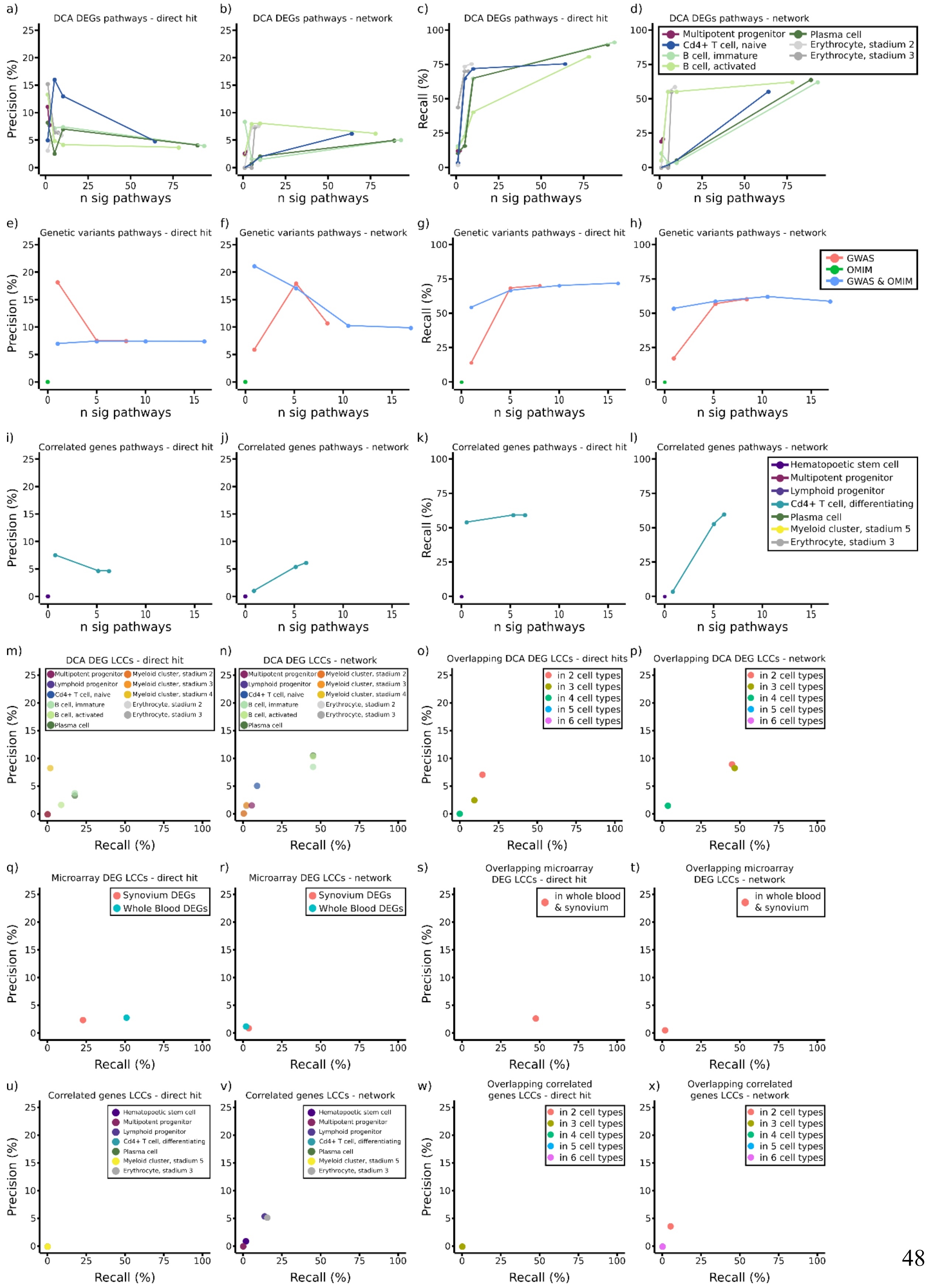
a) DCA DEGs pathways - direct hit
b) DCA DEGs pathways - network
c) DCA DEGs pathways - direct hit
d) DCA DEGs pathways - network
Multipotent progenitor
Cd4+ T cell, naive
B cell, immature
B cell, activated
Plasma cell
Erythrocyte, stadium 2
Erythrocyte, stadium 3
Precision (%)
Recall (%)
n sig pathways
e) Genetic variants pathways - direct hit
f) Genetic variants pathways - network
g) Genetic variants pathways - direct hit
h) Genetic variants pathways - network
GWAS
OMIM
GWAS & OMIM
i) Correlated genes pathways - direct hit
j) Correlated genes pathways - network
k) Correlated genes pathways - direct hit
l) Correlated genes pathways - network
Hematopoetic stem cell
Multipotent progenitor
Lymphoid progenitor
Cd4+ T cell, differentiating
Plasma cell
Myeloid cluster, stadium 5
Erythrocyte, stadium 3
m) DCA DEG LCCs - direct hit
n) DCA DEG LCCs - network
Multipotent progenitor
Lymphoid progenitor
Cd4+ T cell, naive
B cell, immature
B cell, activated
Plasma cell
Myeloid cluster, stadium 2
Myeloid cluster, stadium 3
Myeloid cluster, stadium 4
Erythrocyte, stadium 2
Erythrocyte, stadium 3
o) Overlapping DCA DEG LCCs - direct hits
p) Overlapping DCA DEG LCCs - network
in 2 cell types
in 3 cell types
in 4 cell types
in 5 cell types
in 6 cell types
q) Microarray DEG LCCs - direct hit
r) Microarray DEG LCCs - network
Synovium DEGs
Whole Blood DEGs
s) Overlapping microarray DEG LCCs - direct hit
t) Overlapping microarray DEG LCCs - network
in whole blood & synovium
u) Correlated genes LCCs - direct hit
v) Correlated genes LCCs - network
Hematopoetic stem cell
Multipotent progenitor
Lymphoid progenitor
Cd4+ T cell, differentiating
Plasma cell
Myeloid cluster, stadium 5
Erythrocyte, stadium 3
w) Overlapping correlated genes LCCs - direct hit
x) Overlapping correlated genes LCCs - network
in 2 cell types
in 3 cell types
in 4 cell types
in 5 cell types
in 6 cell types
Precision (%)
Recall (%)

**Fig S19. Precision and recall as a function of different feature selection methods and different data sets.** Continuation of **Fig. S8**. **a-d)** Precision (a, b) and recall (c, d) derived based on the top significantly single-cell derived DEG-enriched KEGG pathways. Selected genes were checked for known drug targets, and drugs that targeted at least one DEG ("Direct hits") were selected as candidates for calculation of precision (a) and recall (c). Selected genes were also used as input to the previously described network-proximity screening, and drugs that showed $z_c < -1.64$ in the literature curated PPIN were selected as candidates for calculation of precision (b) and recall (d). **e-h)** corresponds to a-b) for genetic variations. **i-l)** corresponds to a-b) for genes derived by correlation of each cell type's gene expression values with the arthritis score of the AIA mice. **m&n)** Precision/recall plots for LCCs formed by single-cell derived DEGs. **o&p)** Precision/recall plot for the overlap between LCCs formed by single-cell derived cell type-specific DEGs. **q&r)** corresponding to m&n) for microarray-derived DEGs. **s&t)** corresponding to o&p) for LCCs formed by microarray-derived DEGs. **u&v)** corresponding to m&n) for LCCs formed by cell type-specific genes that correlated with arthritis severity score. **w&x)** corresponding to o&p) for genes overlapping between LCCs formed by correlated genes.

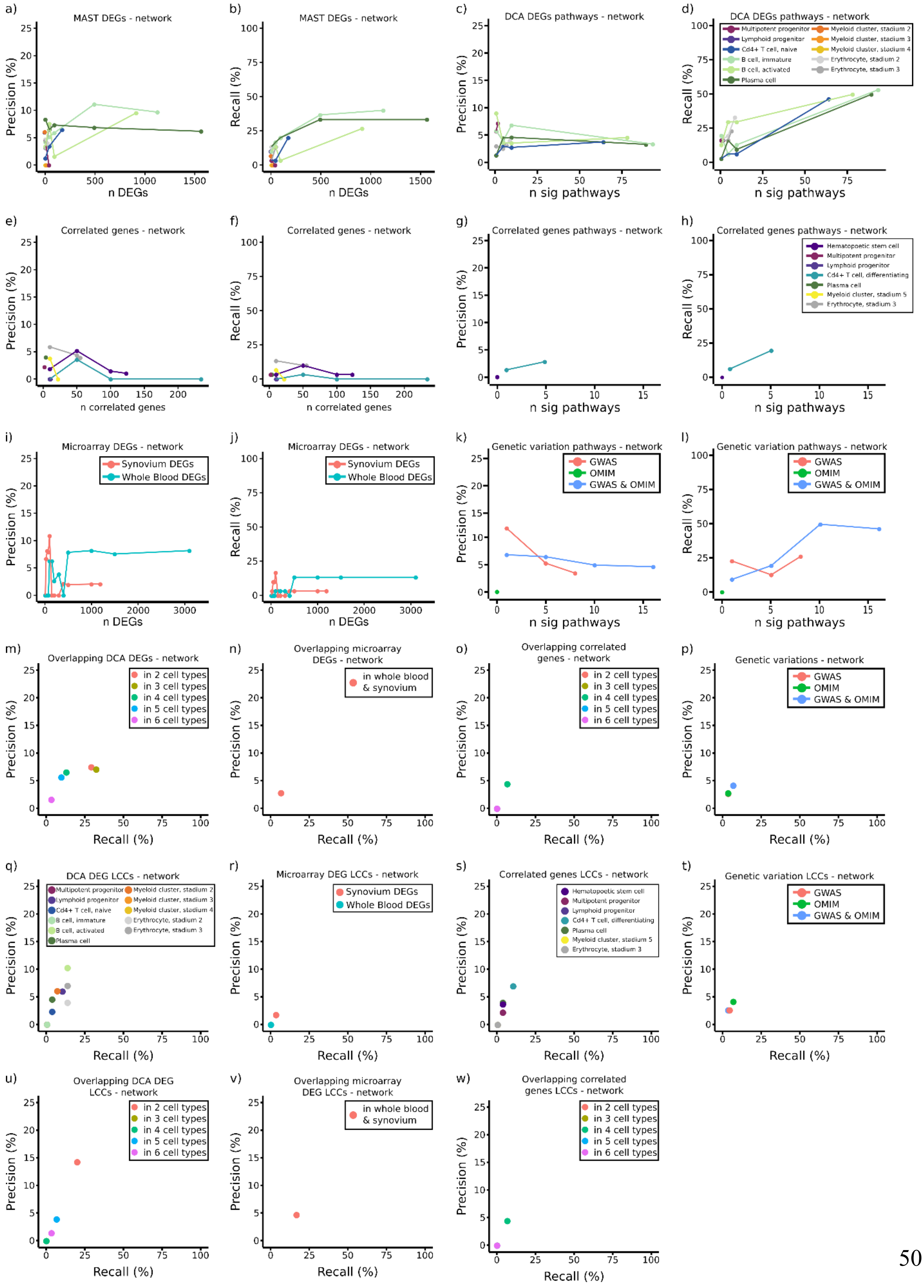
a)
MAST DEGs - network
Precision (%)
n DEGs
b)
MAST DEGs - network
Recall (%)
n DEGs
c)
DCA DEGs pathways - network
Precision (%)
n sig pathways
d)
DCA DEGs pathways - network
Multipotent progenitor
Lymphoid progenitor
Cd4+ T cell, naive
B cell, immature
B cell, activated
Plasma cell
Myeloid cluster, stadium 2
Myeloid cluster, stadium 3
Myeloid cluster, stadium 4
Erythrocyte, stadium 2
Erythrocyte, stadium 3
Recall (%)
n sig pathways
e)
Correlated genes - network
Precision (%)
n correlated genes
f)
Correlated genes - network
Recall (%)
n correlated genes
g)
Correlated genes pathways - network
Precision (%)
n sig pathways
h)
Correlated genes pathways - network
Hematopoetic stem cell
Multipotent progenitor
Lymphoid progenitor
Cd4+ T cell, differentiating
Plasma cell
Myeloid cluster, stadium 5
Erythrocyte, stadium 3
Recall (%)
n sig pathways
i)
Microarray DEGs - network
Synovium DEGs
Whole Blood DEGs
Precision (%)
n DEGs
j)
Microarray DEGs - network
Synovium DEGs
Whole Blood DEGs
Recall (%)
n DEGs
k)
Genetic variation pathways - network
GWAS
OMIM
GWAS & OMIM
Precision (%)
n sig pathways
l)
Genetic variation pathways - network
GWAS
OMIM
GWAS & OMIM
Recall (%)
n sig pathways
m)
Overlapping DCA DEGs - network
in 2 cell types
in 3 cell types
in 4 cell types
in 5 cell types
in 6 cell types
Precision (%)
Recall (%)
n)
Overlapping microarray DEGs - network
in whole blood & synovium
Precision (%)
Recall (%)
o)
Overlapping correlated genes - network
in 2 cell types
in 3 cell types
in 4 cell types
in 5 cell types
in 6 cell types
Precision (%)
Recall (%)
p)
Genetic variations - network
GWAS
OMIM
GWAS & OMIM
Precision (%)
Recall (%)
q)
DCA DEG LCCs - network
Multipotent progenitor
Lymphoid progenitor
Cd4+ T cell, naive
B cell, immature
B cell, activated
Plasma cell
Myeloid cluster, stadium 2
Myeloid cluster, stadium 3
Myeloid cluster, stadium 4
Erythrocyte, stadium 2
Erythrocyte, stadium 3
Precision (%)
Recall (%)
r)
Microarray DEG LCCs - network
Synovium DEGs
Whole Blood DEGs
Precision (%)
Recall (%)
s)
Correlated genes LCCs - network
Hematopoetic stem cell
Multipotent progenitor
Lymphoid progenitor
Cd4+ T cell, differentiating
Plasma cell
Myeloid cluster, stadium 5
Erythrocyte, stadium 3
Precision (%)
Recall (%)
t)
Genetic variation LCCs - network
GWAS
OMIM
GWAS & OMIM
Precision (%)
Recall (%)
u)
Overlapping DCA DEG LCCs - network
in 2 cell types
in 3 cell types
in 4 cell types
in 5 cell types
in 6 cell types
Precision (%)
Recall (%)
v)
Overlapping microarray DEG LCCs - network
in whole blood & synovium
Precision (%)
Recall (%)
w)
Overlapping correlated genes LCCs - network
in 2 cell types
in 3 cell types
in 4 cell types
in 5 cell types
in 6 cell types
Precision (%)
Recall (%)

**Fig S20. Precision and recall in the HuRI protein–protein interaction network (PPIN) as a function of different feature selection methods and different data sets.** For calculation of the above plots, genes that were not found in the HuRI PPIN were removed prior to calculation. This included even the removal of drugs from the calculation if they did not target any gene included in HuRI. This panel exclusively shows the network-proximity screening outcomes in the HuRI PPIN, and drugs that showed $z_c < -1.64$ were selected as candidates for calculation of precision (b) and recall (d). **a&b)** Precision and recall for the top x most significantly single-cell derived DEGs. **c&d)** Precision and recall for DCA DEG-enriched KEGG pathways. **e&f)** corresponding to a&b) for correlated genes. **g&h)** corresponding to c&d) with correlated gene enriched KEGG pathways. **i&j)** corresponding to **a&b)** for microarray-derived DEGs. **k&l)** corresponding to c&d) for genetic variation-enriched KEGG pathways. **m-o)** Precision/recall plot for overlapping single-cell derived DEGs, microarray DEGs and correlated genes, respectively. **p)** Precision/recall plot for genetic variations. **q-t)** Precision/recall plots for LCCs formed by single-cell derived DEGs, microarray DEGs, genes correlated with arthritis score, and genetic variations. **u-w)** Precision/recall plots for LCCs overlapping between cell type-specific single-cell derived DEGs, microarray data sets, and cell type-specific genes correlated with arthritis severity score.

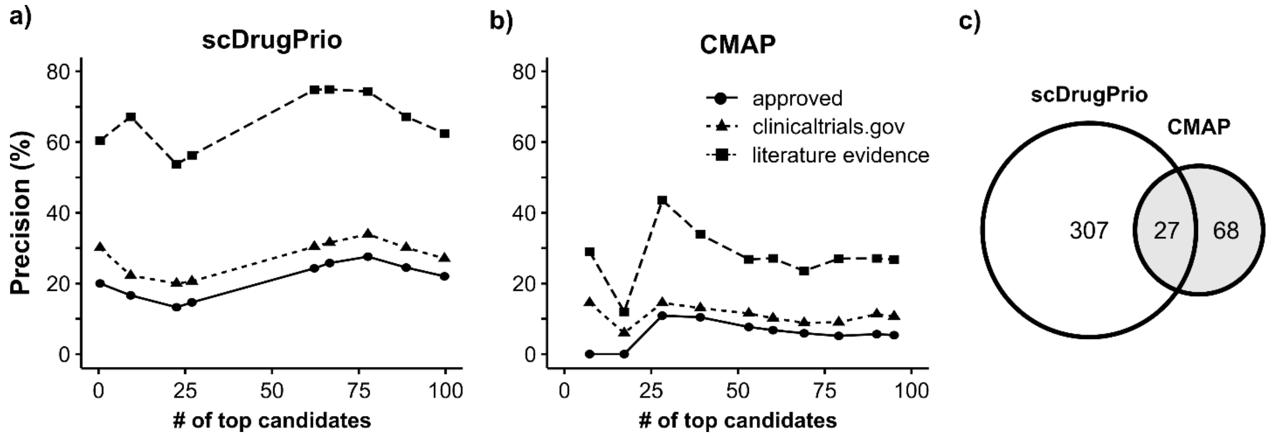


**Fig. S21. Comparison of drug prediction using scDrugPrio and CMAP.** Precision for **a)** the top 100 ranked candidates using scDrugPrio on DEGs derived from scRNA-seq AIA data. In a similar manner, **b)** CMAP prediction was conducted based on DEGs derived from pseudobulk RNA-seq of the AIA data. CMAP prediction was ranked by the CMAP effect measure. Precision is calculated for approved RA drugs as well as for drugs with literature evidence among ranked drug candidates. Both drug predictions used the same DrugBank-derived drug target data for n = 1,840 drugs. **c)** shows the overlap between drug candidates.

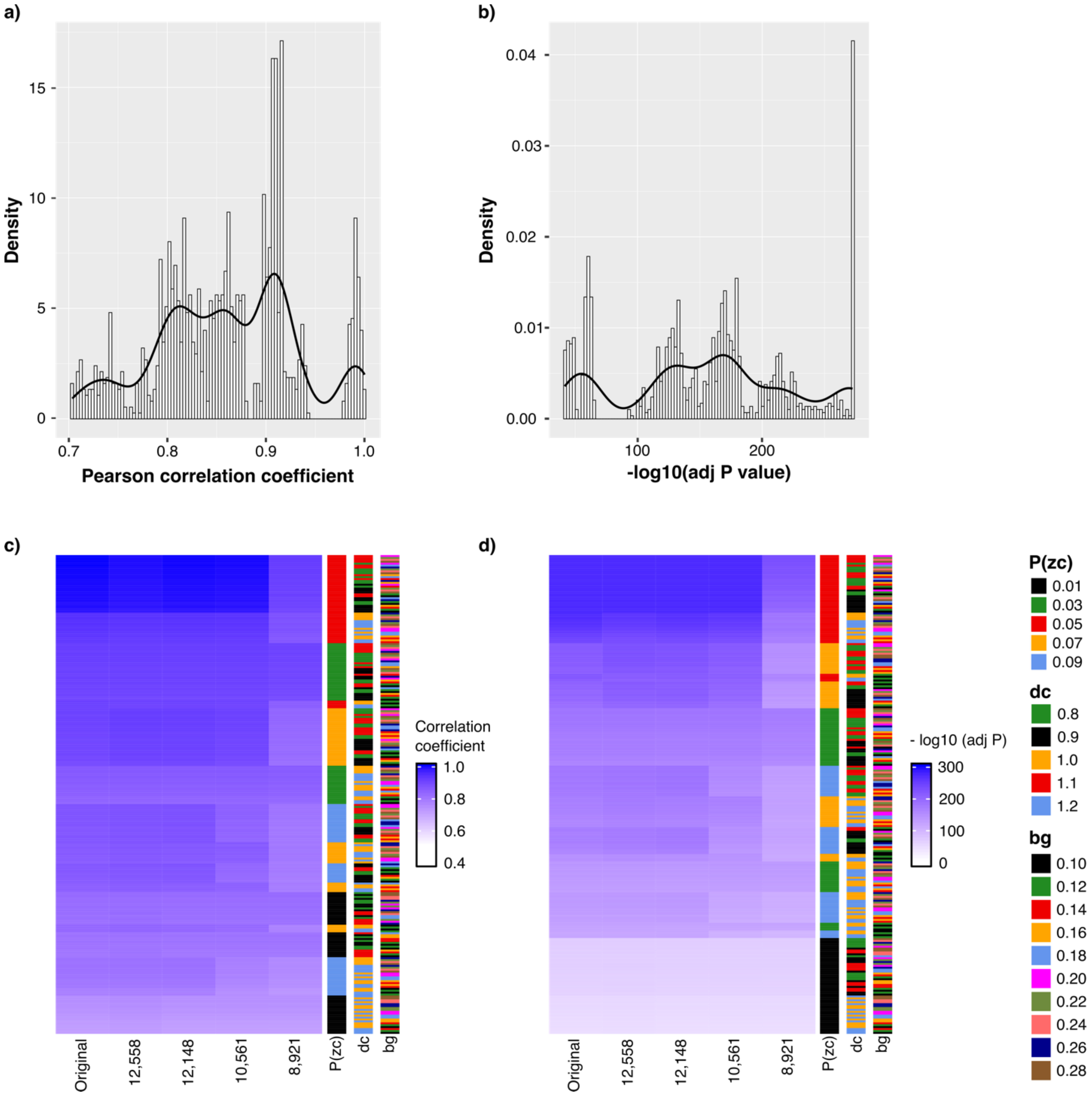


**Fig. S22. Robustness analysis of scDrugPrio.** Robustness was validated by variation of thresholds during recomputation of drug selection and drug ranking for the AIA data set. In this analysis, we varied 1) the number of DEGs that entered network calculations, 2) the P value cut-off for network proximity, derived from $z_c$, 3) network distance cut-off $d_c$, and 4) the cut-off for background gene calculation during NicheNet ligand activity analysis. Standard cut-offs for these included 1) all DEGs (n = 12,769), 2) $P < 0.05$, 3) $dc < 1$, and 4) $Ea(i) \geq 0.2$. Collectively, 1250 drug rankings with varying thresholds were calculated, and the results were compared to the original drug ranking by Pearson correlation of drug ranks between computations. **a)** shows the distribution of Pearson correlation coefficients, **b)** shows the distribution of Bonferroni adjusted P values. Similarly, **c & d)** visualize the Pearson correlation coefficient and adjusted P value, respectively. Side bars indicate the combination of cut-offs used for results in each row. scDrugPrio returns overall returns stable and comparable results over a variety of thresholds.
